# Extremely flexible infection programs in a fungal plant pathogen

**DOI:** 10.1101/229997

**Authors:** Janine Haueisen, Mareike Möller, Christoph J. Eschenbrenner, Jonathan Grandaubert, Heike Seybold, Holger Adamiak, Eva H. Stukenbrock

## Abstract

Filamentous plant pathogens exhibit extraordinary levels of genomic variability that is proposed to facilitate rapid adaptation to changing host environments. However, the impact of genomic variation on phenotypic differentiation in pathogen populations is largely unknown. Here, we address the extent of variability in infection phenotypes of the hemibiotrophic wheat pathogen *Zymoseptoria tritici* by studying three field isolates collected in Denmark, Iran, and the Netherlands. These three isolates differ extensively in genome structure and gene content, but produce similar disease symptoms in the same susceptible wheat cultivar. Using advanced confocal microscopy, staining of reactive oxygen species, and comparative analyses of infection stage-specific RNA-seq data, we demonstrate considerable variation in the temporal and spatial course of infection of the three isolates. Based on microscopic observation, we determined four core infection stages: establishment, biotrophic growth, lifestyle transition, and necrotrophic growth and asexual reproduction. Comparative analyses of the fungal transcriptomes, sequenced for every infection stage, revealed that the gene expression profiles of the isolates differed significantly, and 20% of the genes are differentially expressed between the three isolates during infection. The genes exhibiting isolate-specific expression patterns are enriched in genes encoding effector candidates that are small, secreted, cysteine-rich proteins and putative virulence determinants. Moreover, the differentially expressed genes were located significantly closer to transposable elements, which are enriched for the heterochromatin-associated histone marks H3K9me3 and H3K27me3 on the accessory chromosomes. This observation indicates that transposable elements and epigenetic regulation contribute to the infection-associated transcriptional variation between the isolates. Our findings illustrate how high genetic diversity in a pathogen population can result in highly differentiated infection and expression phenotypes that can support rapid adaptation in changing environments. Furthermore, our study reveals an exceptionally high extent of plasticity in the infection program of an important wheat pathogen and shows a substantial redundancy in infection-related gene expression.

**Author summary:** *Zymoseptoria tritici* is a pathogen that infects wheat and induces necrosis in leaf tissue. *Z. tritici* field populations exhibit high levels of genetic diversity, and here we addressed the consequences of this diversity on infection phenotypes. We conducted a detailed comparison of the infection processes of three *Z. tritici* isolates collected in Denmark, the Netherlands, and Iran. We inoculated leaves of a susceptible wheat cultivar and monitored development of disease symptoms and infection structures in leaf tissue by confocal microscopy. The three isolates exhibited highly differentiated spatial and temporal patterns of infection, although quantitative disease was similar. Furthermore, more than 20% of the genes were differentially expressed in the three isolates during wheat infection. Variation in gene expression is particularly associated with transposable elements, suggesting a role of epigenetic regulation in transcriptional variation among the three isolates. Finally, we find that genes encoding putative virulence determinants were enriched among the differentially expressed genes, suggesting that each of the three *Z. tritici* isolates utilizes different strategies to manipulate host defenses. Our results emphasize that phenotypic diversity plays an important role in pathogen populations and should be considered when developing crop protection strategies.

## Introduction

Population genomics and comparative genome analyses have been applied to characterize genetic variation within and between species of pathogens [1]. Studies of eukaryotic pathogens have demonstrated high levels of intraspecies genetic variability, even in species that predominantly propagate by clonal reproduction [2–5]. In sexual species, frequent recombination contributes to the formation of new genotypes, while transposable elements and repeat-rich genome compartments facilitate the generation of novel genetic variants, most notably in asexual species [6–8]. In addition, many fungal plant pathogens—both sexual and asexual species—carry exceptionally high levels of karyotypic variability that originates from structural chromosome rearrangements and the presence of accessory chromosomes or genome compartments composed of transposable elements [4,5,9–11].

Genetic variation can translate into phenotypic variation that is important for populations to persist in changing environments. High levels of phenotypic variation can provide an adaptive advantage to pathogens exposed to new host resistances or, in agricultural systems, drug treatments. However, while theoretical and empirical data support the importance of genetic variation in rapid adaptation, we still know little about the overall extent and consequences of phenotypic variation in populations of pathogens.

Phenotypic variation in pathogen populations has been studied mainly in the context of virulence and drug resistance. Disease phenotypes have been correlated with genetic maps or genome-wide single nucleotide polymorphism (SNP) data to identify variable sites responsible for distinct virulence phenotypes [12–15]. While virulence and drug resistance traits are main determinants of the overall fitness of a pathogen, other traits also influence infection development. For example, individual strains of pathogens may exhibit variation in the spatial, temporal, and physiological exploration of host tissues, as well in reproductive success. While these traits are not directly linked to virulence, they may greatly impact the fitness of individual isolates and evolution at the population scale [16].

In this study, we addressed the extent of variation in infection phenotypes of a fungal plant pathogen characterized by a high level of genomic variability. We used the wheat pathogen *Zymoseptoria tritici* (syn. *Mycosphaerella graminicola)* as a model to investigate how the development of disease symptoms and the transcriptional program induced during infection varies among three field isolates from geographically distinct locations. *Z. tritici* has a hemibiotrophic lifestyle characterized by an initial biotrophic phase, where the fungus feeds on living host cells, followed by necrotrophic growth where the fungus degrades and takes up nutrients from dead host cells. Genomics, transcriptomics, and proteomics studies have been applied to identify virulence determinants of *Z. tritici*. The haploid genome of *Z. tritici* comprises a high number of accessory chromosomes ranging from 400 kb to 1 Mb in size in the reference isolate IPO323 [17,18]. Recent studies provide evidence for the presence of virulence determinants on the accessory chromosomes, however the genes responsible for these effects have so far not been identified [19]. Furthermore, several genome-wide association (GWAS) and quantitative trait loci (QTL) mapping studies have linked a variety of phenotypic traits to genetic variants and candidate genes [20–24], including the avirulence gene *AvrStb6,* which interacts with the wheat resistance gene *Stb6* [25]. *Z. tritici* has served as a prominent model in population genetic studies of crop pathogens, and genetic variation has been assessed on a local (individual lesions) up to a continental scale. The amount of genetic variation in a *Z. tritici* field population is comparable to the variation found on a continental scale, including multiple regional populations [26–28]. Thus, the plants in a single wheat field are infected by *Z. tritici* isolates with wide genotypic diversity.

Here, we investigated how infection of a susceptible host by genetically and morphologically distinct isolates results in similar levels of quantitative virulence. By combining confocal microscopy, disease monitoring, reactive oxygen species (ROS) localization, and transcriptome analyses, we compiled a detailed characterization of infection phenotypes of the three isolates. We hypothesized that high genetic diversity not only increases the evolutionary potential of the pathogen but also results in a variety of host-pathogen interactions that cause a range of different infection phenotypes. Our combined comparative analyses enabled us to characterize infection morphology and gene expression of the three *Z. tritici* isolates, including a core infection program and isolate-specific infection phenotypes. We conclude that variation in infection and expression phenotypes is important for adaptive evolution of pathogens and needs to be considered in the development of disease control strategies.

## Results and Discussion

### The *Z. tritici* isolates Zt05, Zt09, and Zt10 are equally virulent, but disease symptoms develop at different speeds

We compared virulence phenotypes of three *Z. tritici* isolates Zt05 [29], Zt09 (≙ IPO323AChr18, a derivate of the reference strain IPO323 [18] that lost chromosome 18 [30]), and Zt10 [31], previously collected in Denmark, the Netherlands, and Iran, respectively (S1 Table), on leaves of the highly susceptible winter wheat cultivar Obelisk. We evaluated infections 28 days post inoculation (dpi) by estimating the percentage of leaf area affected by necrosis (Fig 1A) and covered with pycnidia, the asexual fruiting bodies (Fig 1B). The production of pycnidia is an essential measure for pathogen fitness and virulence [32]. Although we observed different levels of necrosis (two-sided Mann-Whitney *U* tests, *P* ≤ 0.0048) we found no significant differences in the amount of pycnidia produced by the three isolates (two-sided Mann-Whitney *U* tests, *P* ≥ 0.034) (S1 Fig).

**Figure 1.**
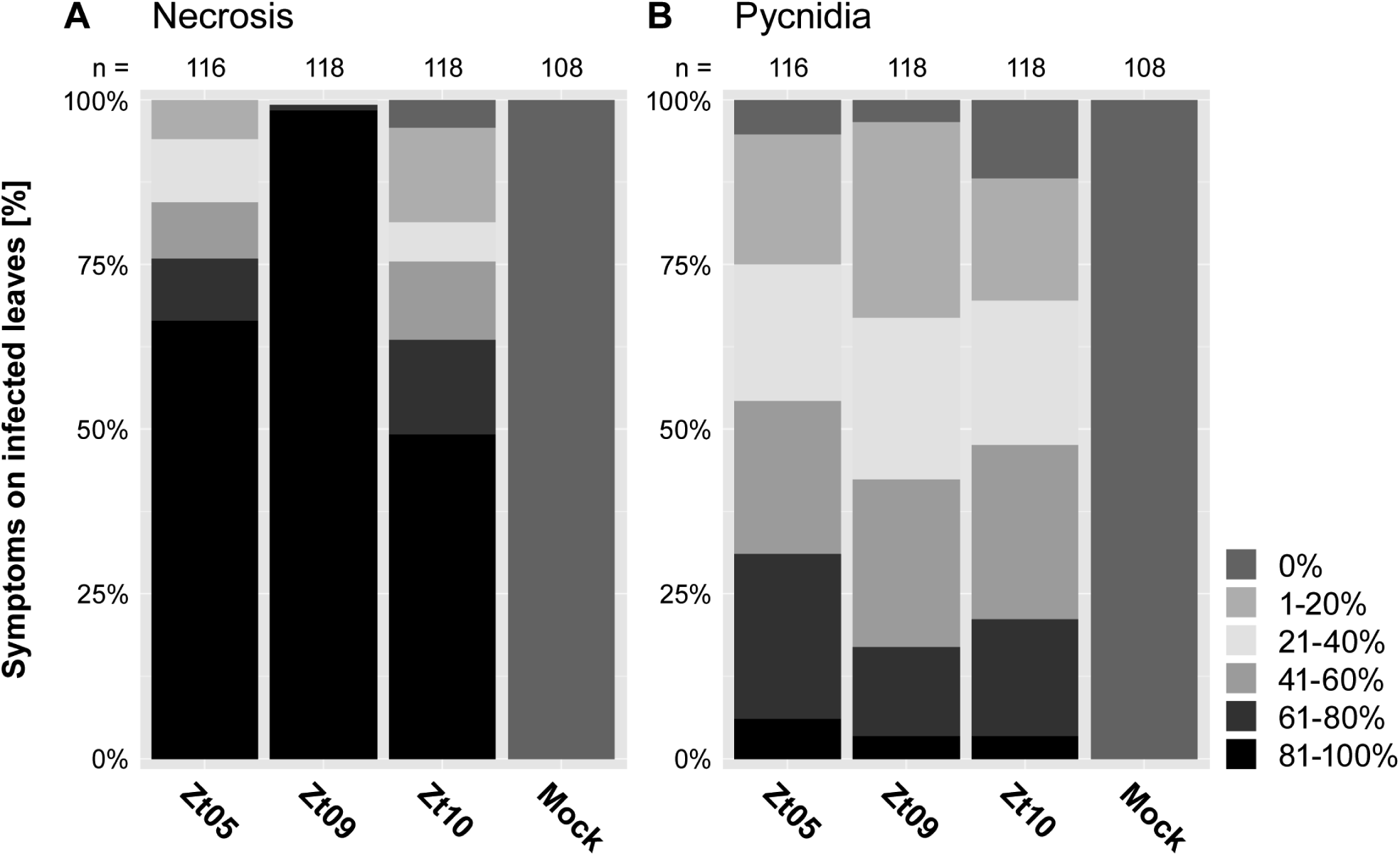
*In-planta* phenotypic assay demonstrates similar pycnidia levels of *Z. tritici* isolates on the susceptible wheat cultivar Obelisk. Quantitative differences in **(A)** necrosis and **(B)** pycnidia coverage of inoculated leaf areas were manually assessed at 28 days post inoculation based on six categories: 0 (without visible symptoms), 1 (1% to 20%), 2 (21% to 40%), 3 (41% to 60%), 4 (61% to 80%), and 5 (81% to 100%). The three isolates caused different levels of necrosis (two-sided Mann-Whitney *U* tests, *P* ≤ 0.0048) but pycnidia levels were not different (two-sided Mann-Whitney *U* tests, *P* ≥ 0.034).

Because the three *Z. tritici* isolates are equally fit and virulent on the wheat cultivar Obelisk, we next investigated whether disease symptoms and pathogen infection develop at a similar pace. We monitored temporal disease progress by screening the leaves every other day for visible necrotic spots and pycnidia. Leaves inoculated with Zt05 and Zt09 showed necrosis and pycnidia significantly earlier than leaves inoculated with Zt10 (one-sided Mann-Whitney *U* tests, *P* ≤ 7.73*10 ^8^) (Fig 2A). The median onset of necrosis caused by Zt09 occurred one day after that caused by Zt05 and is significantly later (one-sided Mann-Whitney *U* test, *P* = 0.0089), although the first pycnidia of both isolates developed at the same time (two-sided Mann-Whitney *U* test, *P* = 0.9455). Thus, although the three *Z. tritici* isolates produced the same pycnidia density in the cultivar Obelisk, disease develops at different paces.

**Figure 2.**
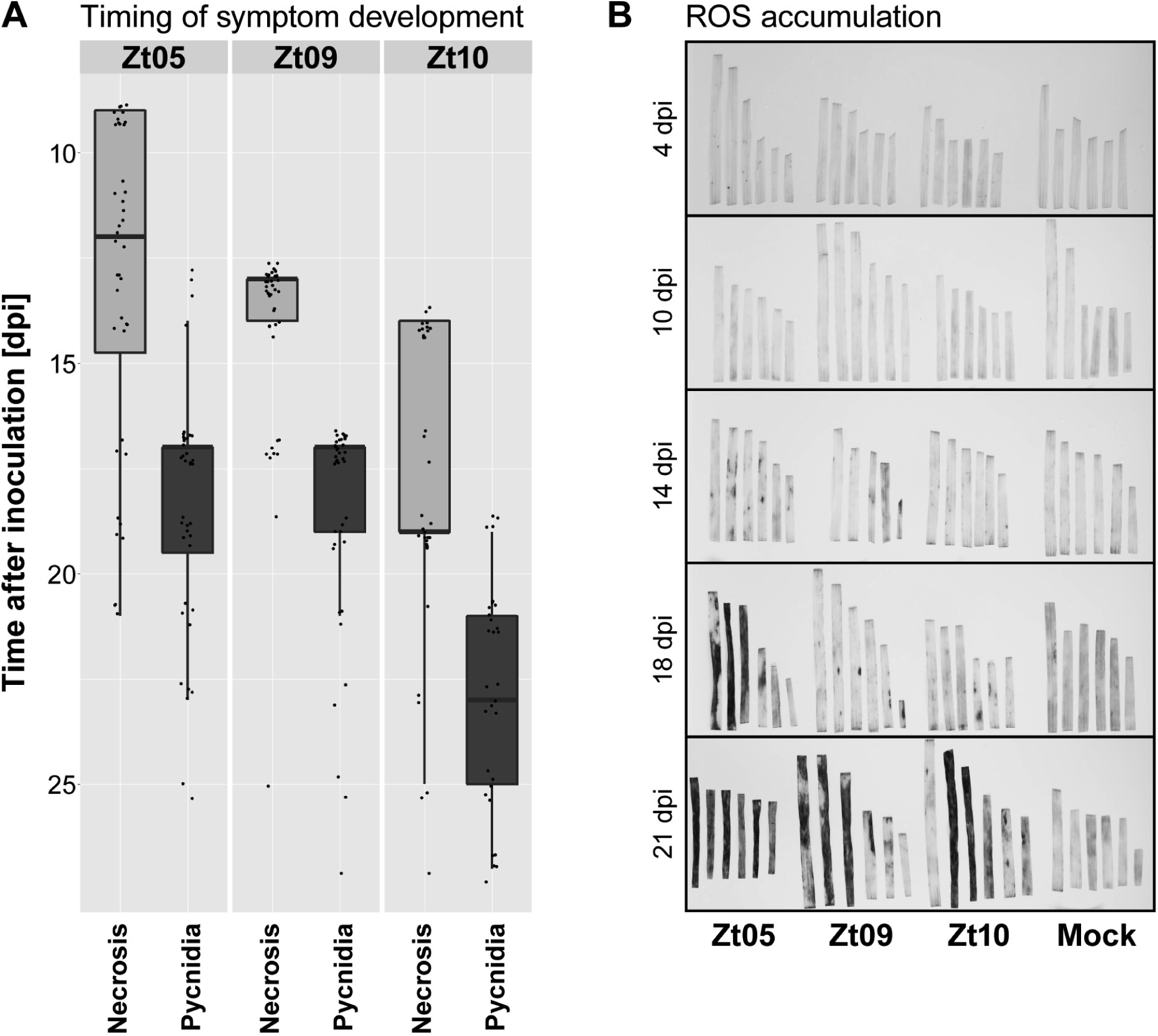
Timing of disease symptom development and H_2_O_2_ accumulation varies between wheat leaves infected with different *Z. tritici* isolates. **(A)** Temporal disease progression for infections with Zt05, Zt09, and Zt10 was measured by daily manual screening for the first occurrence of necrotic spots and pycnidia. For each isolate, 40 leaves of the wheat cultivar Obelisk were inoculated and tested. No disease developed on seven of the leaves inoculated with Zt10. **(B)** Infected leaves were stained for accumulation of the reactive oxygen species H_2_O_2_ at 4, 10, 14, 18, and 21 dpi by 3,3′-diaminobenzidine. Dark red-brown precipitate indicates H_2_O_2_ accumulation and appeared first in leaves infected with Zt05 in leaf areas beginning to undergo necrosis.

ROS play a central role in plant pathogen defense by acting as signalling molecules after pathogen recognition and activating defense responses [33]. We visualized the accumulation of the ROS H_2_O_2_ in infected leaves by diaminobenzidine (DAB) staining to determine whether the observed differences in temporal disease development of *Z. tritici* isolates reflect a temporal variation in host response. Ten to 14 days after inoculation, we observed no ROS (Fig 2B) indicating that *Z. tritici* suppresses the activation of wheat immune responses during this phase. ROS accumulation coincided with the onset of necrosis (Fig 2B, S2 Fig) and, consistent with the faster disease progress, ROS accumulates earlier in leaves infected with Zt05 than Zt09 or Zt10. For Zt09 and Zt10, high ROS concentrations appear at only 18 dpi when Zt05 infected leaves are already largely necrotic and the DAB staining indicates further increased ROS concentrations (Fig 2B, S2 Fig). Thereby, the timing of ROS accumulation in response to the three *Z. tritici* isolates is consistent with the observed differences in the temporal development of disease symptoms in infected wheat tissue.

### *Z. tritici* isolates tolerate different levels of abiotic stress

*Z. tritici* is characterized by a dimorphic lifestyle with hyphal growth during host infection and predominantly yeast-like growth in culture [34]. The yeast/hyphae dimorphism is likely inherited as a multigenic quantitative trait [35] and is essential for pathogenicity [36]. The fungus is exposed to a multitude of environmental influences during infection, dispersal, and other less well characterized stages of the life cycle such as saprotrophic growth and spore dormancy. We compared tolerance of the three *Z. tritici* isolates to several abiotic *in vitro* stressors (temperature, oxidative, osmotic, and cell wall stresses) to assess the variability in growth phenotypes. Colonies of the three isolates exhibited different morphologies and tolerated different levels of abiotic stress (Table 1, S3 Fig). Only osmotic stress led to the same level of reduced growth in all strains. Under all tested conditions, colonies of Zt09 and Zt10 were mainly composed of yeast-like cells, whereas Zt05 predominantly grew as hyphae. On plates with Congo red or calcoflour white, Zt05 formed hyphal colonies, similar to those formed on yeast-malt-sucrose (YMS) control plates, whereas Zt09 and Zt10 were growth-impaired. These results indicate differences in the cell wall composition of yeast-like and hyphal cells, and that yeast-like cells are more susceptible to cell wall-interfering agents.

**Table 1.**
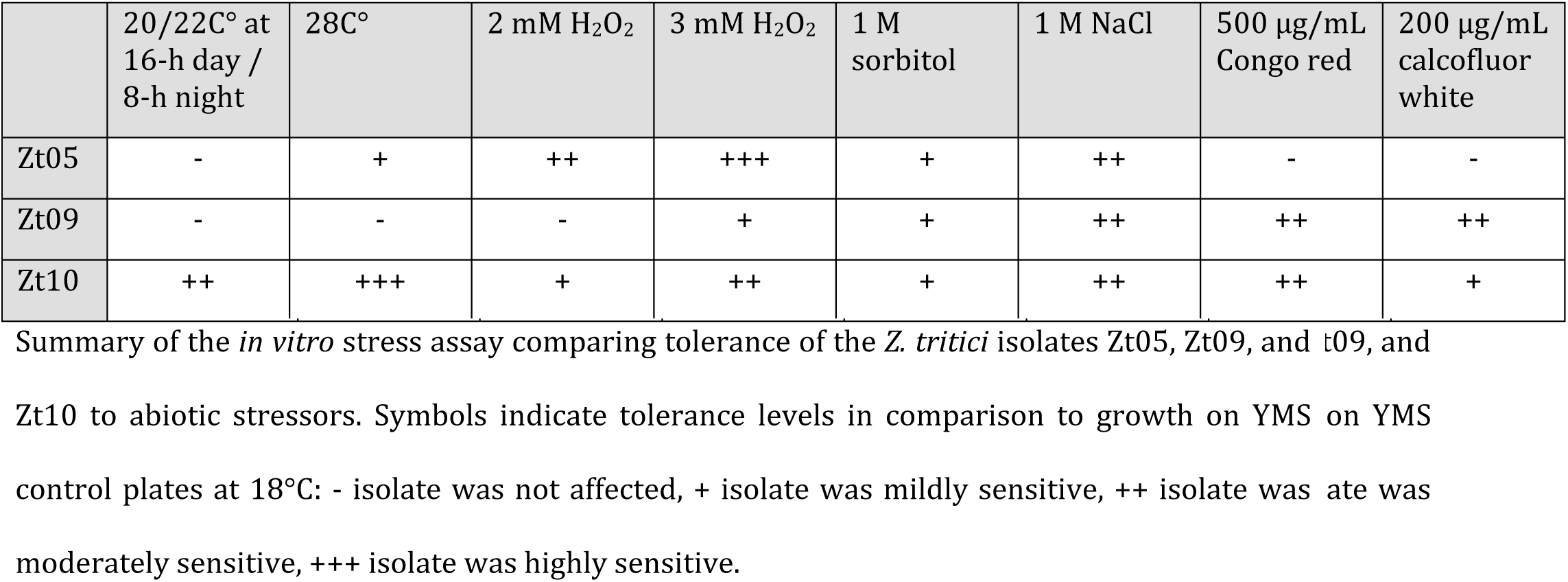
The three *Z. tritici* isolates vary in tolerance to abiotic stressors.

|  | 20/22C° at<br>16-h day /<br>8-h night | 28C° | 2 mM H<sub>2</sub>O<sub>2</sub> | 3 mM H<sub>2</sub>O<sub>2</sub> | 1 M<br>sorbitol | 1 M NaCl | 500 µg/mL<br>Congo red | 200 µg/mL<br>calcofluor<br>white |
| --- | --- | --- | --- | --- | --- | --- | --- | --- |
| Zt05 | - | + | ++ | +++ | + | ++ | - | - |
| Zt09 | - | - | - | + | + | ++ | ++ | ++ |
| Zt10 | ++ | +++ | + | ++ | + | ++ | ++ | + |

Summary of the *in vitro* stress assay comparing tolerance of the *Z. tritici* isolates Zt05, Zt09, and Zt10 to abiotic stressors. Symbols indicate tolerance levels in comparison to growth on YMS control plates at 18°C: - isolate was not affected, + isolate was mildly sensitive, ++ isolate was moderately sensitive, +++ isolate was highly sensitive.

Elevated temperatures greatly impact development of Zt10 that formed strongly melanized colonies at 20/22°C and 28°C. Melanin pigments have several important functions in fungal pathogens, including protection against harsh environmental conditions [37]. Zt10 was collected in the Ilam Province, one of the hottest and driest regions in Iran [38], and the increased melanization could reflect local adaptation to extreme temperatures and temperature fluctuations [39], desiccation, and increased UV radiation.

Oxidative stress eventually diminishes growth of all isolates, although Zt09 tolerated exposure to H2O2 more than Zt05 and Zt10. *Z. tritici* experiences oxidative stress *in planta* from ROS produced as a host defense response or released from dead tissue. In general, higher tolerance to ROS is advantageous [40], for example during necrotrophic growth and pycnidia formation in dead mesophyll tissue [41]. However, mechanisms to detoxify extracellular ROS must be tightly regulated to avoid ROS levels that are toxic to the hosts [33], and we speculate that the observed differences in H_2_O_2_ tolerance reflect divergent adaptation of the *Z. tritici* isolates to host populations with different defense responses to pathogen invasion. Together, the *in vitro* stress assay revealed unanticipated intraspecies variation in tolerance to abiotic stresses among the *Z. tritici* isolates, especially considering that all were isolated in agro-ecosystems from the same host species, *Triticum aestivum.* The variation in colony morphology and stress responses may reflect different adaptations of the *Z. tritici* isolates to their local environments, and our observations suggest that ecological adaptation of fungal plant pathogens can be a strong driver of phenotypic divergence.

### *Z. tritici* infection is characterized by four core developmental stages

Next, we aimed to morphologically characterize host colonization of the three *Z. tritici* isolates. We conducted detailed confocal microscopy analyses in which we scanned 101 leaves harvested at 16 time points after inoculation. Analyses of large z-stacks of longitudinal optical sections allowed us to infer the spatial and temporal fungal colonization on and in infected leaves. First, we focused on the commonalities in host colonization between the different isolates and identified a sequence of four stages that we define as the core infection program of *Z. tritici* (Fig 3).

**Figure 3.**
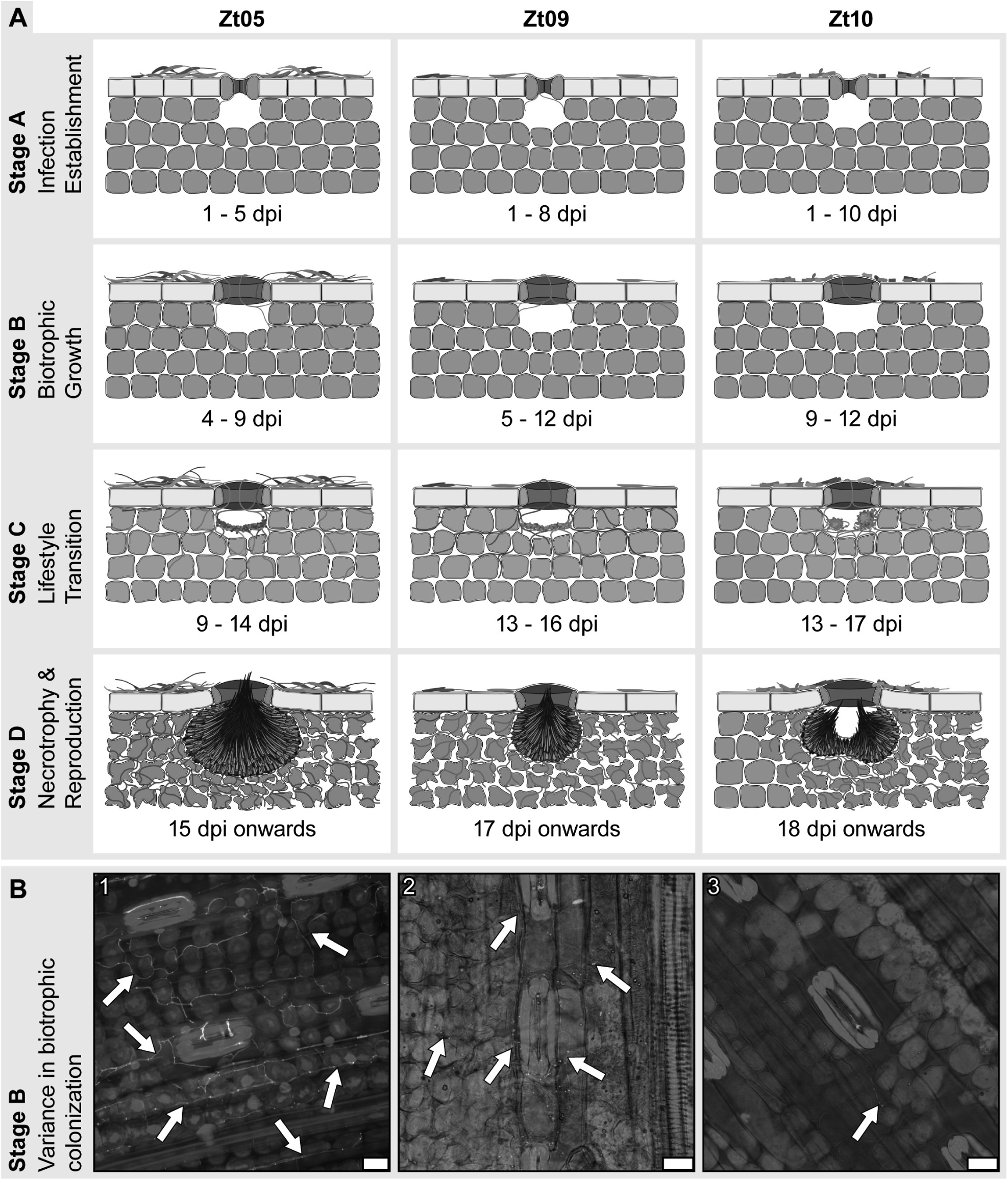
*Z. tritici* wheat infections are characterized by four distinct infection stages and isolate-specific infection development. **(A)** Schematic drawings of the key features that characterize the four infection stages of *Z. tritici* and illustrate the infection phenotypes of isolates Zt05, Zt09, and Zt10 on the wheat cultivar Obelisk. **(B)** Micrographs showing *Z. tritici* hyphae (arrows) during biotrophic growth inside wheat leaves. Maximum projections of confocal image z-stacks. Nuclei and wheat cells are displayed in *purple* and fungal hyphae or septae in *green.* The panel shows biotrophic colonization of **(1)** isolate Zt05 at 7 dpi, **(2)** Zt09 at 11 dpi, and **(3)** Zt10 at 9 dpi. Scale bars = 25 μm.

Stage A, or infection establishment, involves the penetration of wheat leaf tissue by fungal hyphae. Fungal cells germinate after inoculation on the leaf surface, indicating that germination is triggered extrinsically after the fungus senses particular plant-derived cues, as has been previously shown for *Fusarium oxysporum* [42]. Germ tubes develop into infection hyphae, some of which grow in the direction of stomata (Fig 3A stage A, S1 and S2 Animation). During stomatal penetration and in the sub-stomatal cavities, infection hyphae grow in tight contact with the guard cells. We occasionally noticed slight swelling of hyphae on top of stomatal openings resembling primitive appressoria [43,44]. However, we never observed penetration of epidermal cells.

Stage B refers to the symptomless, biotrophic intercellular colonization of the wheat mesophyll (Fig 3A stage B, S3 and S4 Animation). During this stage, the pathogen explores host tissue while avoiding recognition by the host immune system, which otherwise would have resulted in a resistance response in the infected leaf tissue (Fig 2B). Interestingly, hyphae first grow in the interspace of epidermis and mesophyll, where they spread via the grooves between adjacent epidermal cells before deeper mesophyll cell layers are colonized.

Infection stage C comprises the transition from biotrophic to necrotrophic growth when the first disease symptoms develop (Fig 3A, stage C). For asexual reproduction, *Z. tritici* requires large amounts of nutrients that are released from dead host cells. Fungal hyphae branch extensively and colonize all mesophyll layers, with hyphae growing around individual plant cells as they die. Primal structures of pycnidia start to develop (S5-S7 Animation), and ring-like scaffolds form in sub-stomatal cavities where hyphae align and build stromata that give rise to conidiogenous cells. Finally, infection stage D is characterized by necrotrophic colonization and asexual reproduction (Fig 3A, stage D). Hyphae colonize an environment that is nutrient-rich but also hostile due to the high abundance of ROS, as demonstrated by DAB staining (Fig 2B). Within necrotic lesions, mesophyll tissue is heavily colonized, plant cells are dead, and mature pycnidia are visible (S8 and S9 Animation). Hyphae wrap around collapsed host cells, possibly to improve the acquisition of nutrients and protect resources from competing saprotrophs. Sub-stomatal cavities are occupied by sub-globose pycnidia which harbour hyaline, oblong asexual pycnidiospores that extrude through stomatal openings.

While the four infection stages of *Z. tritici* can be clearly distinguished, infections by different cells of one isolate are not completely synchronized; different infection stages can be present simultaneously on the same inoculated leaf.

During biotrophic colonization of wheat tissue, fluorescence emitted from fluorescein isothiocyanate conjugated to wheat germ agglutinin (WGA-FITC) primarily came from septae and was weak compared to that during necrotrophic colonization, during which fluorescence was also emitted from interseptal regions. Previously, similar observations were reported in endophytic and epiphyllous [45] and biotrophic and necrotrophic hyphae [46]. WGA binds to N-acetylglucosamine residues that build chitin, an elicitor of plant immunity [47]. Fungal plant pathogens can prevent recognition, e.g. through chitin-binding LysM effectors [48–50] like the extracellular LysM protein ChELP2, that was also shown to limit accessibility of chitin to WGA in biotrophic hyphae of *C. higginsianum* [46]. In *Z. tritici,* two LysM effectors protect hyphae from plant chitinases [51], and Mg3LysM shields chitin from recognition by wheat receptors [52]. Hence, it is possible that Mg3LysM also limits binding of WGA to chitin during biotrophic, but not necrotrophic, colonization of *Z. tritici* leading to the differences in fluorescence signals from biotrophic and necrotrophic hyphae.

### Highly distinct development of host infection by the three *Z. tritici* isolates

While we clearly recognized the four core infection stages for the isolates Zt05, Zt09, and Zt10, we also observed differences. In stage A, a main difference was the time between inoculation and first stomatal penetration. Infection hyphae of Zt05 enter stomata at 1 to 5 dpi, whereas germ tube formation and stomatal penetration for Zt09 and Zt10 occur later (Fig 3A, stage A). For Zt05, we observed strong epiphyllous proliferation and, in contrast to the two other isolates, frequently the occurrence of several hyphae entering a single stoma (Fig 3A, stage A: Zt05). Stomatal penetrations were evenly distributed for Zt05 and Zt09 within inoculated leaf areas, while Zt10 penetrations were more clustered, leading to patchy infections (Fig 3A, stage C and D: Zt10).

The most striking difference between the isolates was the extent of biotrophic colonization during stage B (Fig 3B). Zt05 develops expanded biotrophic hyphal networks with long “runner” hyphae growing longitudinally between epidermis and mesophyll (Fig 3A, stage B: Zt05, Fig 3B.1, S3 Animation). Zt09 produces fewer hyphae that are located mainly between epidermis and mesophyll and in the first mesophyll cell layer (Fig 3A, stage B: Zt09, Fig 3B.2). Biotrophic growth of Zt10 is limited to the mesophyll cells adjacent to sub-stomatal cavities (Fig 3A, stage B: Zt10, Fig 3B.3). Because biotrophic colonization depends on successful evasion of host immunity [53], the different extent of colonization could reflect different strategies to bypass recognition in a given host genotype.

During stages C and D, the differences in infection development primarily relate to temporal variances. First, Zt05 switches to necrotrophic growth (9 to 14 dpi), followed by Zt09 (13 to 16 dpi) and Zt10 (13 and 17 dpi) (Fig 3A, stage C). The isolates enter stage D in the same order. Furthermore, Zt10 forms two pycnidia in one sub-stomatal cavity (Fig 3A, stages C and D: Zt10, S7 Animation) more often than Zt05 and Zt09. Taken together, the infection development of the studied *Z. tritici* isolates is highly divergent, although the final production of asexual pycnidia does not differ significantly (Fig 1), suggesting that the isolate-specific characteristics in host-pathogen interactions add up to equally successful strategies for colonization and reproduction in a susceptible wheat cultivar. We conclude that infection development of *Z. tritici* can be highly flexible with respect to the timing of the lifestyle transition and the spatial distribution of infecting hyphae inside host tissue.

### Genomes of the three *Z. tritici* isolates exhibit high variation and different chromosome composition

Next, we investigated the genomes of the three isolates. High levels of genomic variability have previously been described within and between species in the genus *Zymoseptoria* to which, in particular, the content and composition of the accessory chromosomes contribute [18,28,54–56]. A previously conducted whole genome comparison using the isolate IPO323 (39.7 Mb) [18] as reference, identified 500,177 single nucleotide polymorphisms (SNPs) in Zt05 and 617,431 SNPs in Zt10, indicating a considerable genetic distance between the three isolates [57]. In order to further quantify genomic variation between the three isolates in our study, we performed electrophoretic karyotyping and whole genome long read-sequencing (S1 Text).

We visualized and compared small chromosomes in the range of 225 to 1,125 kb that are known to exhibit size variation and presence-absence polymorphisms by pulsed-field gel electrophoresis (PFGE). We observed very different karyotypes with no small chromosomes of the same size (S4 Fig). Further, the PFGE results suggest that Zt05 and Zt10 possess at least seven and four putative accessory chromosomes, respectively and show length polymorphisms of the smallest core chromosomes 12 (~1.463 kb) and 13 (~1,186 kb) compared to Zt09, consistent with a previous study [58]. However, the loss of chromosome 18 (~574 kb) in Zt09 in comparison to IPO323 could not be visualized by PFGE as the applied conditions allow no separation from chromosomes 17 (~584 kb) and 16 (~607 kb).

To further assess variation in genomic content and structure, we compared the synteny of Zt05 and Zt10 contigs to the chromosomes of the *Z. tritici* reference IPO323 [18]. To this end, we assembled long-read SMRT Sequencing data for Zt05 and Zt10 and obtained high-quality *de novo* genome assemblies with contig N50 values of 2.45 Mb and 2.93 Mb and assembly sizes of 41.95 Mb and 39.33 Mb, respectively (S2 Table). In total, we obtained 62 unique contigs (unitigs) for Zt05 and 22 for Zt10. We identified telomeric repeats at both ends of 15 unitigs in Zt05 and 17 unitigs in Zt10 and consider these to be completely assembled chromosomes (S2 Table). By whole-chromosome synteny analyses using SyMAP, we identified large syntenic DNA blocks for all 21 chromosomes of IPO323 in the Zt05 assembly, while Zt10 lacked homologs of chromosomes 18, 20 and 21 (S5 and S6 Figs, S2 Table). Although karyotypes and chromosome structure are very different in Zt05 and Zt10 in comparison to the reference IPO323 and hence the derived Zt09 (≙IPO323AChr18), we find homologous regions from all chromosomes and only ~ 2.58 Mb (Zt05) and ~ 2.75 Mb (Zt10) of unique DNA.

To identify the genes that are shared between the three *Z. tritici* isolates, we performed nucleotide BLAST analyses using the coding sequences of the 11,839 annotated genes of the reference IPO323 as input [59]. We identified 11,138 IPO323 genes (94.08%) in Zt05 and 10,745 (90.76%) in Zt10 (e-value cut-off 1e^-3^, identity ≥90%, query coverage between 90% and 110%). The gene presence/absence patterns correlate with the absence of large syntenic DNA blocks of chromosomes 18, 20, and 21 in Zt10 (S6 Fig, S2 Table). 91% of genes on core chromosomes are shared, while only 49% (313 of 643) of the genes located on accessory chromosomes are present in Zt05, Zt09, and Zt10 (S2 Table). Similarly, only 85% (370 of 434) of the previously identified genes encoding candidate secreted effector proteins (CSEPs) [6] were found in all isolates, pointing to high levels of plasticity in the effector repertoire of the three isolates Zt05, Zt09, and Zt10. In total, 10,426 genes were present in Zt05, Zt09, and Zt10 and were considered to be *Z. tritici* core genes for further analyses (S3 Table). In summary, the genome comparison of Zt05, Zt09, and Zt10 shows a high extent of variation at single nucleotide positions as well as structural variation including differences in the total gene content.

### Generation of isolate- and stage-specific transcriptomes based on confocal microscopy analyses

Given the morphological and temporal differences in infection development, we next asked how gene expression profiles differ between the *Z. tritici* isolates Zt05, Zt09, and Zt10 during wheat infection. Previous studies have demonstrated transcriptional re-programming in *Z. tritici* during infection [30,60–64] and shown different transcriptional programs of strains that differ in extent of virulence [65]. Those studies focused primarily on the reference isolate IPO323, and the sequenced material was sampled at defined time points of infection that were not coordinated to distinct infection stages as described above.

We collected leaf material at up to nine time points per isolate and conducted confocal microscopy analyses to select samples for RNA extraction and transcriptome sequencing based on the morphological infection stage (S7 Fig, S4 Table). We generated stage-specific RNA-seq datasets corresponding to the four core infection stages, allowing us to compare the isolate-specific expression profiles at the same developmental stage of infection. Our final dataset comprises four stage-specific transcriptomes per isolate with two biological replicates per sample (Table 2).

**Table 2.** Summary of the stage-specific transcriptomes (A-D) of the three *Z. tritici* isolates

| Isolate | Infection stage | Sample | Time point (dpi) | No. of filtered reads | No. of reads mapped to genome | % reads mapped to genome | No. of genes FPKM $\geq 2^*$ | No. of genes FPKM $\geq 10^*$ |
| --- | --- | --- | --- | --- | --- | --- | --- | --- |
| **Zt05<sup>1</sup>** | A | Zt05-Ta_A_01 | 3 | 107,507,137 | 15,213,307 | 14.15 | 9,302 | 7,455 |
|  |  | Zt05-Ta_A_02 |  | 81,479,903 | 10,509,515 | 12.90 |  |  |
|  | B | Zt05-Ta_B_01 | 8 | 113,732,295 | 15,057,425 | 13.24 | 9,404 | 7,623 |
|  |  | Zt05-Ta_B_02 |  | 128,271,704 | 14,856,314 | 11.58 |  |  |
|  | C | Zt05-Ta_C_01 | 13 | 91,298,814 | 26,756,807 | 29.31 | 9,538 | 7,982 |
|  |  | Zt05-Ta_C_02 |  | 135,198,719 | 34,329,884 | 25.39 |  |  |
|  | D | Zt05-Ta_D_01 | 20 | 86,100,462 | 41,871,582 | 48.63 | 9,585 | 7,914 |
|  |  | Zt05-Ta_D_02 |  | 119,101,106 | 76,123,086 | 63.91 |  |  |
| **Zt09<sup>2</sup>** | A | Zt09-Ta_A_01 | 4 | 129,342,007 | 5,868,572 | 4.54 | 9,435 | 7,482 |
|  |  | Zt09-Ta_A_02 |  | 92,711,865 | 5,582,561 | 6.02 |  |  |
|  | B | Zt09-Ta_B_01 | 11 | 96,767,482 | 5,034,677 | 5.20 | 9,718 | 7,910 |
|  |  | Zt09-Ta_B_02 |  | 103,428,015 | 5,964,566 | 5.77 |  |  |
|  | C | Zt09-Ta_C_01 | 13 | 121,529,652 | 31,264,373 | 25.73 | 9,892 | 8,220 |
|  |  | Zt09-Ta_C_02 |  | 93,253,633 | 27,790,773 | 29.80 |  |  |
|  | D | Zt09-Ta_D_01 | 20 | 110,296,264 | 84,263,562 | 76.40 | 9,867 | 7,949 |
|  |  | Zt09-Ta_D_02 |  | 101,757,635 | 75,044,880 | 73.75 |  |  |
| **Zt10<sup>3</sup>** | A | Zt10-Ta_A_01 | 6 | 93,557,587 | 4,895,650 | 5.23 | 8,814 | 7,219 |
|  |  | Zt10-Ta_A_02 |  | 91,111,840 | 4,836,535 | 5.31 |  |  |
|  | B | Zt10-Ta_B_01 | 11 | 94,828,793 | 5,951,869 | 6.28 | 9,068 | 7,407 |
|  |  | Zt10-Ta_B_02 |  | 110,255,245 | 7,896,227 | 7.16 |  |  |
|  | C | Zt10-Ta_C_01 | 13 | 86,652,710 | 20,804,110 | 24.01 | 9,241 | 7,742 |
|  |  | Zt10-Ta_C_02 |  | 91,070,127 | 8,620,099 | 9.47 |  |  |
|  | D | Zt10-Ta_D_01 | 24 | 98,493,807 | 29,939,216 | 30.40 | 9,062 | 7,308 |
|  |  | Zt10-Ta_D_02 |  | 93,690,628 | 34,607,371 | 36.94 |  |  |

Overview of RNA-seq datasets including time point of sampling, number of sequenced reads post filtering, number of mapped reads, percentage of mapped reads, and numbers of transcribed genes. \* FPKM values were calculated using Cuffdiff2 and are normalized over all infection stages within the respective isolate (normalization method: geometric, dispersion method: per-condition). <sup>1</sup> 11,138 genes of IPO323 (94.08%) found by nucleotide blast for Zt05. <sup>2</sup> 11,754 of the 11,839 genes predicted and annotated for IPO323 [59]; 85 genes located on chromosome 18 were not considered. <sup>3</sup> 10,745 genes of IPO323 (90.76%) found by nucleotide blast for Zt10.

We obtained 89.2 to 147.5 million single-end, strand-specific reads per replicate (in total >2.7 billion reads) that were quality trimmed and filtered. Between 4.54% (early infection) and 76.4% (late infection) of the reads could be mapped to the genome of the respective isolate, reflecting the infection stage-specific amount of fungal biomass (Table 2, S5 Table, S1 Text). Across all isolates, transcriptomes of stages A and B, representing biotrophic growth, cluster together and are clearly different from transcriptomes of stages C and D that likewise cluster and represent necrotrophic growth of *Z. tritici* (S8 and S9 Figs). Exploring the transcriptome datasets based on gene read counts shows the greatest variation of biological replicates for Zt10 at stage C (S10 Fig), possibly reflecting variability in the infection development of the two biological replicates.

### Core *Zymoseptoria tritici* transcriptional program during wheat infection

The mean expression of genes located on accessory chromosomes was between 6-fold and 20fold lower than the expression levels of genes located on core chromosomes (S6 Table). We performed differential gene expression analyses to compare expression of the 10,426 *Z. tritici* core genes. We identified 597 genes that were differentially expressed between the infection stages (DESeq2, *P*_adj_ ≤ 0.01, |log_2_ fold change| ≥2) and show the same expression kinetics in all three isolates (Fig 4A). Interestingly, 79 of these genes were differentially expressed between several infection stages, suggesting dynamic, wave-like expression kinetics (S11 Fig). A total of 246 genes were differentially expressed (S7 Table) between stage A and stage B; the vast majority of these (242) were up-regulated in stage B. In stage A, three of the four genes that were up-regulated encode candidate secreted effector proteins (CSEPs), and the fourth encodes a carbohydrate active enzyme (CAZyme) similar to an extracellular chitosanase (*Zt09_chr_11_00040*). This gene is significantly down-regulated or not expressed during later infection (stages B to D), suggesting a role of the enzyme during early establishment in the leaf, similar to the role of a homolog described in *Fusarium solani* [66]. Another gene (*Zt09_chr_6_00402*) that was strongly up-regulated in all isolates during early infection encodes a putative hsp30-like small heat shock protein, possibly reflecting a response to stressful environmental conditions on the wheat leaf surface [67].

**Figure 4.**
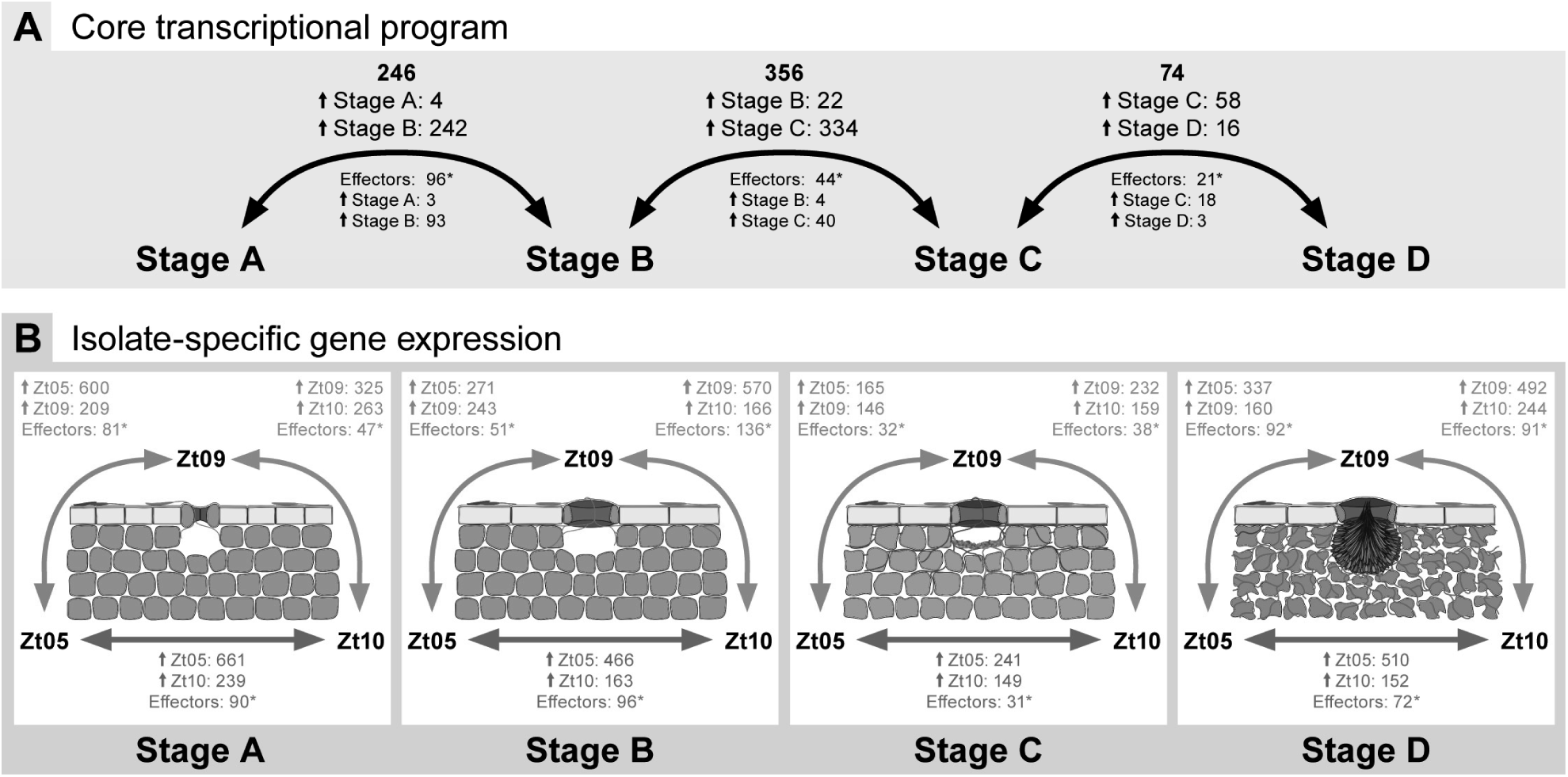
*Z. tritici* core transcriptional program during wheat infection and isolate-specific expression during the four infection stages. Numbers of significantly differentially expressed genes across all isolates **(A)** between the four core *Z. tritici* infection stages and **(B)** between the isolates within the infection stages (between Zt05 and Zt09: orange arrows, between Zt05 and Zt10: purple arrows, between Zt09 and Zt10: green arrows). Small arrows (Î) with stage or isolate names indicate the number of genes specifically up-regulated during that stage or in that isolate for the respective comparison. Differential gene expression analyses performed with DESeq2. Genes were considered to be significantly differentially expressed if *P*_adj_ ≤ 0.01 and |log_2_ fold change| ≥2. *Indicates significant enrichment of effector candidates among differentially expressed genes (Fischer's exact tests, *P* < 0.001). Effector candidates encode secreted proteins putatively involved in modulating molecular host-pathogen interactions [73].

The 242 genes up-regulated during biotrophic colonization are enriched with Gene Ontology (GO) groups involved in proteolysis (GO:0006508; 27 genes) and amino acid transmembrane transport (GO:0003333; 5 genes) (P < 0.01, Fischer’s exact test). Furthermore, three previously characterized LysM homologs [51] and two homologs (*Zt09_chr_11_00358, Zt09_chr_13_00167*) of *Ecp2,* an effector gene of the tomato-infecting fungus *Cladosporium fulvum* [68], are also strongly up-regulated during early infection, emphasizing the importance of these genes for biotrophic colonization. PFAM domain analysis further shows enrichment of genes encoding cytochrome P450- and polyketide synthase-like proteins that possibly play a role in the production of secondary metabolites (P < 0.001, *χ*^2^ test).

In stage B, 22 genes are up-regulated compared to stage C (S8 Table), including four genes encoding CSEPs of unknown function and a gene encoding the putative non-secreted catalase *Zt09_chr_6_00289.* Metabolite profiling showed that oxidative catabolism of lipids plays an important role for *Z. tritici* during biotrophic colonization [63]. High catabolic activity in the peroxisome entails accumulation of H_2_O_2_, which likely requires high abundance of catalase to maintain cellular redox homeostasis. *Zt09_chr_10_00421* is also highly expressed during biotrophic growth and down-regulated at later infection stages. It encodes a protein similar to siderophore iron transporter 1, previously described to be involved in the uptake of iron [69] which is essential for fungal growth and pathogenesis [70].

In stage C, 334 genes are significantly up-regulated compared to stage B (S8 Table) and 58 genes in comparison to stage D (S9 Table). Genes up-regulated from B to C are enriched with GO groups involved in metabolic processes (GO:0008152; 97 genes), in particular L-arabinose metabolic processes (GO:0046373; 4 genes), and transmembrane transport (GO:0055085; 25 genes). Similarly, a PFAM analysis shows an enrichment of genes encoding transporters; CAZymes including different groups of glycosyl hydrolases, serine hydrolases, alpha-L-arabinofuranosidases and cutinases that play important roles as plant tissue and cell wall degrading enzymes [71]; polyketide synthases; and cytochrome P450s. This transcriptional reprogramming reflects the physiological changes that *Z. tritici* undergoes during the transition from biotrophic to necrotrophic growth and is consistent with our microscopic observations. Among the 58 genes down-regulated from C to D (S9 Table) we identified GO groups involved in arabinan metabolic processes (GO:0031221; one gene) and an enrichment of PFAM domains related to beta-ketoacyl-ACP synthases, which are known to be involved in fatty acid production and important for the generation of new cell membrane, as well as cytochrome P450s, polyketide synthases, hydrophobic surface binding protein A [72], and tyrosinases.

Only 16 genes were significantly up-regulated from stage C to D, which is when the pycnidia mature (S9 Table), indicating overall similar transcription profiles during the two necrotrophic stages. Genes that are up-regulated during necrotrophic growth and reproduction are predicted to encode proteins similar to CAZymes, transporters, and proteins containing RNA-binding domains. Up-regulation of the secreted catalase-like protein-encoding gene *Zt09_chr_5_00821* shows the importance of detoxification of the ROS H_2_O_2_, which is highly abundant in necrotic leaf tissue as shown by DAB staining (Fig 2B).

In summary, we identified a core set of genes that show the same expression pattern in the three isolates during infection development. This core set includes genes encoding putative effectors as well as enzymes predicted to play a role in the breakdown and metabolism of plant cell components.

### Core biotrophic and necrotrophic effector candidates with shared expression profiles in *Z. tritici* isolates

Given their importance in plant-pathogen interactions, we particularly focused our analyses on genes encoding candidate secreted effector proteins (CSEPs) [73]. CSEP genes are significantly enriched among the core differentially expressed genes (P ≤ 2.7*10^13^, Fischer’s exact tests) (Fig 4A), indicating highly dynamic expression profiles of many core effectors during all stages of wheat infection. We filtered the differentially expressed CSEP genes according to their expression profiles (S12 Fig, S13 Fig) to identify putative key genes for biotrophic and necrotrophic growth inside the host (Table 3 and 4).

**Table 3.**
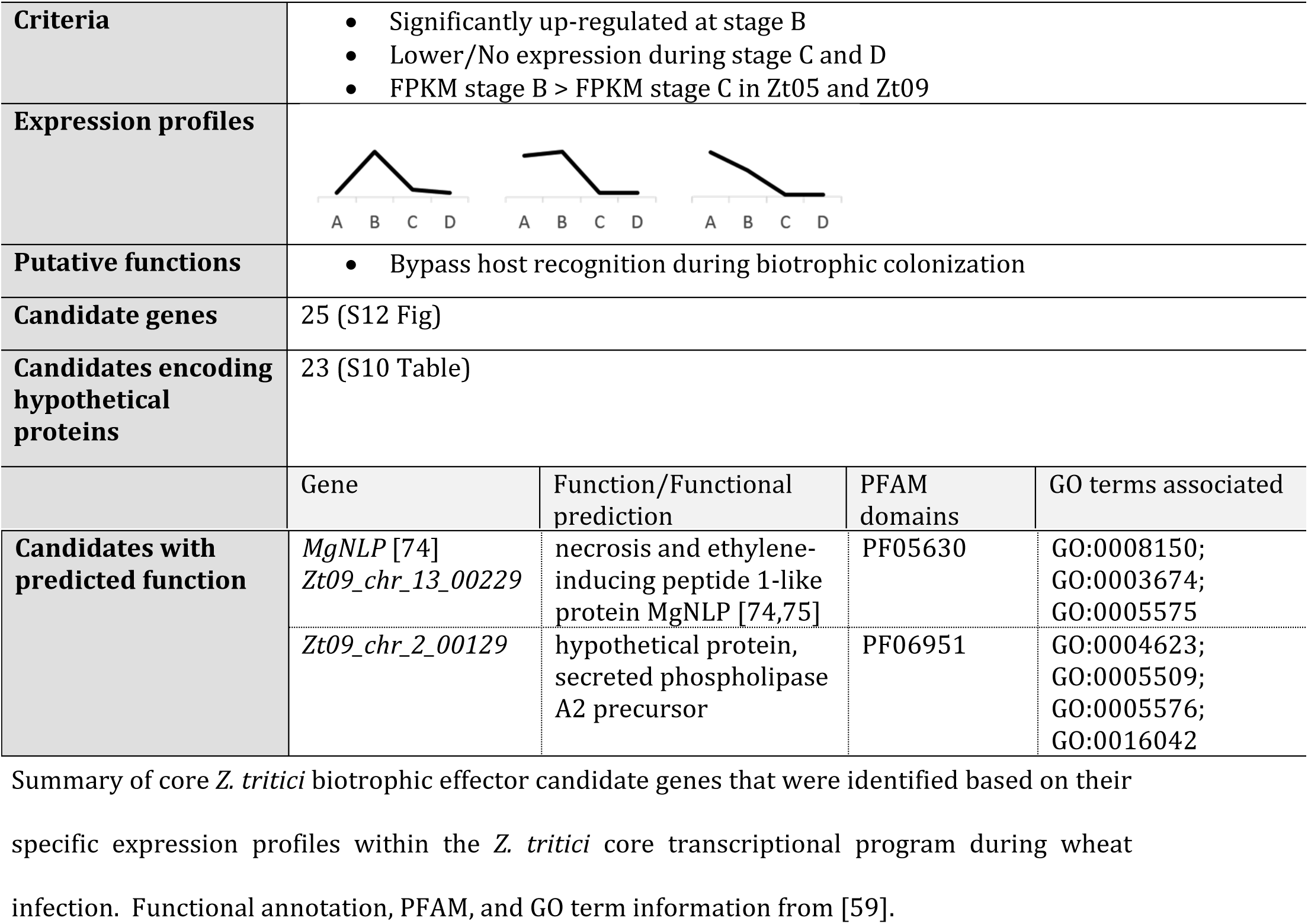
*Z. tritici* core biotrophic effector candidate genes

|  |  |  |  |  |
| --- | --- | --- | --- | --- |
| **Criteria** | Significantly up-regulated at stage B<br>Lower/No expression during stage C and D<br>FPKM stage B > FPKM stage C in Zt05 and Zt09 |  |  |  |
| **Expression profiles** |  |  |  |  |
| **Putative functions** | Bypass host recognition during biotrophic colonization |  |  |  |
| **Candidate genes** | 25 (S12 Fig) |  |  |  |
| **Candidates encoding hypothetical proteins** | 23 (S10 Table) |  |  |  |
|  | Gene | Function/Functional prediction | PFAM domains | GO terms associated |
| **Candidates with predicted function** | *MgNLP* [74]<br>*Zt09_chr_13_00229* | necrosis and ethylene-inducing peptide 1-like protein MgNLP [74,75] | PF05630 | GO:0008150;<br>GO:0003674;<br>GO:0005575 |
|  | *Zt09_chr_2_00129* | hypothetical protein, secreted phospholipase A2 precursor | PF06951 | GO:0004623;<br>GO:0005509;<br>GO:0005576;<br>GO:0016042 |

Summary of core *Z. tritici* biotrophic effector candidate genes that were identified based on their specific expression profiles within the *Z. tritici* core transcriptional program during wheat infection. Functional annotation, PFAM, and GO term information from [59].

**Table 4.**
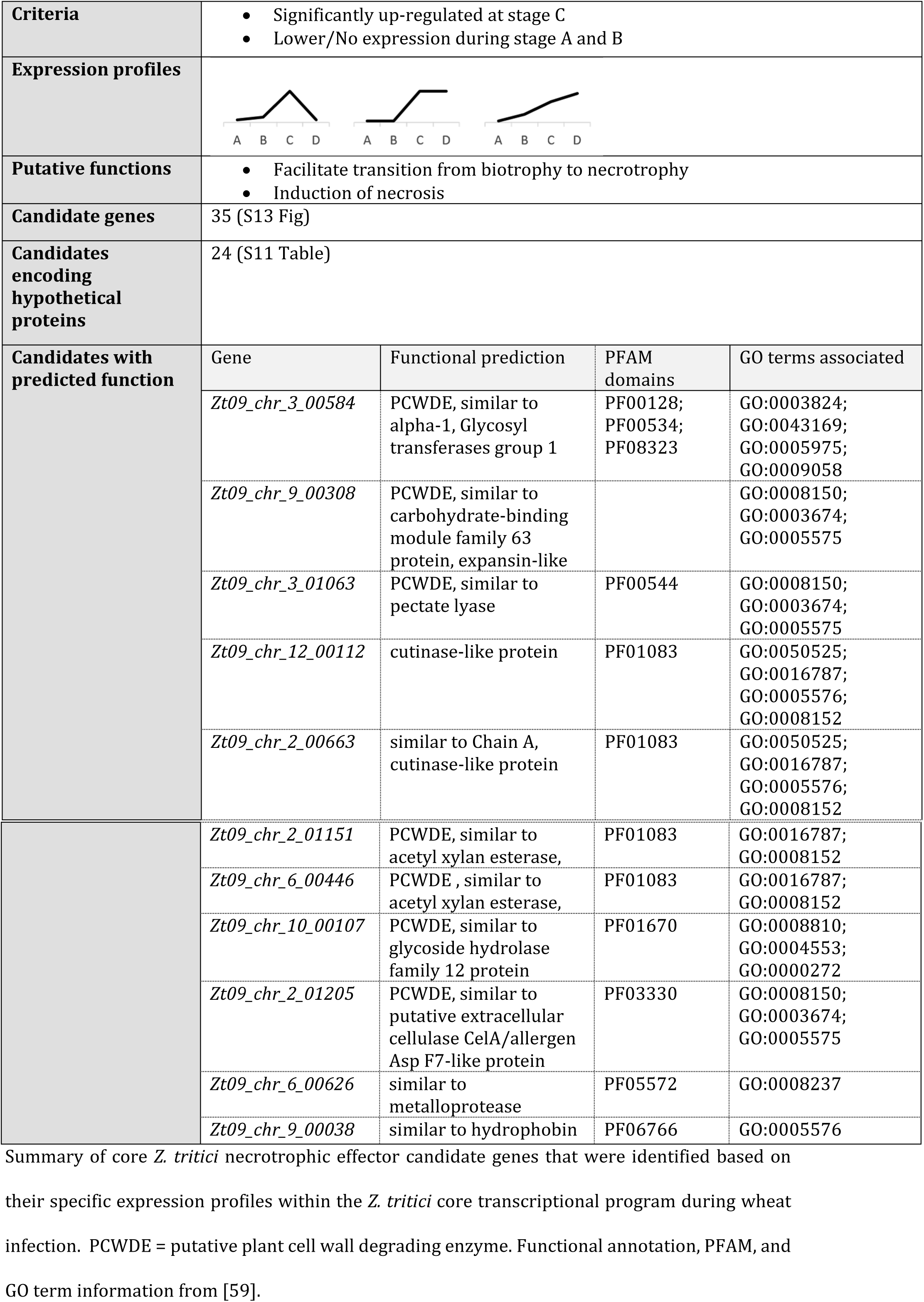
*Z. tritici* core necrotrophic effector candidate genes

|  |  |  |  |  |
| --- | --- | --- | --- | --- |
| **Criteria** | Significantly up-regulated at stage C<br>Lower/No expression during stage A and B |  |  |  |
| **Expression profiles** |  |  |  |  |
| **Putative functions** | Facilitate transition from biotrophy to necrotrophy<br>Induction of necrosis |  |  |  |
| **Candidate genes** | 35 (S13 Fig) |  |  |  |
| **Candidates encoding hypothetical proteins** | 24 (S11 Table) |  |  |  |
| **Candidates with predicted function** | Gene | Functional prediction | PFAM domains | GO terms associated |
|  | *Zt09_chr_3_00584* | PCWDE, similar to alpha-1, Glycosyl transferases group 1 | PF00128;<br>PF00534;<br>PF08323 | GO:0003824;<br>GO:0043169;<br>GO:0005975;<br>GO:0009058 |
|  | *Zt09_chr_9_00308* | PCWDE, similar to carbohydrate-binding module family 63 protein, expansin-like |  | GO:0008150;<br>GO:0003674;<br>GO:0005575 |
|  | *Zt09_chr_3_01063* | PCWDE, similar to pectate lyase | PF00544 | GO:0008150;<br>GO:0003674;<br>GO:0005575 |
|  | *Zt09_chr_12_00112* | cutinase-like protein | PF01083 | GO:0050525;<br>GO:0016787;<br>GO:0005576;<br>GO:0008152 |
|  | *Zt09_chr_2_00663* | similar to Chain A, cutinase-like protein | PF01083 | GO:0050525;<br>GO:0016787;<br>GO:0005576;<br>GO:0008152 |
|  | *Zt09_chr_2_01151* | PCWDE, similar to acetyl xylan esterase, | PF01083 | GO:0016787;<br>GO:0008152 |
|  | *Zt09_chr_6_00446* | PCWDE, similar to acetyl xylan esterase, | PF01083 | GO:0016787;<br>GO:0008152 |
|  | *Zt09_chr_10_00107* | PCWDE, similar to glycoside hydrolase family 12 protein | PF01670 | GO:0008810;<br>GO:0004553;<br>GO:0000272 |
|  | *Zt09_chr_2_01205* | PCWDE, similar to putative extracellular cellulase CelA/allergen Asp F7-like protein | PF03330 | GO:0008150;<br>GO:0003674;<br>GO:0005575 |
|  | *Zt09_chr_6_00626* | similar to metalloprotease | PF05572 | GO:0008237 |
|  | *Zt09_chr_9_00038* | similar to hydrophobin | PF06766 | GO:0005576 |

Summary of core *Z. tritici* necrotrophic effector candidate genes that were identified based on their specific expression profiles within the *Z. tritici* core transcriptional program during wheat infection. PCWDE = putative plant cell wall degrading enzyme. Functional annotation, PFAM, and GO term information from [59].

During symptomless stage B, 78 CSEP genes are specifically up-regulated, highlighting that a large suite of *Z. tritici* effectors is induced after stomatal penetration and is required for biotrophic colonization of the mesophyll. We narrowed down these genes to 25 core biotrophic CSEP genes (Table 3, S12 Fig, S10 Table) that mostly encode hypothetical proteins and may have a role in bypassing host recognition. In comparison to Zt05 and Zt09, expression of the biotrophic effectors in Zt10 is lower during stage B, and expression profiles often greatly deviated during the other infection stages. This diverging expression phenotype likely reflects the strongly limited biotrophic colonization that we observed for Zt10 (Fig 3B).

Among the biotrophic CSEP genes is *MgNLP* (*Zt09_chr_13_00229*) that encodes the necrosis and ethylene-inducing peptide 1-like protein MgNLP in *Z. tritici* [74]. *MgNLP* is strongly expressed before the transition to necrotrophy in wheat; however it only induces necrosis in dicots and its role in *Z. tritici* is still unknown [74,75].

A set of 35 CSEP genes are specifically up-regulated at stage C (S13 Fig) and represent candidates for necrotrophic core effectors (Table 4, S11 Table). These genes may be involved in the transition from biotrophic to necrotrophic growth and the induction of necrosis. 9 CSEP genes encode putative plant cell wall degrading enzymes and cutinase-like proteins, demonstrating that the lifestyle switch to necrotrophy involves intensified degradation of plant tissue and cell wall components. The gene *Zt09_chr_9_00038* encodes a putative hydrophobin; hydrophobins are small fungal-specific proteins with various functions [76], such as formation of a protective surface layer on hyphae and conidia [77] or as toxins in plant-pathogen interactions [78]. Further, *Zt09_chr_7_00263* encodes a putative secreted metalloprotease, which are known fungal virulence factors in animal pathogens [79] and are significantly induced during the transition to necrotrophy in the hemibiotrophic anthracnose-causing *Colletotrichum higginsianum* [80]. In *Fusarium verticillioides* and *F. oxysporum* f. sp. *lycopersici,* the secreted metalloproteases Fv-cmp and FoMep1 cleave antifungal extracellular host chitinases [81,82].

### Isolate-specific transcriptional changes during wheat infection

The 597 genes that we identified as differentially expressed between the stages show the same regulatory profile in each of the three isolates and we consider them as part of the core *Z. tritici* transcriptional infection program. However, we observed that transcript levels of many genes strongly deviate between the isolates within an infection stage.

To further study how the infection phenotypes of the *Z. tritici* isolates relate to differences in gene expression, we compared expression profiles during the infection stages (Fig 4B). In total, 2,377 (~22.8%) of the 10,426 shared genes are differentially expressed between the *Z. tritici* isolates during wheat infection (Table 5, S12 and S13 Tables), suggesting a high extent of redundancy and flexibility in the transcriptional program of *Z. tritici* during infection.

**Table 5.**
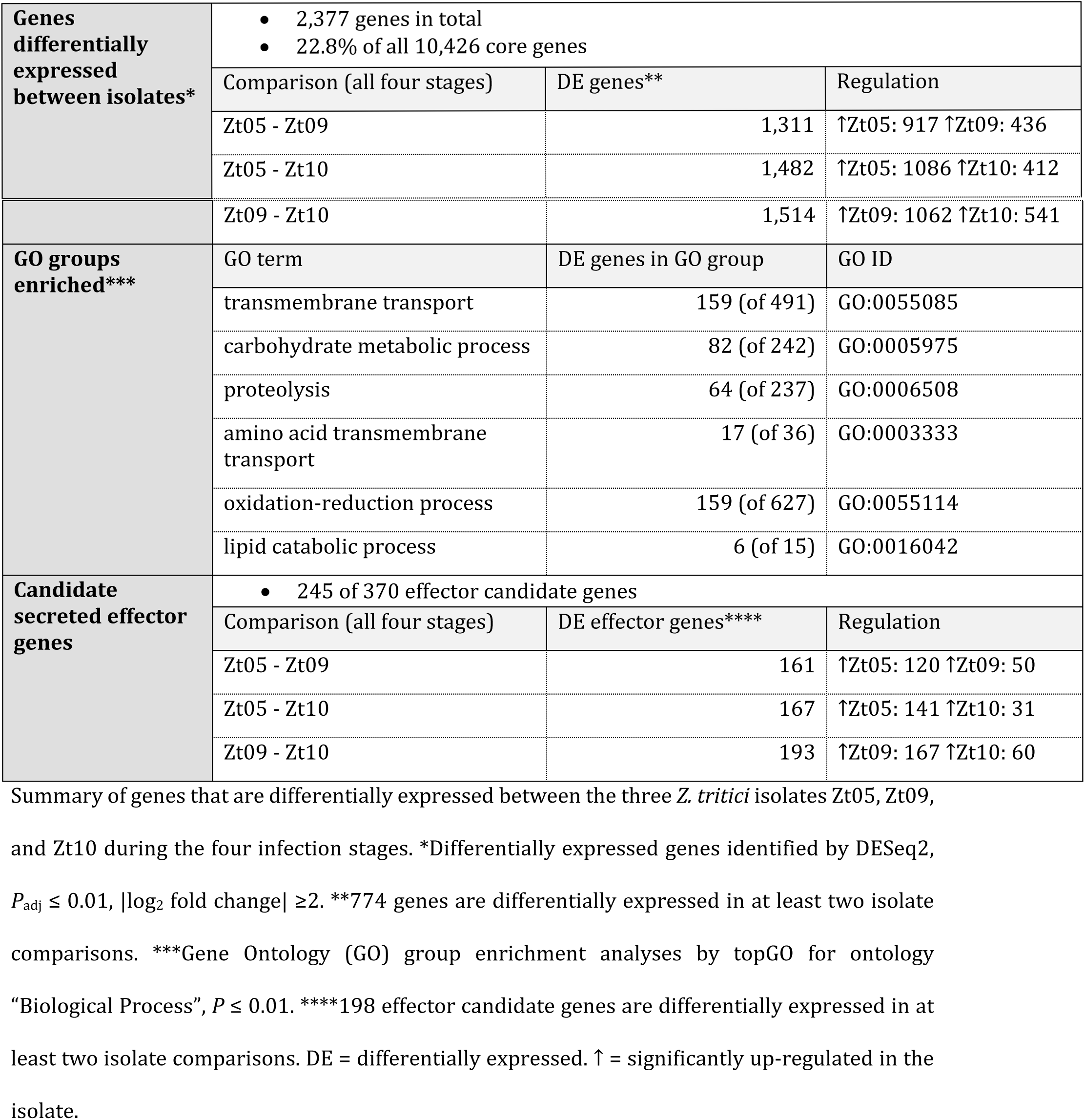
Genes with isolate-specific expression profiles during wheat infection

For all isolate comparisons, the identified differentially expressed genes are significantly enriched in CSEP genes (*P* ≤ 1.18*10^-7^, Fischer's exact tests) (Fig 4B) indicating isolate-specific effector transcriptional profiles (S14 Table). Figure 5 summarizes the expression kinetics of five CSEP genes in the three isolates during the four infection stages. These examples illustrate the differentiated expression profiles of CSEP genes in *Z. tritici* isolates during wheat infection (Fig 5) They encode one hypothetical effector (*Zt09_chr_12_00427*) and secreted proteins with various functions, including a hydrophobin (*Zt09_chr_9_00020*), a DNase (*Zt09_chr_2_01162*), and the ribonuclease Zt6 (*Zt09_chr_3_00610*), which possesses ribotoxin-like activity and is cytotoxic against plants and various microbes [83]. Among them, there is also a gene (*Zt09_chr_4_00039*) encoding a protein with homology to the phytotoxin cerato-platanin that was shown to induce necrosis and defense responses in the plant pathogen *Ceratocystis fimbriata* [84]. During all infection stages, *Zt09_chr_4_00039* is significantly higher expressed in Zt09 than in Zt10, as well as during stages B and D in comparison to Zt05 and might contribute to the higher necrosis levels caused by Zt09 (Fig 1A).

**Figure 5.**
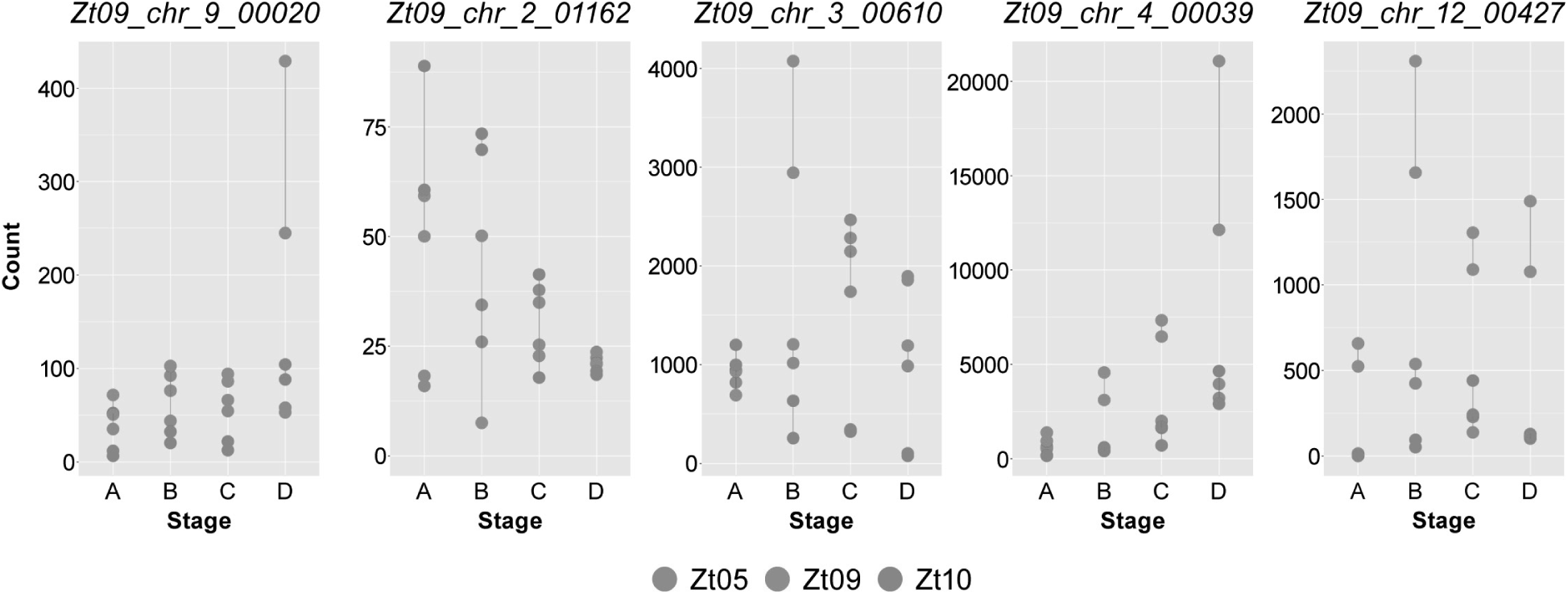
Five effector candidates with highly different expression profiles during wheat infection in the three isolates. The plots display normalized read counts for five effector candidate genes calculated by DESeq2 [95] for the twelve RNA-seq datasets. Read counts were normalized across the four core infection stages (A to D) and the three *Z. tritici* isolates Zt05, Zt09, and Zt10 and represent a measure of relative gene expression between the infection stages and between the isolates.

In addition to the differences in the expression of CSEP genes, we also noted isolate-specific expression patterns for genes located on accessory chromosomes. For example, three neighboring genes located on chromosome 19 in Zt09 (*Zt09_chr_19_00071*, *Zt09_chr_19_00072,* and *Zt09_chr_19_00073)* are significantly higher expressed in Zt10 during all four infection stages (S14 Fig, S13 Table), and other genes located ~70 kb downstream are likewise specifically up-regulated in Zt10 compared to one or both of the other isolates. In Zt10, these genes are located on unitig 16, which is syntenic to the accessory chromosome 19 of Zt09. However, transposable elements (RLG elements) are located downstream of *Zt09_chr_19_00072* and upstream of *Zt09_chr_19_00073* in Zt09, and these are not present within the vicinity of these genes on unitig 16 in Zt10. Instead, the right arm of unitig 16 of Zt10 (~100 MB) is inverted compared to Zt09 and Zt05, and the inversion starts up-stream of the gene *Zt09_chr_19_00073,* reflecting a significant sequence variation in the accessory chromosomes.

### Transposable elements are associated with the differentially expressed genes

Detailed analyses of the *Z. tritici* transcriptomes revealed considerable variation in the transcriptional landscapes among isolates. The extent of genetic differentiation between Zt05, Zt09, and Zt10 likely accounts for much of the transcriptional variation in the form of SNPs in regulatory sequences. However, other layers of gene regulation may also contribute to the heterogeneous transcriptional landscapes. We hypothesized that epigenetic transcriptional regulation, such as co-regulation of sequences associated with transposable elements, could impact gene expression variation. In a previous study, we showed that transposable elements and the accessory chromosomes of *Z. tritici* are enriched with the histone modifications H3K9me3 and H3K27me3, which are associated with repressive regions of chromatin [85]. In *Fusarium graminearum,* the histone modification H3K27me3 is associated with gene clusters encoding secondary metabolites and pathogenicity-related traits [86]. It is possible that variation in the distribution of histone modifications like H3K27me3 across the genome sequences of Zt05, Zt09, and Zt10 contributes to the dramatic variation in expression phenotypes.

To test this, we assessed the distances of all genes to the closest annotated transposable element. In the genomes of all three isolates, we found that isolate-specific differentially expressed genes are located significantly closer to transposable elements than genes that were not differentially expressed (Mann-Whitney U tests, *P* < 2.2*10^-16^). Within the differentially expressed genes, isolate-specific up-regulated genes (genes that are significantly up-regulated in one isolate in contrast to the others) are significantly enriched within a distance of 2 kb to transposable elements in Zt05 and Zt09 (Fisher’s exact tests, p ≤ 0.0094), but not in Zt10. We note however, that the transposable element annotation in Zt10 is not as complete as in Zt05 and Zt09 as it was based on the Illumina short read assembly which is 6.7 Mb smaller than the *de novo* genome assembly of SMRT Sequencing reads. We also analyzed the *in vitro* histone 3 methylation data for Zt09 [85] and found that differentially expressed genes are indeed significantly closer to H3K9me3 and H3K27me3 peaks (Mann-Whitney U tests, *P* < 2.2*10^-16^). Further, we observed significant enrichment of genes significantly up-regulated in Zt09 in comparison to Zt05 and Zt10 within a distance of 2 kb to H3K9me3 and H3K27me3 peaks (Fisher’s exact tests, *P* < 1.55*10^-15^), but down-regulated genes were only enriched in the vicinity of H3K27me3 peaks (distance ≤2 kb, Fisher’s exact test, *P* = 1.35*10^-15^). Poor transcription of genes located on the accessory chromosomes was explained by enrichment of H3K27me3 covering the entire chromosomes and H3K9me3, which is mostly associated with repetitive DNA [85]. Our findings indicate that during host infection chromatin state of repeat-rich genome compartments is highly dynamic and changes between “active” euchromatin and “repressive” heterochromatin, as suggested in *Leptosphaeria maculans* [87]. Further, our observation suggests that the fine-scale distribution of epigenetic marks likely differs between the genomes of *Z. tritici* isolates and contributes to the isolate-specific gene expression phenotypes that we observed. To further visualize the transcriptional landscape across the three *Z. tritici* genomes, we calculated expression values (FPKM) of 1 kb windows (S6 Table) and plotted them in heatmaps along the chromosomes and unitigs (S15-S17 Figs). This approach allows visualization of the transcriptional landscapes at a high resolution and resulted in the identification of heterogeneous gene expression patterns across chromosomes, such as chromosome 19 in Zt10 (S17 Fig), and suggests conservation of previously identified patterns. Almost no loci on the right arm of chromosome 7 were transcribed in Zt05, Zt09, and Zt10 (S16 Fig), as was previously found in Zt09 and IPO323 [30,63]. This chromosomal segment has characteristics of an accessory chromosome, as it is significantly enriched with H3K27me3 that mediates transcriptional silencing [85]. While syntenic chromosomal regions generally have a similar composition of transcribed and silenced loci, the fine-scale distribution of transcriptional cold- and hot-spots is clearly different between the genomes of the three isolates studied.

## Conclusion

We conducted a detailed comparison of infection phenotypes of three pathogenic *Z. tritici* isolates that are equally virulent in a susceptible host genotype and show an unexpectedly high extent of plasticity in the infection program of a fungal plant pathogen. The three isolates differ significantly in their genomic composition, and we show that the genetic variation of the three isolates translates into highly distinct infection phenotypes that deviate temporally and spatially. The transcriptional programs associated with host colonization show a high degree of variability between the three isolates: more than 20% of the core genes are differentially expressed between the three *Z. tritici* isolates during the four infection stages. This suggests strong redundancy in the *Z. tritici* “infection program” between isolates. Effector candidates are enriched among the differentially expressed genes, suggesting that the three isolates employ different molecular strategies to manipulate host defenses. Strikingly, highly variable infection programs result in the same level of virulence, showing that “host specialization” in *Z. tritici* involves a very flexible strategy to exploit wheat tissue for growth and reproduction. As necrotic lesions are usually composed of several distinct *Z. tritici* genotypes, it is highly relevant to investigate whether strains in one lesion have similar or different infection phenotypes. Various infection strategies within a lesion could complement each other or, in contrast, have antagonistic effects and facilitate competition.

An intriguing question that emerges from our analyses is which factors cause deviation in gene expression phenotypes in *Z. tritici*. Genetic variants associated with transcriptional regulation likely contribute to differences in gene regulation. However, we hypothesize that variation in epigenetic traits promotes different transcriptional programs. Genome-wide patterns of transcriptional activity (S15-S17 Figs) indeed suggest some variation in the physical distribution of transcriptionally active and silent regions, which may result from distinct epigenetic landscapes related to histone modifications or DNA methylation.

We hypothesize that highly diverging infection phenotypes are not exclusive among isolates of *Z. tritici* and are likely found in populations of other pathogens that retain high levels of genetic diversity. Variation in infection and expression profiles contributes another layer of polymorphism to pathogen populations and may be important for the pathogen to rapidly adjust to environmental changes. Hence, the resulting diversity of infection phenotypes needs to be acknowledged to understand pathogen evolution and develop sustainable crop protection strategies.

## Materials and Methods

### Isolates and growth conditions

Cells of *Zymoseptoria tritici* isolates (S1 Table) were inoculated from glycerol stocks onto YMS agar (0.4% [w/v] yeast extract, 0.4% [w/v] malt extract, 0.4% [w/v] sucrose, 2% [w/v] bacto agar) and grown at 18°C for 5 days. Single cells were grown in liquid YMS (200 rpm, 18°C) for 2 days and harvested by centrifugation (3500 rpm for 10 min).

### Plant infection experiments

For all plant infection experiments, we used 14-day-old seedlings of the winter wheat (*Triticum aestivum)* cultivar Obelisk (Wiersum Plantbreeding, Winschoten, Netherlands). The fungal inoculum was adjusted to 1 × 10^8^ cells/mL in 0.1% [v/v] Tween 20 (Roth, Karlsruhe, Germany) and brushed onto labeled areas (8 to 12 cm) of the second leaf of each plant. The same treatment without fungal cells was conducted for mock controls. After inoculation, plants were incubated at 22°C [day]/20°C [night] and 100% humidity with a 16-h light period for 48 h. Then, humidity was reduced to 70%. Plants were grown for 3 or 4 weeks after inoculation, depending on the experiment.

### *In planta* phenotypic assays

To compare quantitative virulence of Zt05, Zt09, and Zt10 on wheat, we performed three independent, randomized infection experiments with blinded inoculation and evaluation. Inoculated leaf areas of 460 leaves were evaluated at 28 days post infection (dpi) by scoring the observed disease symptoms based on the percentage of leaf area covered by necrosis and pycnidia as previously described [88]. We differentiated six categories: 0 (no visible symptoms), 1 (1-20%), 2 (21-40%), 3 (41-60%), 4 (61-80%), and 5 (81-100%). Statistical differences were evaluated by Mann-Whitney *U* test considering differences significant if *P* ≤ 0.01.

To compare the temporal development of disease, we manually inspected 40 inoculated leaves between 9 and 27 dpi and registered the occurrence of first visible symptoms every two days. Individual leaves were visualized using a Leica S8APO equipped with a Leica DFC450 camera.

To localize accumulation of the reactive oxygen species H_2_O_2_ within infected leaf tissue, we conducted 3,3’-diaminobenzidine (DAB) staining [89] at 4, 10, 14, 18, and 21 dpi (S2 Text). The presence of H_2_O_2_ is indicated by reddish-brown precipitate in cleared leaves. Samples were documented before (iPhone 7 camera) and after (Canon EOS 600D) staining.

### Phenotypic assays *in vitro*

To compare tolerance towards stress conditions and assess the *in vitro* phenotypes of the *Z. tritici* isolates, we conducted a stress assay as previously described [88]. After five days, we compared growth on solid YMS medium at 18°C to growth on YMS medium exposed to stress conditions: temperature (20/22°C with 16-h day/8-h night rhythm, 28°C in darkness), oxidative stress (2 and 3 mM H_2_O_2_), osmotic stress (1 M NaCl, 1 M sorbitol), and cell wall stress (500 μg/mL Congo red, 200 μg/mL calcofluor white). Colony development was documented using a Canon EOS 600D. Each stress treatment was replicated three times.

### Analysis of *Z. tritici* wheat infection by confocal microscopy

Structures of *Z. tritici* isolates inside and on the surface of wheat leaves were analyzed by confocal laser scanning microscopy. We harvested infected wheat leaves at 3-5, 7, 8, 10-14, 17, 19-21, 25, and 28 dpi and analyzed the interactions between fungal hyphae and wheat tissue. Likewise, we analyzed infected leaves to determine the infection stage of leaf samples used for RNA extraction (see below). In total, we studied 37 infected wheat leaves for Zt05, 34 for Zt09, and 30 for Zt10, analyzed at least 15 infection events per leaf sample by confocal microscopy, and created a total of 113 confocal image z-stacks. Cleared leaf material was stained with wheat germ agglutinin conjugated to fluorescein isothiocyanate (WGA-FITC) in combination with propidium iodide (PI) (S2 Text for staining protocol). Microscopy was conducted using a Leica TCS SP5 (Leica Microsystems, Germany) and a Zeiss LSM880 (Carl Zeiss Microscopy, Germany). FITC was excited at 488 nm (argon laser) and detected between 500-540 nm. PI was excited at 561 nm (diode-pumped solid-state laser) and detected between 600-670 nm. Image stacks were obtained with a *x/y* scanning resolution of 1024 x 1024 (Leica) or 1500 x 1500 pixels (Zeiss) and a step size of 0.5 - 1 μm in *z.* Analyses, visualization, and processing of image z-stacks were performed using Leica Application Suite Advanced Fluorescence (Leica Microsystems, Germany), ZEN black and Zen blue (Carl Zeiss Microscopy, Germany), and AMIRA^®^ (FEI™ Visualization Science Group, Germany). Animations of image z-stacks are .avi format and can be played in VLC media player (available at http://www.videolan.org/vlc/).

### Transcriptome analyses of *Z. tritici* isolates during wheat infection

High-quality total RNA from *Z.* tritici-infected wheat material was isolated using the TRIzol™ reagent (Invitrogen, Karlsruhe, Germany) according to the manufacturer’s instructions. Material of three wheat leaves was harvested synchronously, pooled, and immediately homogenized in liquid nitrogen. The resulting leaf powder (100 mg) was used for RNA extraction. Because our analyses revealed differences in the temporal development of infection between the isolates, we set up independent sampling schedules for each isolate to compare transcriptomes of the same infection stage (S4 Table). We collected infected leaf material at one to three determined time points and assigned infection stage based on examination of central sections (1 - 2 cm) of each leaf by confocal microscopy (S7 Fig). We determined the best representatives of each infection stage and chose two samples per isolate at each stage as biological replicates for transcriptome sequencing (Table 2). Preparation of strand-specific RNA-seq libraries including polyA enrichment was performed at the Max Planck Genome Center, Cologne, Germany (http://mpgc.mpipz.mpg.de) using the NEBNext Ultra™ Directional RNA Library Prep Kit for Illumina according to the manufacturer’s protocol (New England BioLabs, Frankfurt/Main, Germany) with an input of 1 μg total RNA. Sequencing, performed using an Illumina HiSeq 2500 platform, generated strand-specific, 100-base, single-end reads with an average yield of 112 million reads per sample (S5 Table). We assessed the quality of sequencing data with *FastQC* v0.11.2 (http://www.bioinformatics.babraham.ac.uk/projects/fastqc/), removed residual TruSeq adapter sequences, and applied a stringent read trimming and quality filtering protocol using *FASTX-toolkit* v0.0.14 (http://hannonlab.cshl.edu/fastx_toolkit/) and *Trimmomatic* [90] v0.33 (S3 Text for details). The resulting 88-bp reads were mapped against the genome of the respective *Z. tritici* isolate with *TopHat2* v2.0.9 [91]. Read alignments were stored in SAM format, and indexing, sorting, and conversion to BAM format was performed using SAMtools v0.1.19 [92]. The relative abundance of transcripts for predicted genes was calculated in FPKM by *Cuffdiff2* v2.2.1 [93]. Total raw read counts per gene were estimated with *HTSeq* v0.6.1p1 using union mode [94]. Gene coordinates in the Zt05 (S15 Table) and Zt10 (S16 Table) genomes were obtained by mapping the predicted genes of IPO323 using nucleotide BLAST alignments (e-value cutoff 1e^-3^, identity ≥90%, query coverage between 90% and 110%). Differential gene expression analyses between *Z. tritici* infection stages and isolates were performed in R using the Bioconductor package *DESeq2* v1.10.1 [95]. Significantly differentially expressed genes were determined with *P*_adj_ ≤ 0.01 and |log**2** fold-change ≥2|. The R package topGO [96] was used to perform Gene Ontology (GO) term enrichment analyses within the differentially expressed genes. *P* values for each GO term [59] were calculated using Fischer’s exact test applying the topGO algorithm “weight01” that takes into account GO term hierarchy. We reported categories significant with *P* ≤ 0.01 for the ontology “Biological Process”. PFAM domain enrichment analyses were performed using a custom python script, and *P* values were calculated using χ^2^ tests. To analyze genomic distances between differentially expressed genes and transposable elements (TEs), we annotated TEs as described in [59] for Zt05 (S17 Table) and Zt10 (S18 Table) and used the published TE annotation of IPO323 for Zt09 [59]. Distances between the genes of interest and the closest annotated TEs were calculated with bedtools v2.26 [97]. Likewise, we used ChIP-seq peak data [85] to calculate distances between genes and the closest H3K9me3 and H3K27me3 peaks. Statistical analyses were performed in R. For an overview of all programs and codes used to process and analyze transcriptome data, including the applied settings and parameters, see S3 Text.

### Pulsed-field gel electrophoresis

A non-protoplast protocol (S2 Text) was used to produce DNA plugs for separation of small chromosomes (~0.2 to 1.6 Mb) by pulsed-field gel electrophoresis (PFGE) [98]. Chromosomal DNA of *Saccharomyces cerevisiae* (Bio-Rad) was used as standard size marker. Gels were stained for 30 min in 1 μg/mL ethidium bromide solution, and chromosome bands were detected with Thyphoon Trio™ (GE).

### *De novo* genome assemblies of *Z. tritici* isolates Zt05 and Zt10 and synteny analyses

High molecular weight DNA of Zt05 and Zt10 was extracted from single cells grown in liquid YMS, using a modified version of the cetyltrimethylammonium bromide (CTAB) extraction protocol [99], and used as input to prepare Pacific Biosciences (PacBio) SMRTbell libraries that were size-selected with a 10- to 15-kb cut-off. Single-molecule real-time (SMRT) sequencing was performed on four SMRT cells and run on a PacBio RS II instrument at the Max Planck Genome Center in Cologne, Germany (http://mpgc.mpipz.mpg.de). Genome assemblies of Zt05 and Zt10 based on the generated PacBio long reads were done as previously described [56] using *HGAP* [100] v3.0 included in the *SMRTanalysis suite* v2.3.0. Briefly, we applied default settings for *HGAP* runs and tested the influence of different minimum seed read lengths (13 kb, 15 kb, 19 kb, and 21 kb) used for initiation of self-correction. A 19-kb minimum seed read length cut-off generated the most favorable results in terms of pre-assembly yield, assembly N50, and length of total assembly. Assembled unitigs were polished by applying default settings of *Quiver,* which is part of the *SMRTanalysis suite.* Unitigs in which median PacBio read coverage deviated more than a factor of 1. 5X from all contigs were removed from the final assemblies. Synteny of *Z. tritici* reference strain IPO323 and the Zt05 and Zt10 unitigs was compared using SyMAP [101] v4.2 applying default settings and considering all unitigs ≥1,000 kb (Zt05) and ≥10,000 kb (Zt10). To estimate the amount of unique DNA in Zt05 and Zt10 in comparison to IPO323 and Zt09 respectively, we generated pairwise genome alignments with *Mugsy* v1.r2.2 [102] applying default settings. Alignments were analyzed using a custom python script to extract unique DNA blocks with a minimum length of 1 bp.

## Data availability

All generated RNA-seq datasets have been deposited at the NCBI Gene Expression Omnibus and are accessible with the accession number GSE106136. *De novo* genome assemblies of isolates Zt05 and Zt10 are available under accession numbers PEBP00000000 and PEBO00000000. The genome sequence of the reference isolate IPO323 used for transcriptome analysis of Zt09 is available at: http://genome.jgi.doe.gov/Mycgr3/Mycgr3.home.html. Genome assemblies based on whole genome shotgun sequencing (Illumina) were also used; the assembly for Zt10 is available at GenBank (Zt10 = STIR04_A26b) GCA_000223645.2. Sequencing data and assembly for Zt05 (Zt05 = MgDk09_U34) are available through NCBI BioProject PRJNA312067 [57].

## Acknowledgements

We thank Ronny Kellner for insightful comments to a previous version of this manuscript, Julien Y. Dutheil for support with comparative transcriptome analyses and Petra Happel for help with the *in vitro* stress assays.

## Author contributions

Conceptualization: JH, EHS. Investigation: JH, MM, HS. Genome and transcriptome sequencing data curation and analyses: JH, MM, CJE, JG, EHS. Confocal microscopy analyses and data visualization: JH, HA. Preparation and writing of manuscript: JH, EHS. Editing of manuscript: JH, MM, HS, EHS.

## Supporting Information

**S1 Table. *Zymoseptoria tritici* isolates used in this study.**

**S2 Table. SMRT Sequencing-based *de novo* genome assemblies, synteny analyses, and conserved genes.**

The table summarizes basic statistics of *de novo* genome assemblies generated for isolates Zt05 and Zt10 based on SMRT Sequencing long-read data, results of synteny analyses between IPO323/Zt09 chromosomes and Zt05 and Zt10 unitigs, and the presence/absence of IPO323/Zt09 genes and effector candidates in the genomes of Zt05 and Zt10.

**S3 Table. *Z. tritici* core genes and effector candidates.**

List of 10,426 genes and 370 effector candidate genes that are present in the genomes of the *Z. tritici* isolates Zt05, Zt09, and Zt10 based on nBLAST analyses. Genes are based on the *Z. tritici* genome annotation [59] and effector gene candidates were predicted by [6].

**S4 Table. Isolate-specific sampling schedules for transcriptome sequencing of infection stages.**

Post-inoculation time points were scheduled based on previous plant infection experiments to cover each of the four *Z. tritici* infection stages. Samples for transcriptome sequencing and analyses were collected at one to three different times points and eventually selected based on the results of microscopic analyses of central leaf sections. Selected samples are marked by *.

**S5 Table. Detailed overview of stage-specific transcriptomes of *Z. tritici* isolates during wheat infection generated in this study.**

**S6 Table. Comparison of expression on core and accessory chromosomes of *Z. tritici* during wheat infection.**

FPKM expression values calculated with Cuffdiff2 [93] for genes and 1-kb windows located on the core (CC) and accessory (AC) chromosomes.

**S7 Table. Genes that are significantly differentially expressed between infection stage A and B across all *Z. tritici* isolates.**

**S8Table. Genes that are significantly differentially expressed between infection stage B and C across all *Z. tritici* isolates.**

**S9 Table. Genes that are significantly differentially expressed between infection stage C and D across all *Z. tritici* isolates.**

**S10 Table. *Z. tritici* biotrophic core effector candidates.**

**S11 Table. *Z. tritici* necrotrophic core effector candidates.**

**S12 Table. Core *Z. tritici* genes that are differentially expressed between the three isolates during wheat infection.**

**S13 Table. Core *Z. tritici* genes that are differentially expressed between the three isolates during wheat infection, sorted by up-regulation per isolate.**

**S14 Table. *Z. tritici* effector candidate genes that are differentially expressed between the three isolates during wheat infection.**

**S15 Table. Gene annotation for the *Z. tritici* isolate Zt05.**

Gene annotation in .gff file format for the Zt05 genome assembly based on Illumina short reads (NCBI BioSample: SAMN04494882).

**S 16Table. Gene annotation for the *Z. tritici* isolate Zt10.**

Gene annotation in .gff file format for the Zt10 genome assembly based on Illumina short reads (NCBI accession number: GCA_000223645.2).

**S17 Table. Transposable element annotation for the *Z. tritici* isolate Zt05.**

Transposable element annotation in .gff file format for the Zt05 genome assembly based on Illumina short reads (NCBI BioSample: SAMN04494882).

**S18 Table. Transposable element annotation for the *Z. tritici* isolate Zt10.**

Transposable element annotation in .gff file format for the Zt10 genome assembly based on Illumina short reads (NCBI accession number: GCA_000223645.2).

**S1 Figure.**
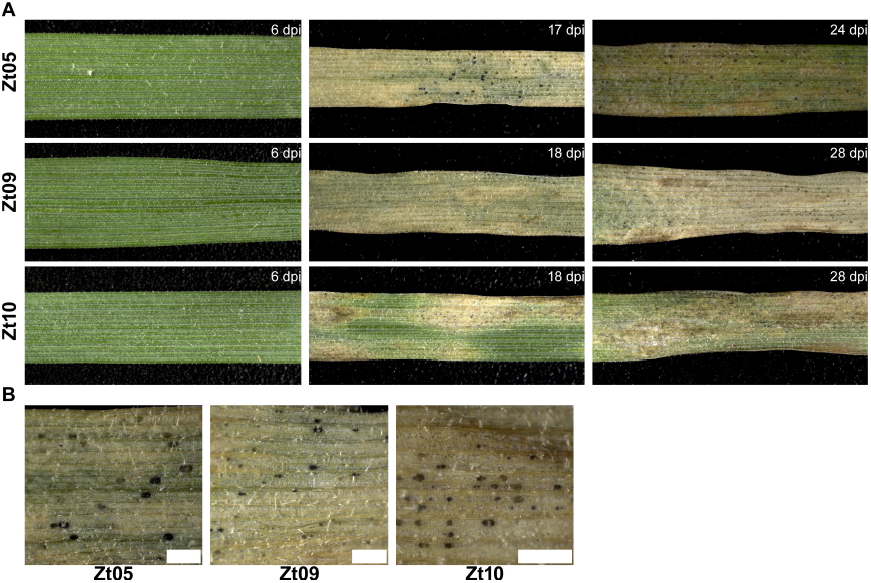
Disease development on wheat leaves infected with *Z. tritici* isolates Zt05, Zt09, and Zt10. (A) Photographs of wheat leaves taken at different time points after inoculation with *Z. tritici* isolates Zt05, Zt09, and Zt10. **(B)** Infected wheat leaves contained similar numbers of pycnidia. Scale bars = 500 μm.

**S2 Figure.**
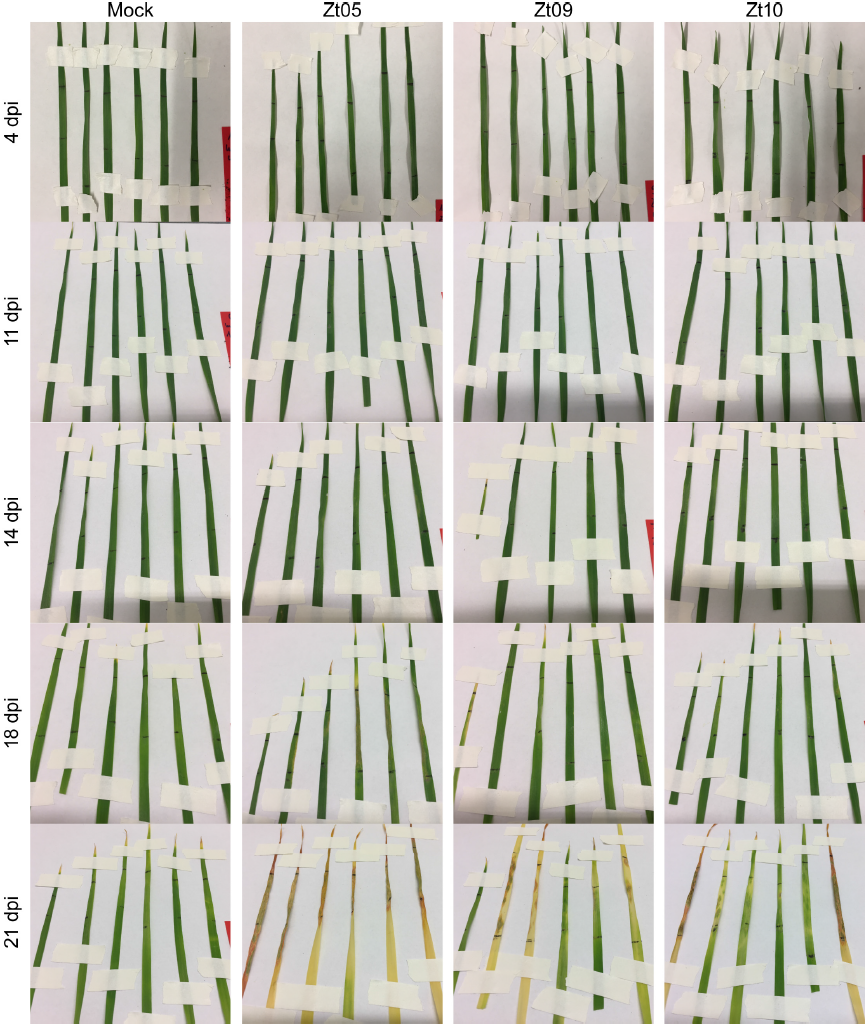
Symptom development on wheat leaves used for the ROS staining assay. Photographs of *Triticum aestivum* cv. Obelisk leaves taken at 4, 11, 14, 18, and 21 days post inoculation with *Z. tritici* isolates Zt05, Zt09, and Zt10 and mock treatment. Leaves were subsequently subjected to ROS detection staining.

**S3 Figure.**
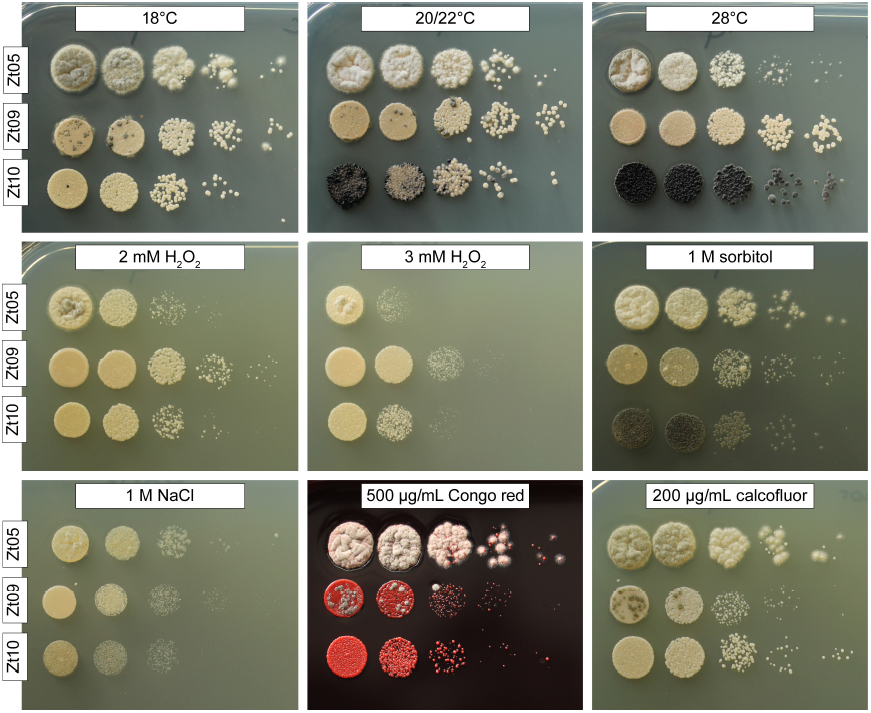
Differences in colony morphology and abiotic stress tolerance between *Z. tritici* isolates. Growth of the isolates Zt05, Zt09, and Zt10 was tested under multiple stress conditions in comparison to the standard cultivation condition *in vitro* (solid YMS medium at 18°C, no light): growing conditions of wheat (20/22°C at 16-h day/8-h night rhythm), heat stress (28°C), oxidative stress (2 and 3 mM H_2_O_2_), osmotic stress (1 M sorbitol, 1 M NaCl), and cell wall stress (500 μg/mL Congo red, 200 μg/mL calcofluor white).

**S4 Figure.**
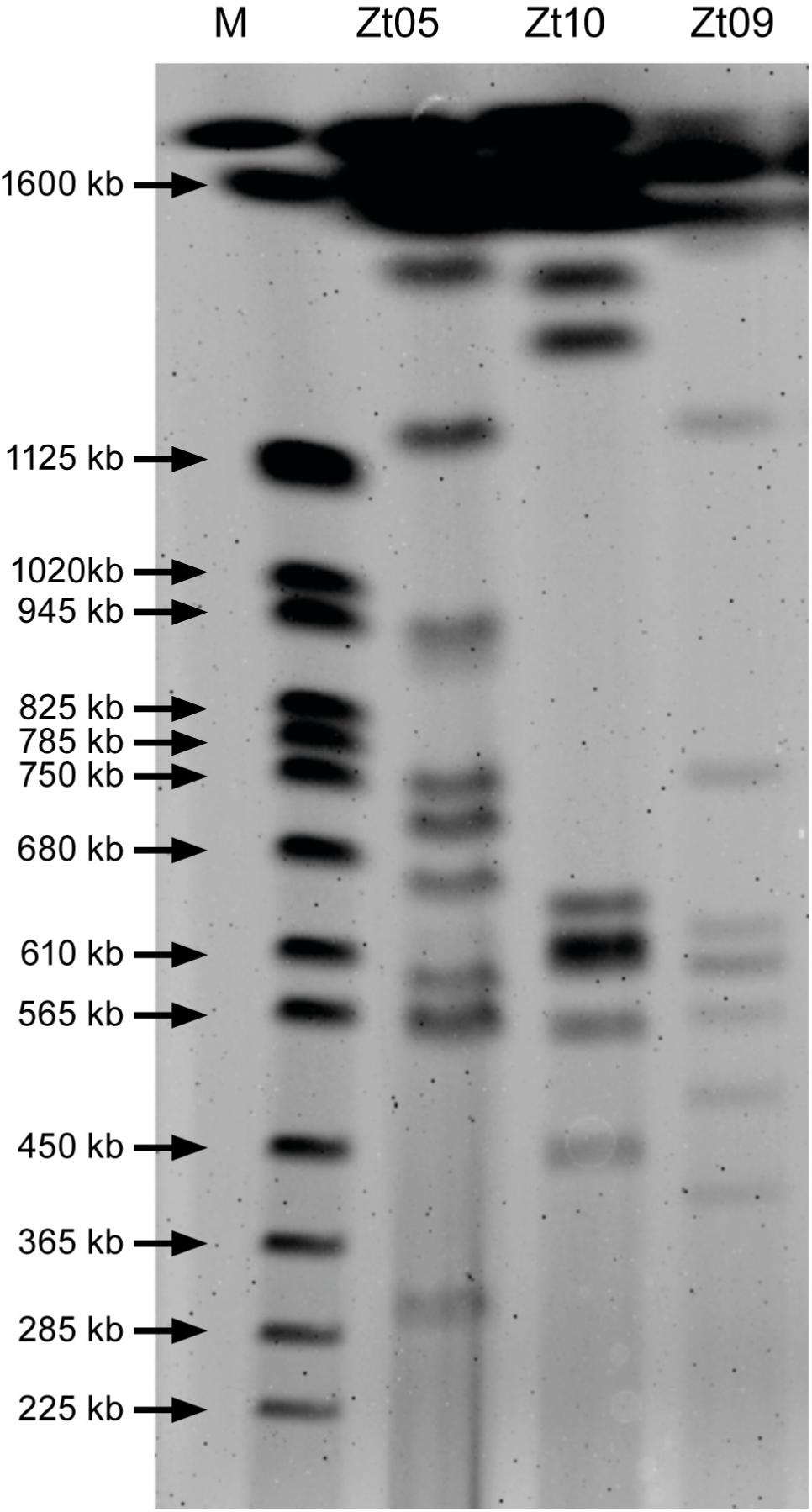
Karyotype variation of *Z. tritici* field isolates. Pulsed-field gel electrophoresis shows number and size variations for small chromosomes (~225 to 1,460 kb) of *Z. tritici* isolates Zt05, Zt10, and Zt09. Standard chromosome size marker (M): *Saccharomyces cerevisiae.*

**S5 Figure.**
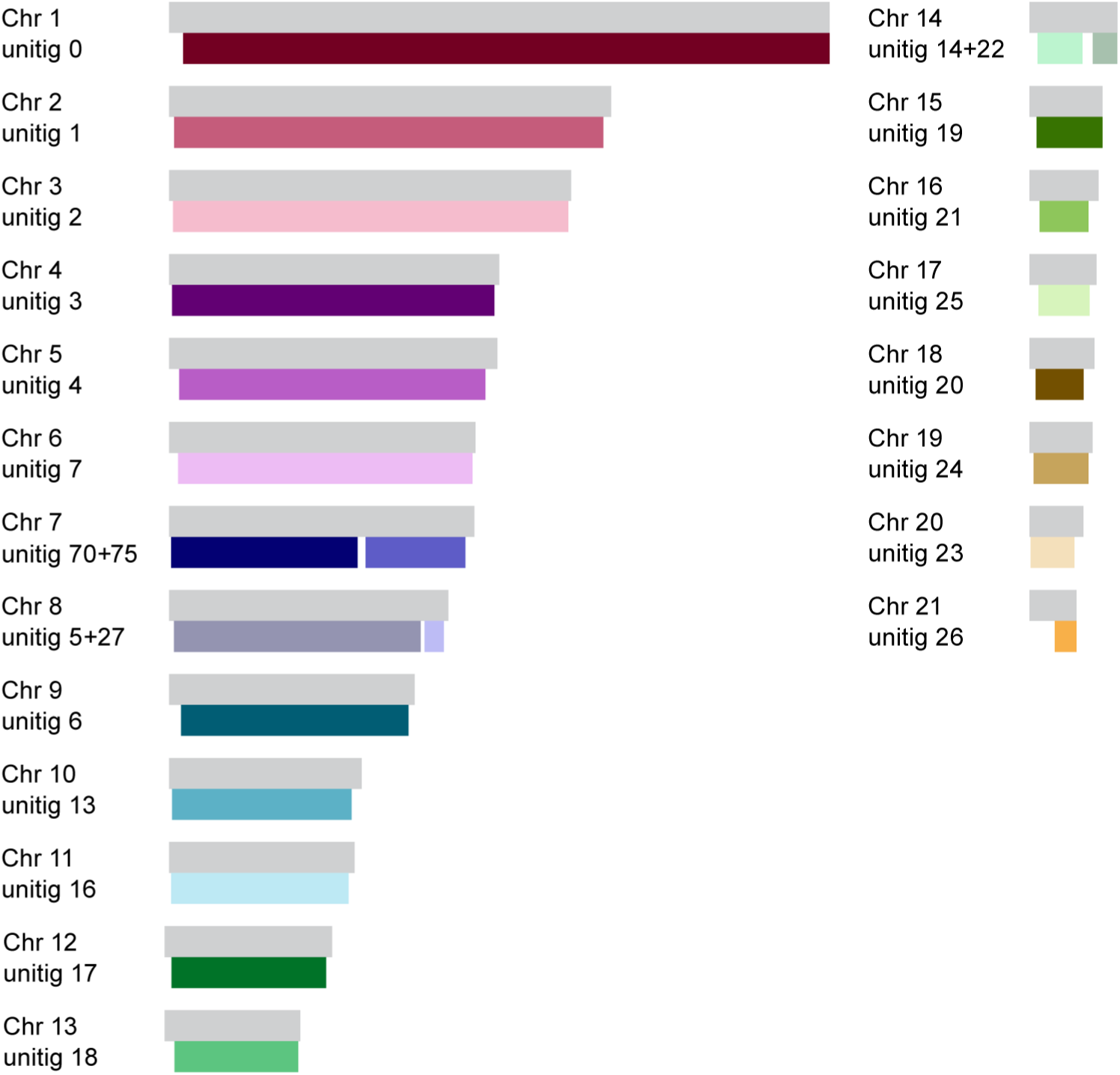
Synteny between Zt05 *de novo* assembled unitigs and IPO323 chromosomes.

**S6 Figure.**
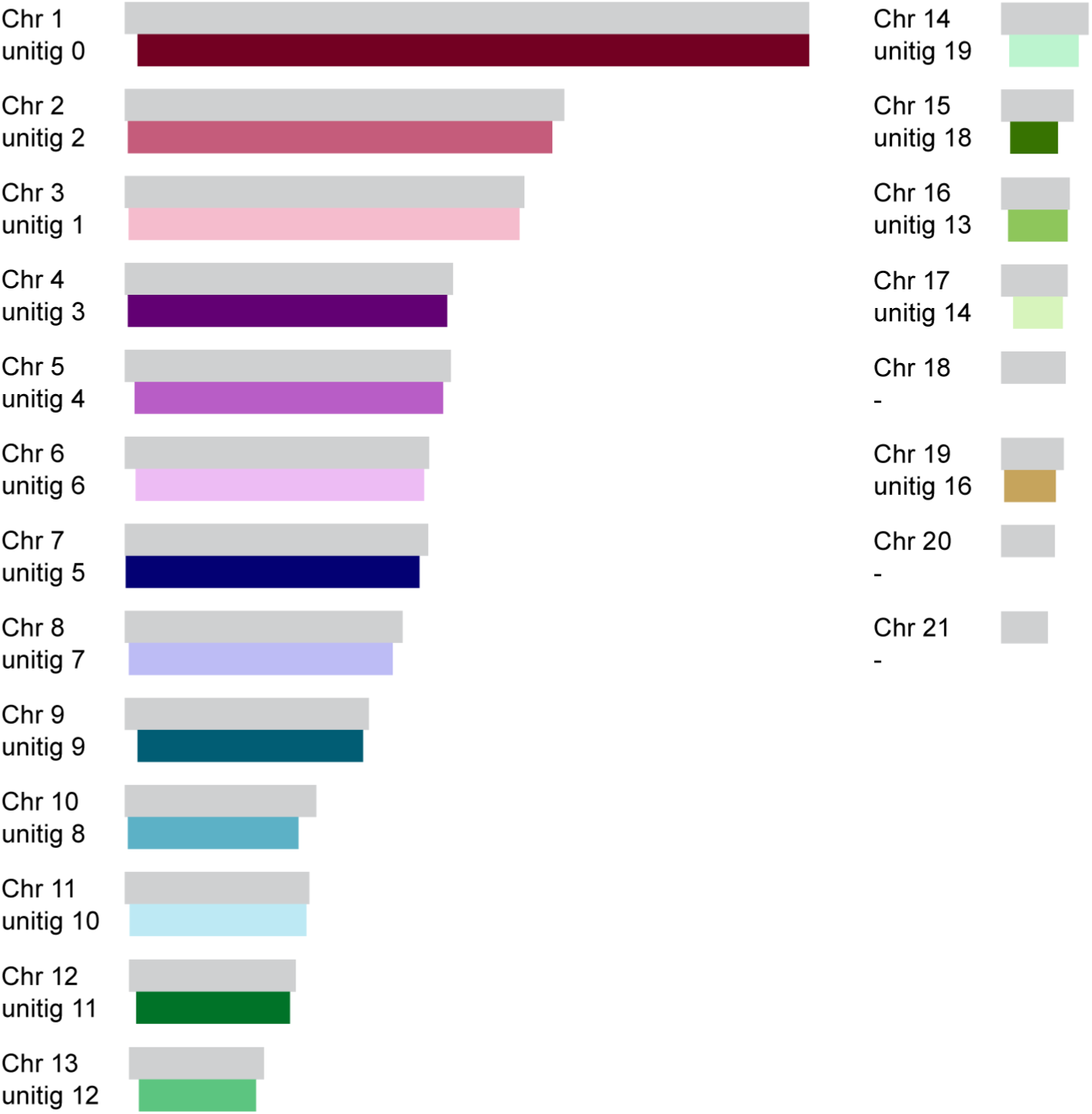
Synteny between Zt10 *de novo* assembled unitigs and IPO323 chromosomes.

**S7 Figure.**
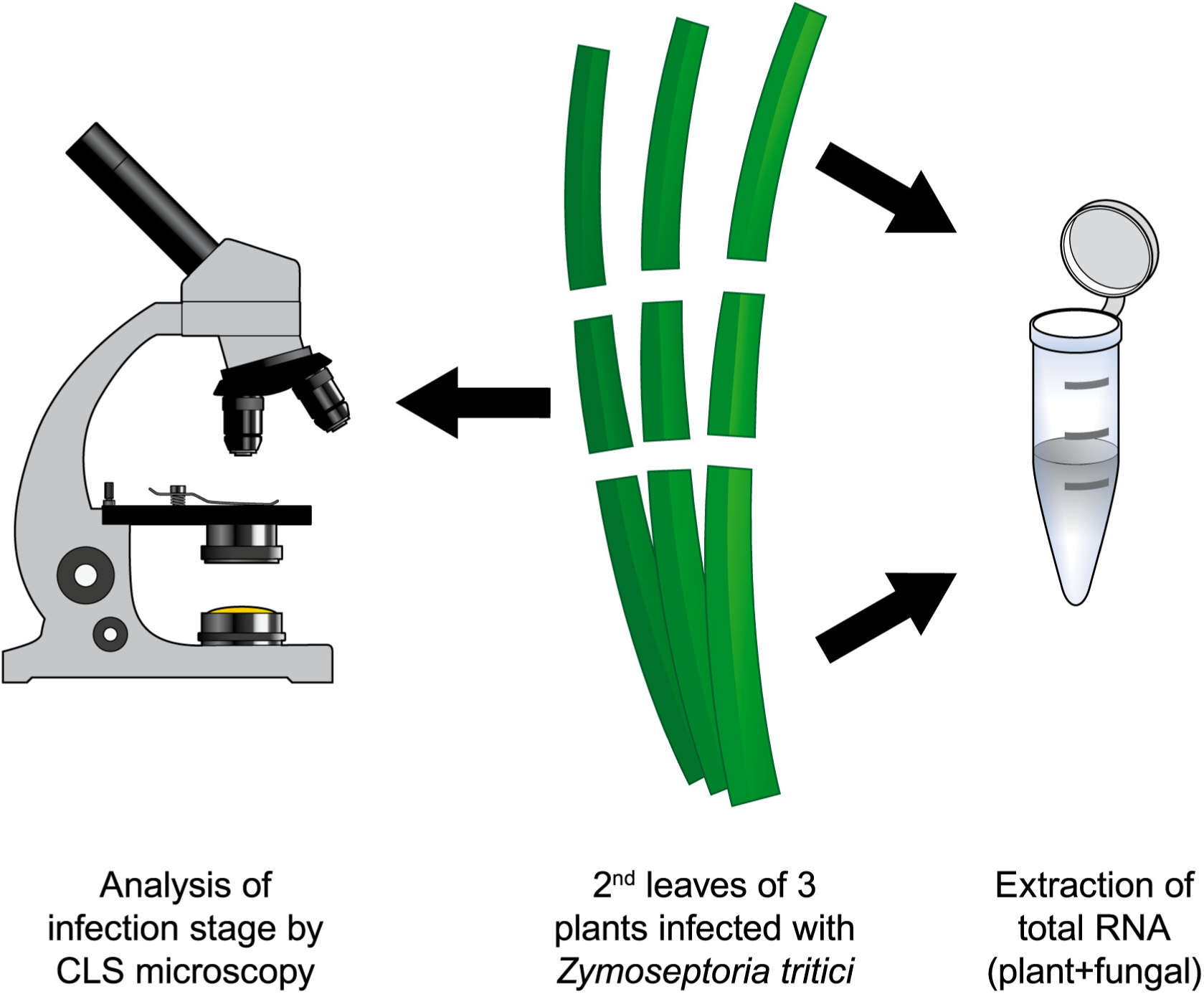
Generation of isolate- and stage-specific transcriptomes was enabled by confocal microscopy analyses. The schematic drawing illustrates how we selected samples for RNA-seq. Central sections of *Z.* tritici-infected wheat leaves from three independent plants (second leaf of each plant) were stained and analyzed by confocal laser-scanning microscopy while the remaining infected leaf material was pooled and ground in liquid nitrogen for total RNA extraction. RNA samples subjected to sequencing were chosen based on the morphological infection stage that we observed in the central leaf section by microscopy.

**S8 Figure.**
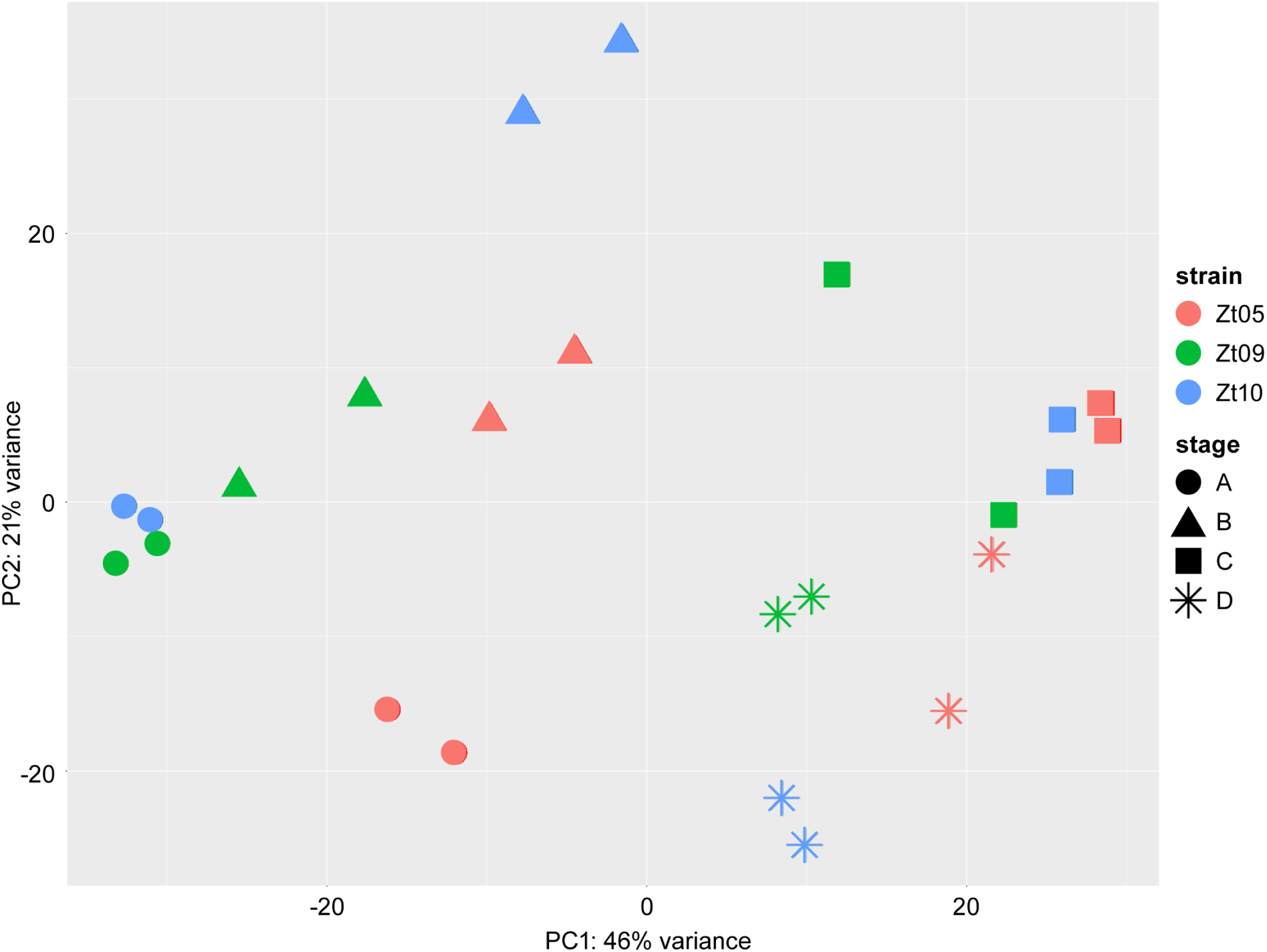
RNA-seq data principal component analysis plot based on rlog-transformed read counts for *Z. tritici* core genes. PC1 separates datasets from infections stages A and B from stages C and D. Stage-specific datasets from all isolates cluster together.

**S9 Figure.**
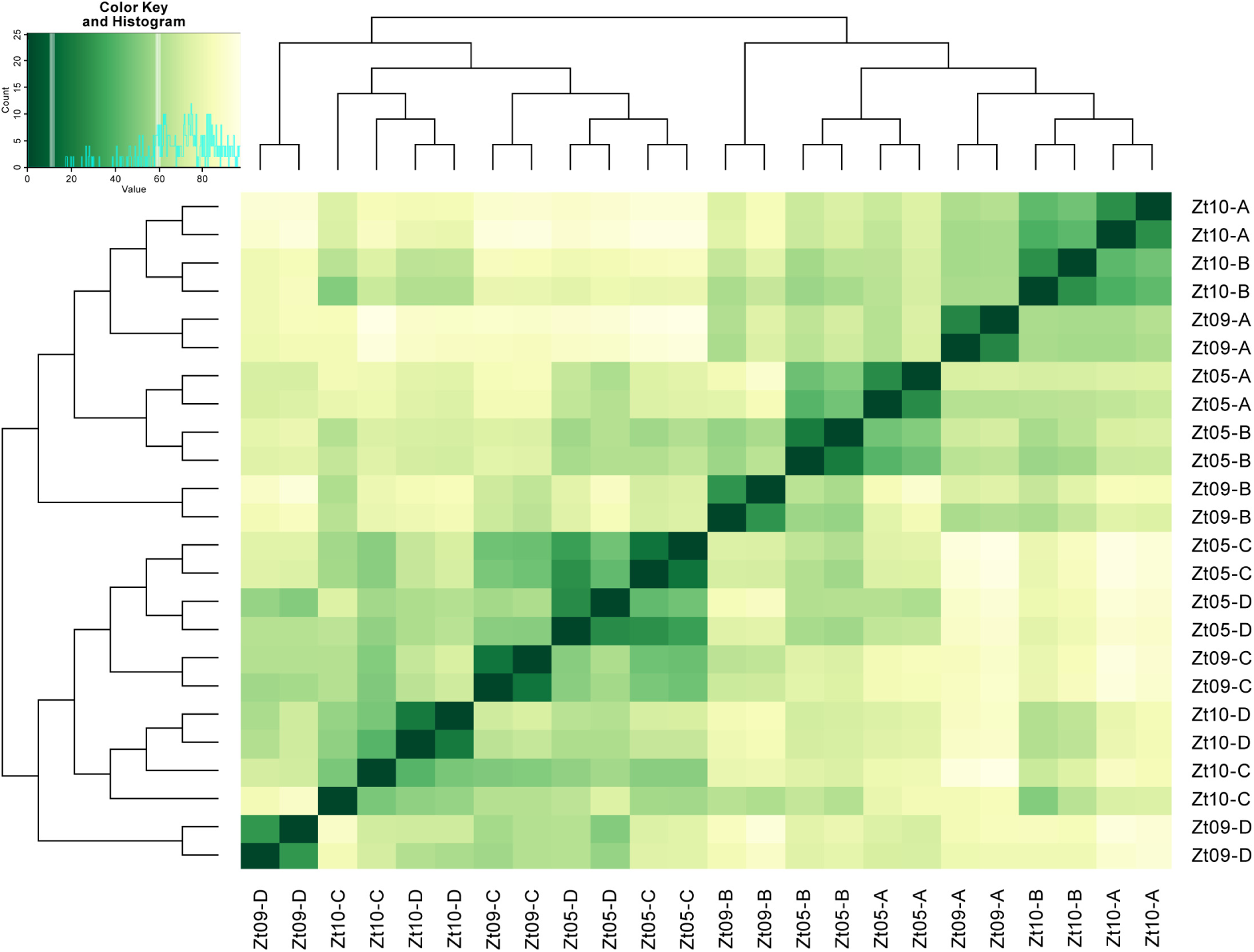
Transcriptome data distance matrix based on rlog-transformed read counts for *Z. tritici* core genes. Datasets from stages A and B representing biotrophic growth form one cluster as do datasets of stages C and D representing necrotrophic growth.

**S10 Figure.**
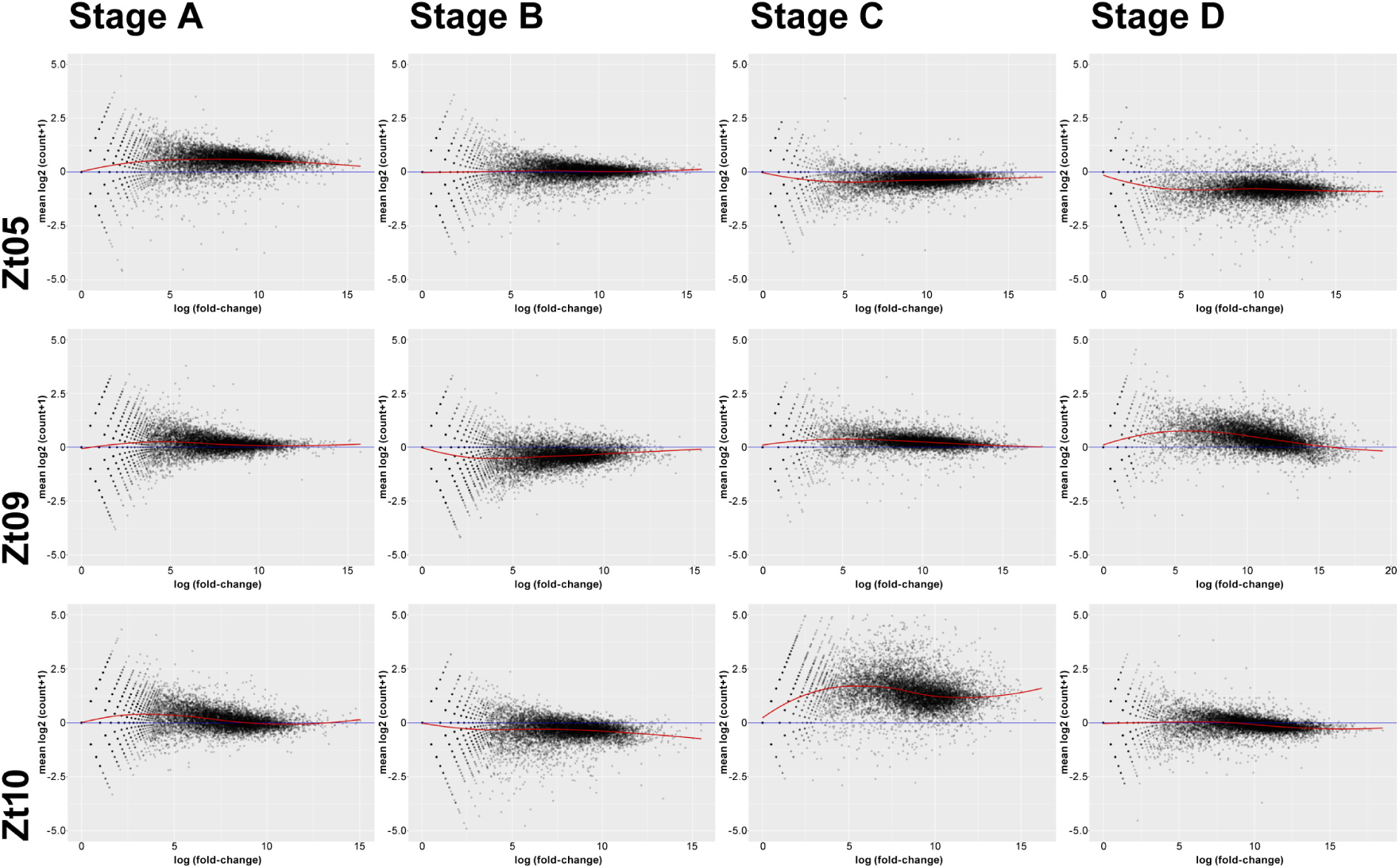
MA plots comparing replicates for each RNA-seq dataset. Pairwise comparisons of replicates for each wheat infection stage of each *Z. tritici* isolate without normalization. x-axis: mean log_2_ (read count per gene+1), y-axis: log (fold-change). The greatest variation among replicates was between the Zt10 stage C datasets.

**S11 Figure.**
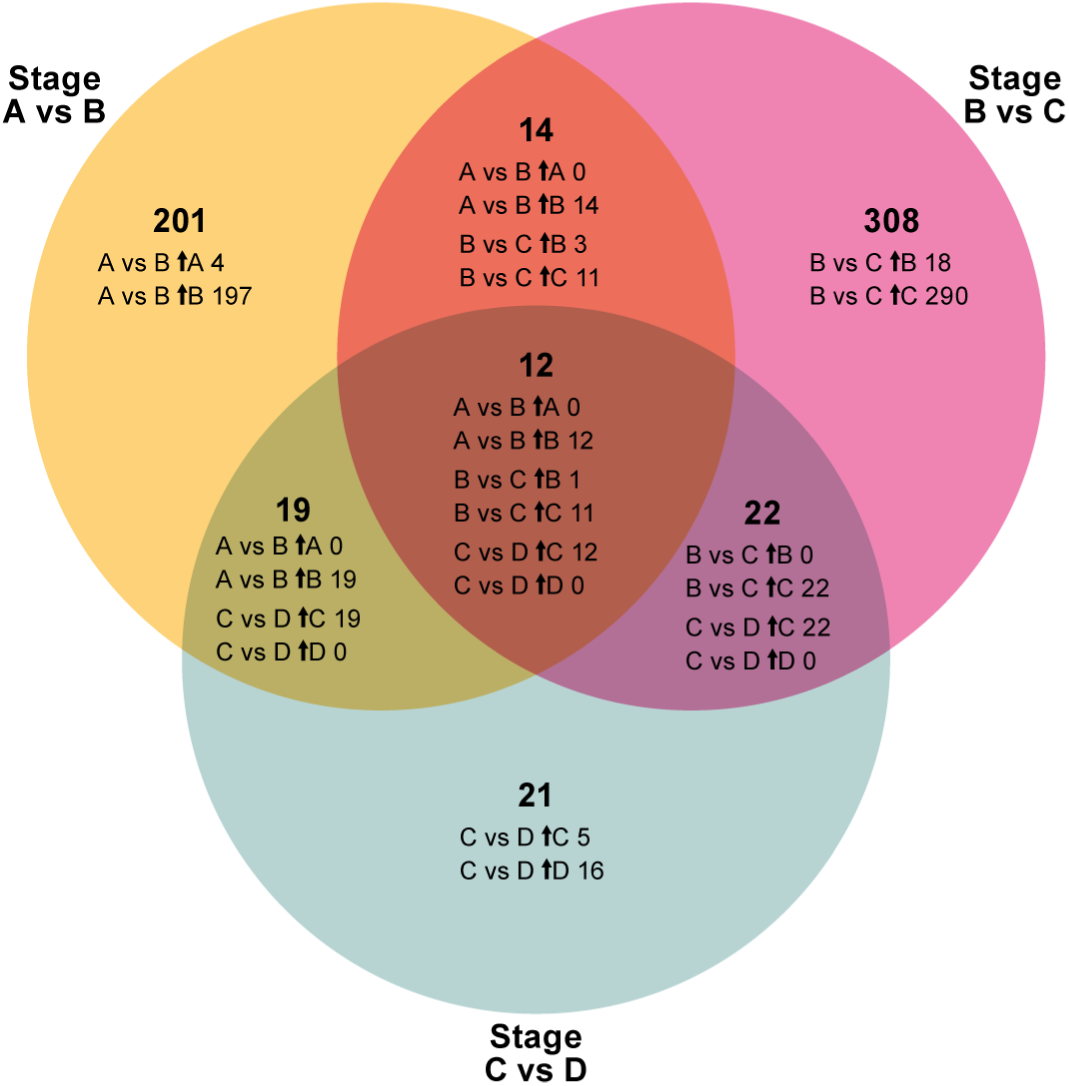
597 genes are differentially expressed between the infection stages in all three isolates, and 79 genes are differentially expressed between more than two stages. The Venn diagram illustrates how genes that are differentially expressed between *Z. tritici* infection stages are shared between stage comparisons. Differential expression analyses were performed with DESeq2. Differentially expressed genes have *P*_adj_ ≤ 0.01 and an absolute log_2_ fold change between infection stages of ≥2. Small arrows (℩) indicate the stage in which genes are significantly up-regulated.

**S12 Figure 1-3.**
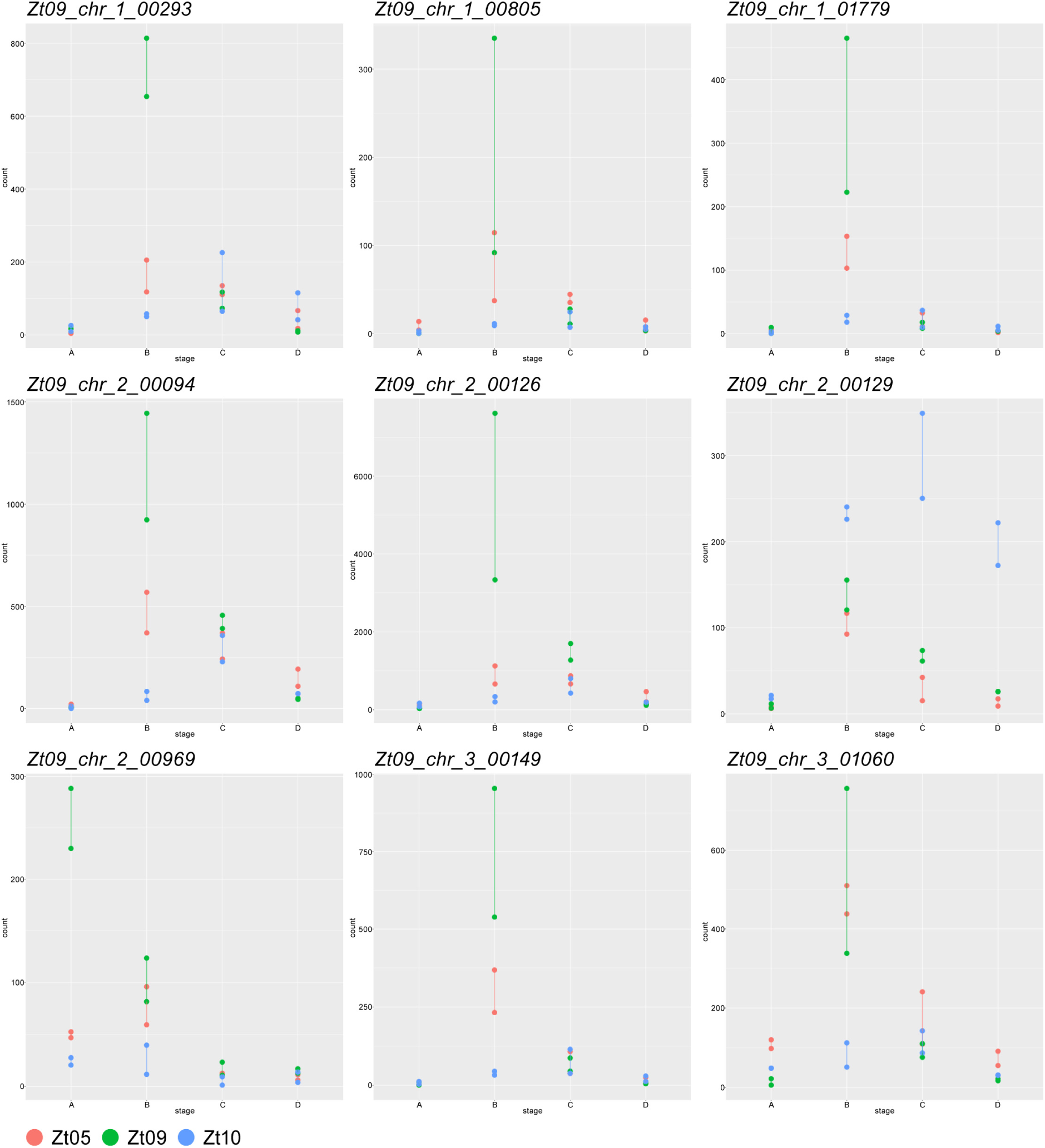
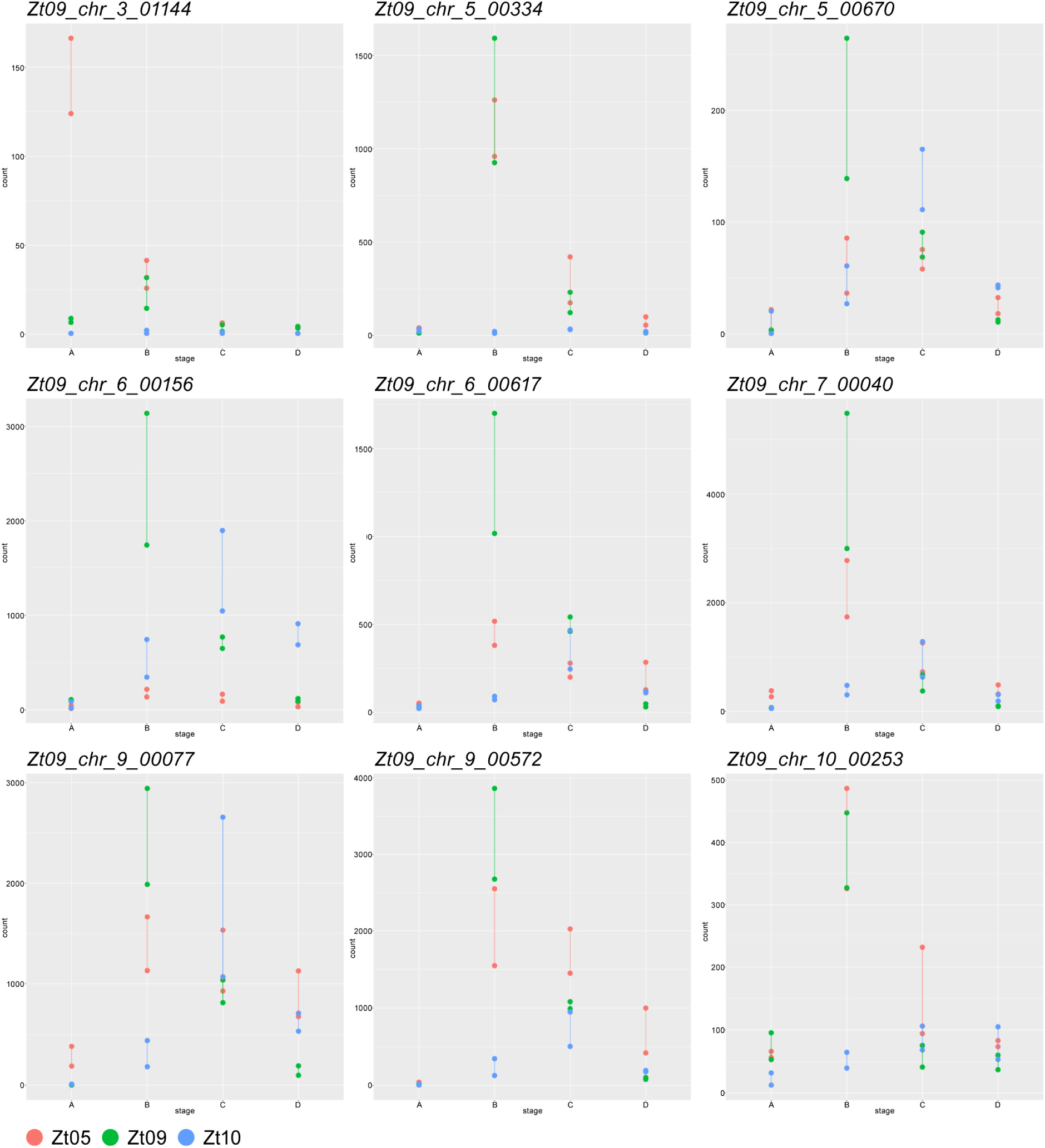
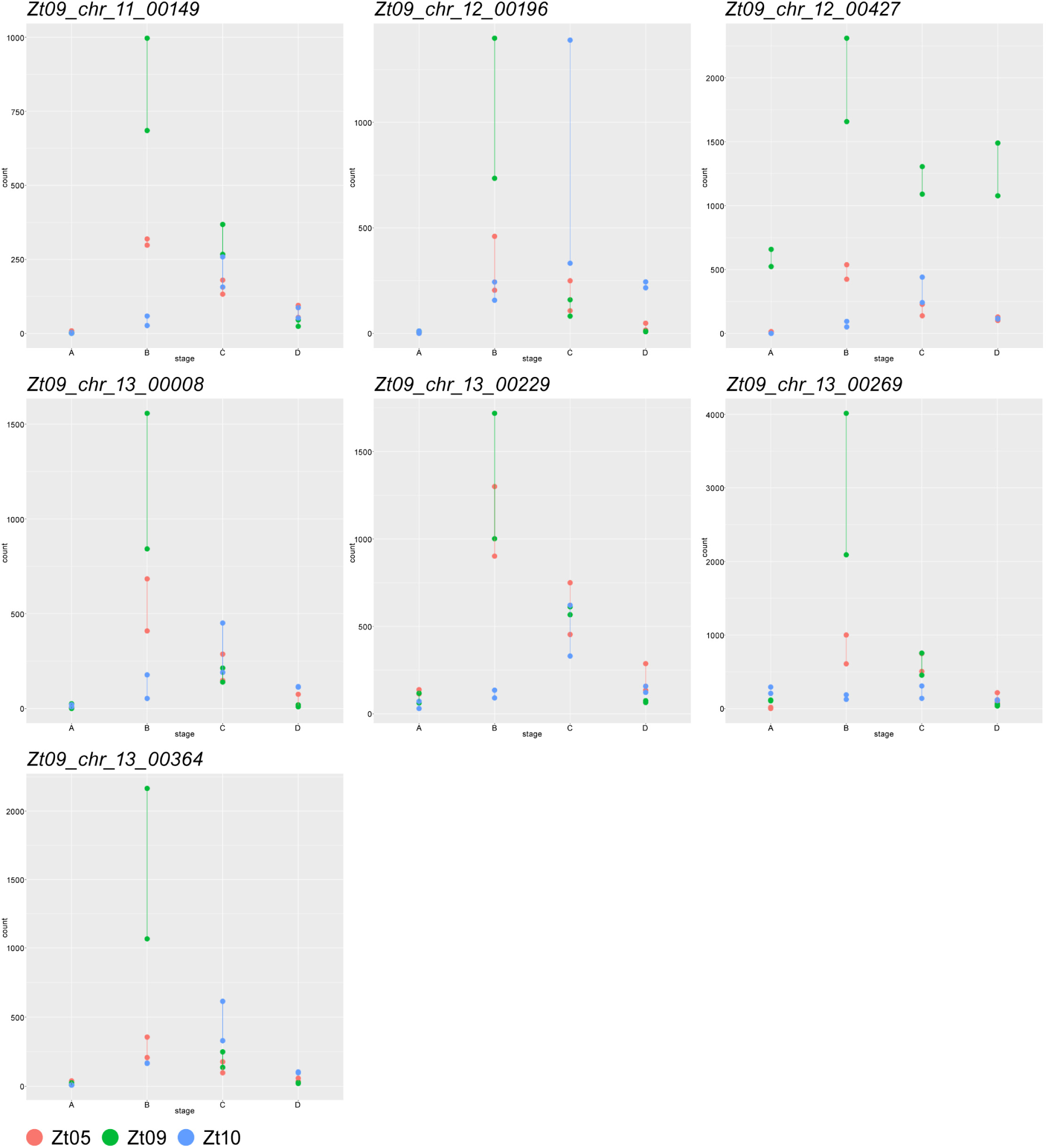
Expression profiles of core *Z. tritici* biotrophic effector candidates based on normalized read counts per gene. Read counts were normalized across the four core infection stages (A to D) and the three *Z. tritici* isolates Zt05, Zt09, and Zt10 and represent a measure of relative gene expression between infection stages and between isolates.

**S13 Figure 1-4.**
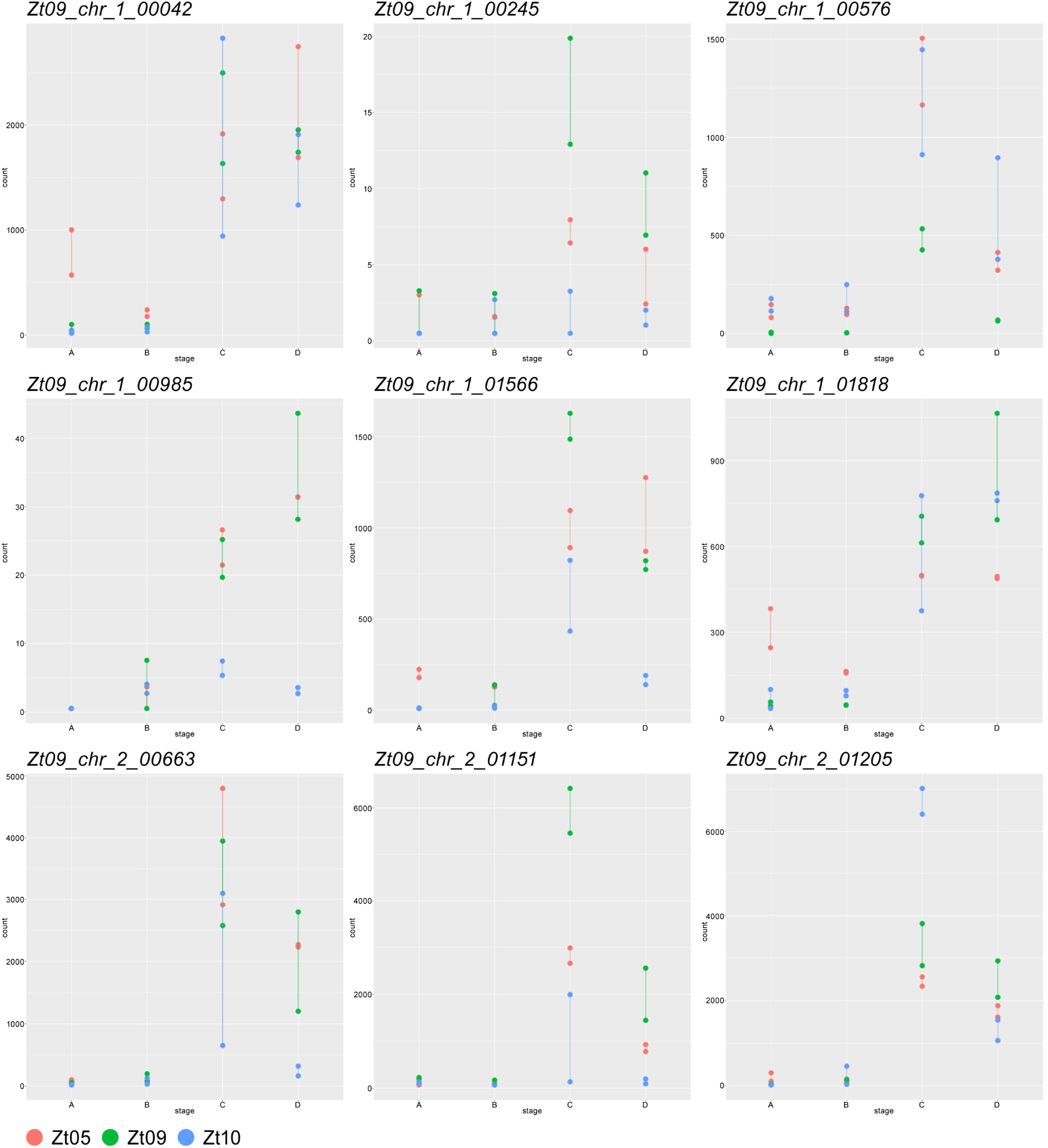
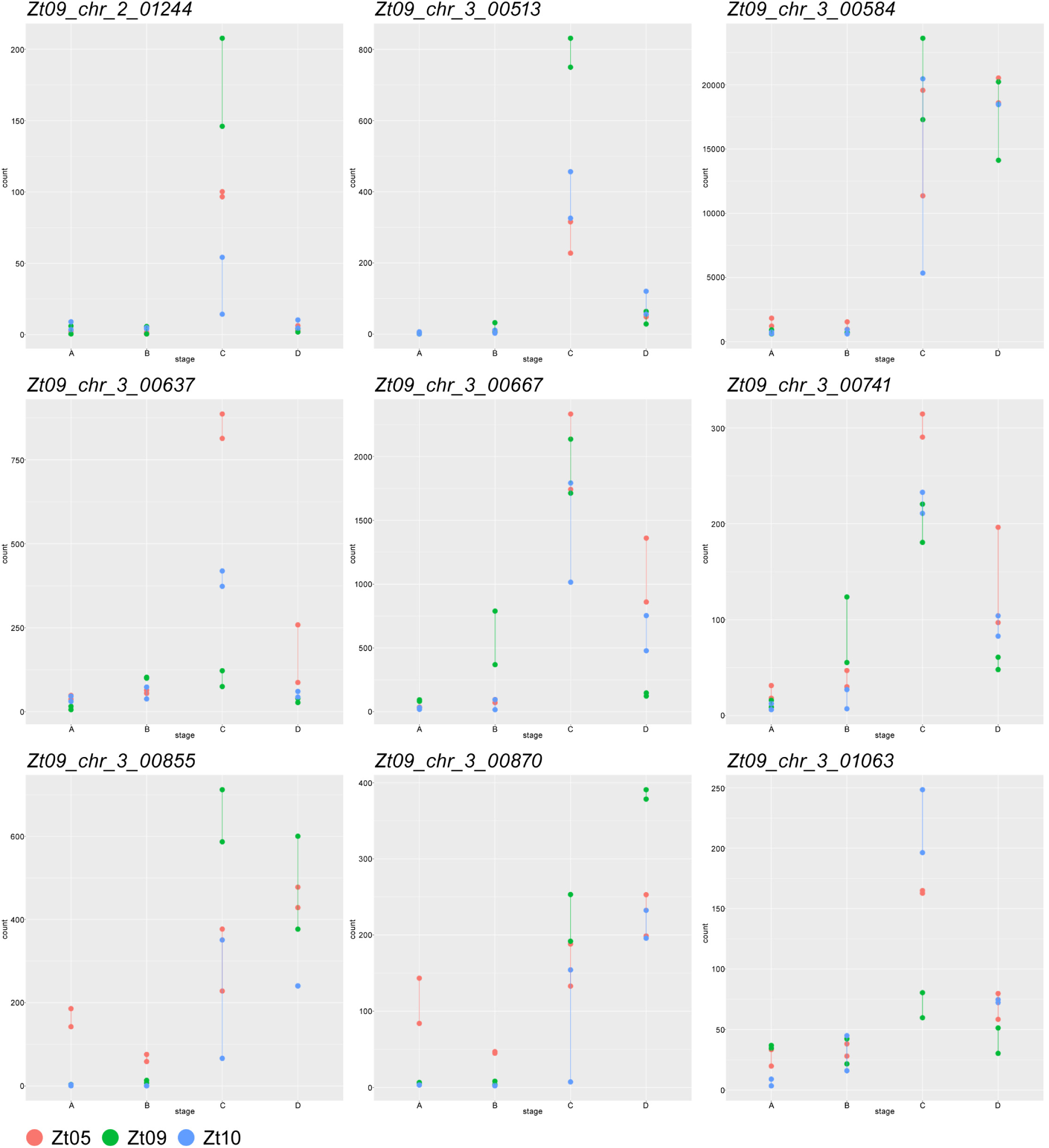
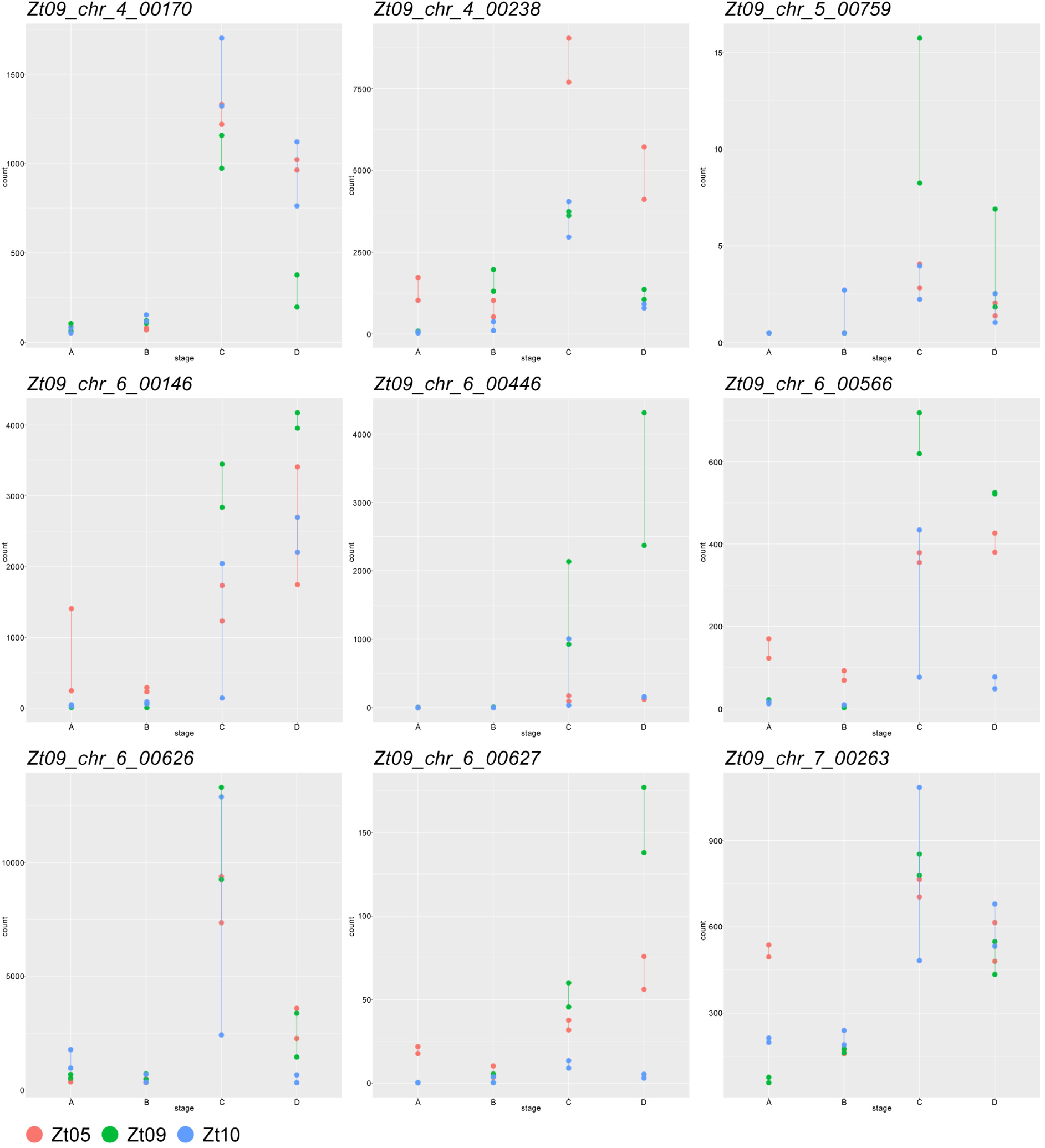
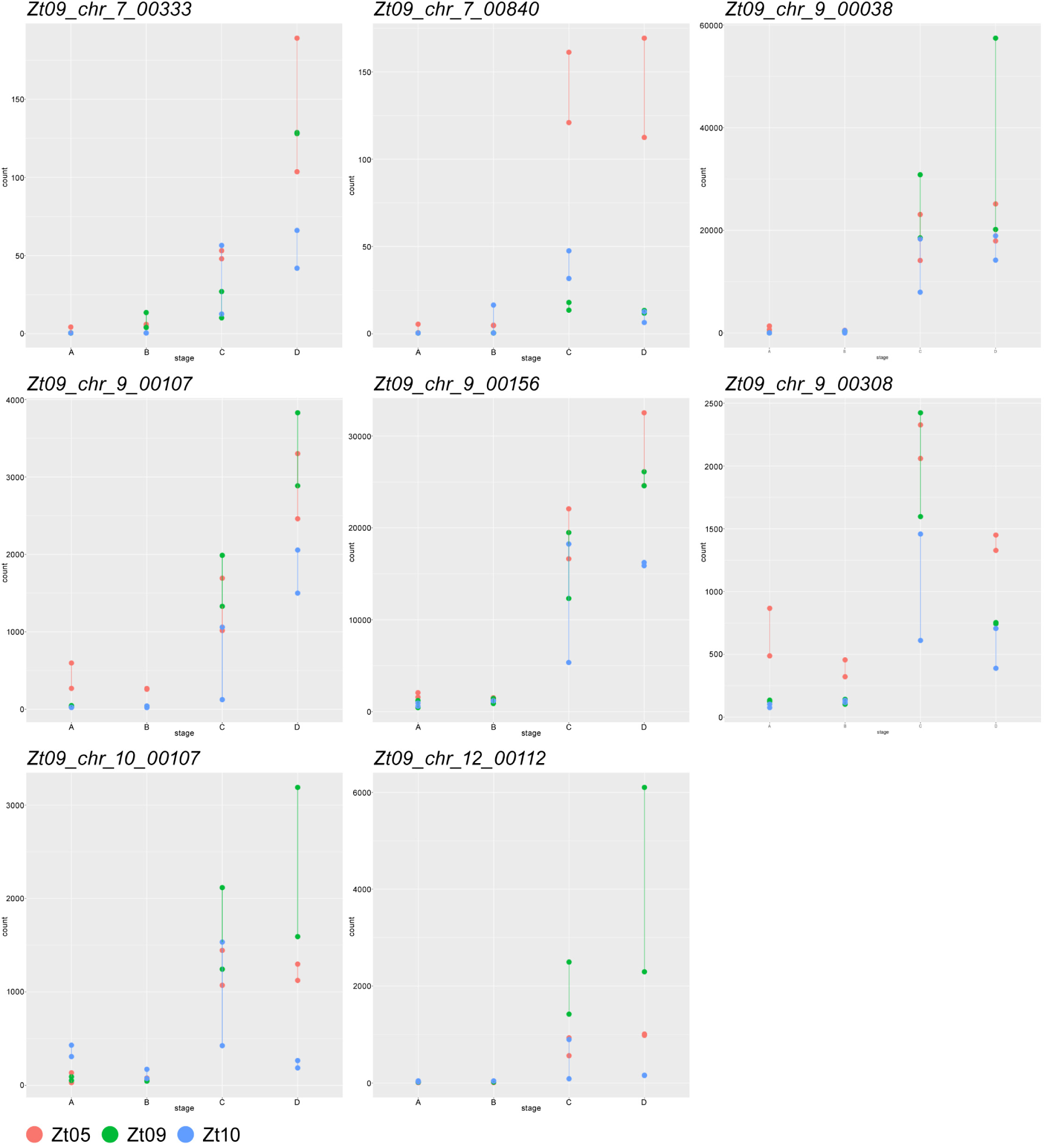
Expression profiles of *Z. tritici* core necrotrophic effector candidates based on normalized read counts per gene. Read counts were normalized across the four core infection stages (A to D) and the three *Z. tritici* isolates Zt05, Zt09, and Zt10 and represent a measure of relative gene expression between infection stages and between isolates.

**S14 Figure.**
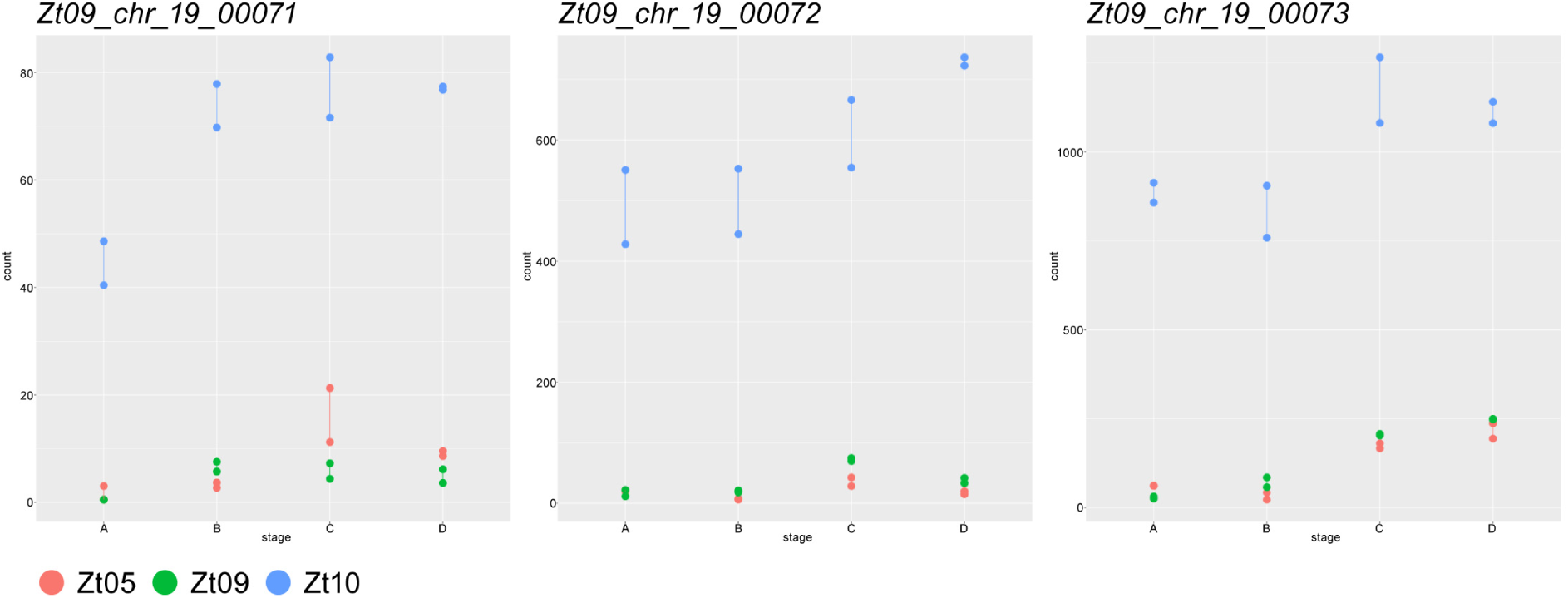
Expression profiles of three *Z. tritici* genes located on accessory chromosome 19 in Zt09. The neighboring genes *Zt09_chr_19_00071, Zt09_chr_19_00072,* and *Zt09_chr_19_00073* are significantly higher expressed in Zt10 during all four infection stages. Read counts were normalized across the four core infection stages (A to D) and the three *Z. tritici* isolates (Zt05, Zt09, and Zt10) and represent a measure of relative gene expression between infection stages and between isolates.

**S15 Figure.**
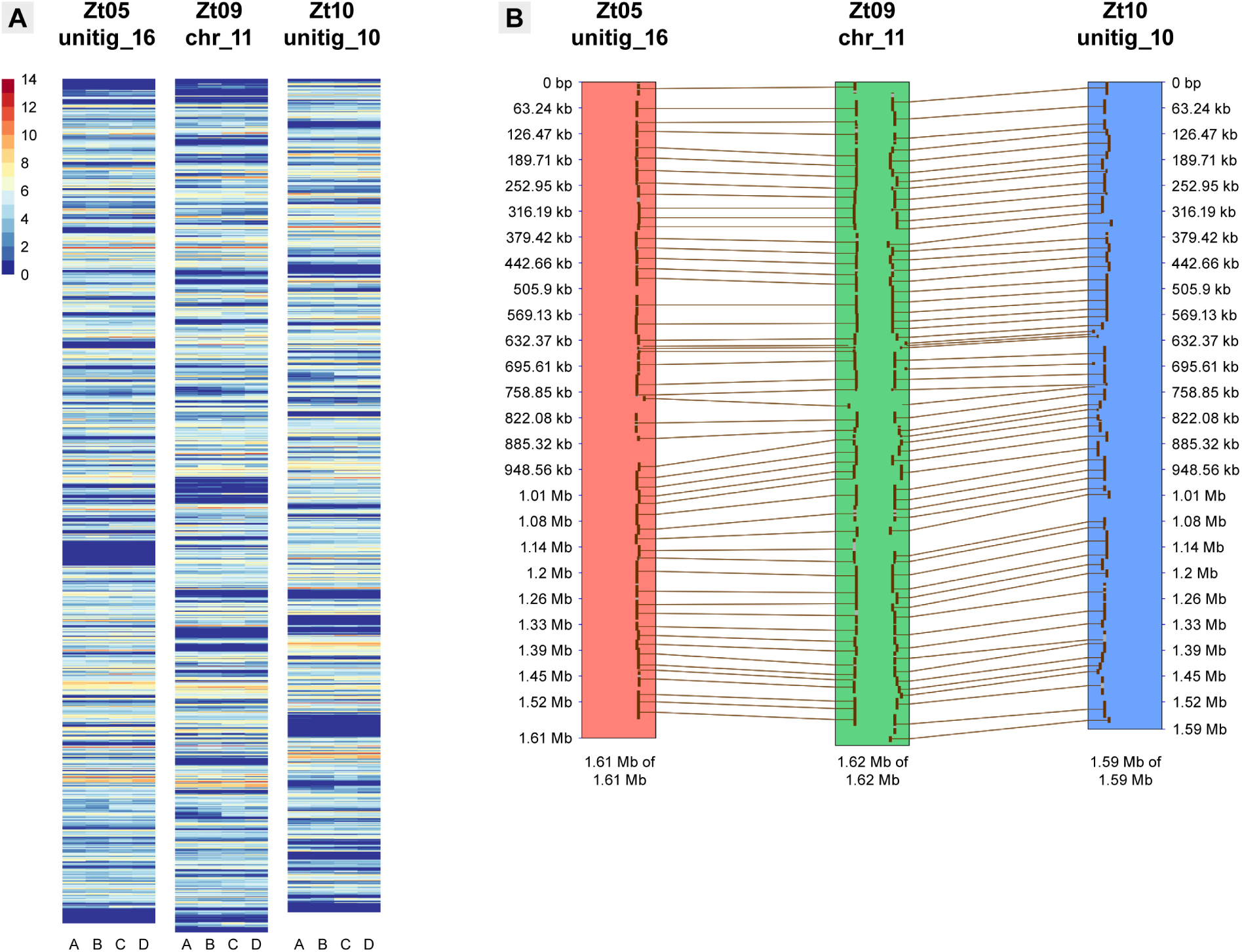
Distribution of transcriptionally active loci on core chromosome 11. (A) Heatmaps of log_2_-transformed FPKM expression values for 1-kb windows along IP0323/Zt09 core chromosome 11 and unitigs 16 and 10 in Zt05 and Zt10 for the four wheat infection stages. (B) Synteny plot comparing IP0323/Zt09 chromosome 11 and unitigs 16 and 10 in Zt05 and Zt10.

**S16 Figure.**
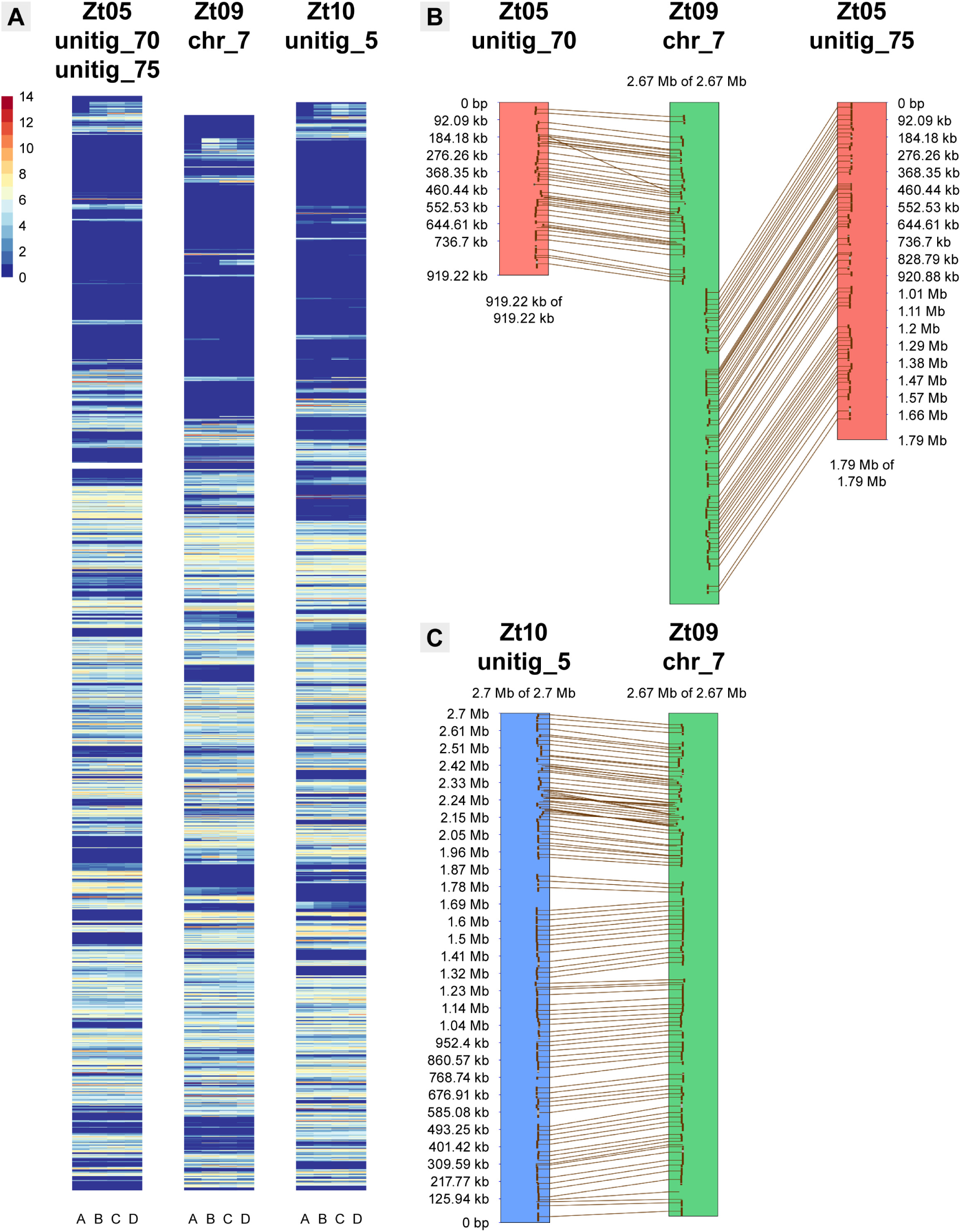
Distribution of transcriptionally active loci on core chromosome 7. **(A)** Heatmaps of log_2_-transformed FPKM expression values for 1-kb windows along IPO323/Zt09 core chromosome 7 and unitig 5 in Zt10 and unitigs 70 and 75 in Zt05 for the four wheat infection stages. **(B)** Synteny plots comparing IPO323/Zt09 chromosome 7 and unitigs 70 and 75 of Zt05 and **(C)** unitig 5 of Zt10.

**S17 Figure.**
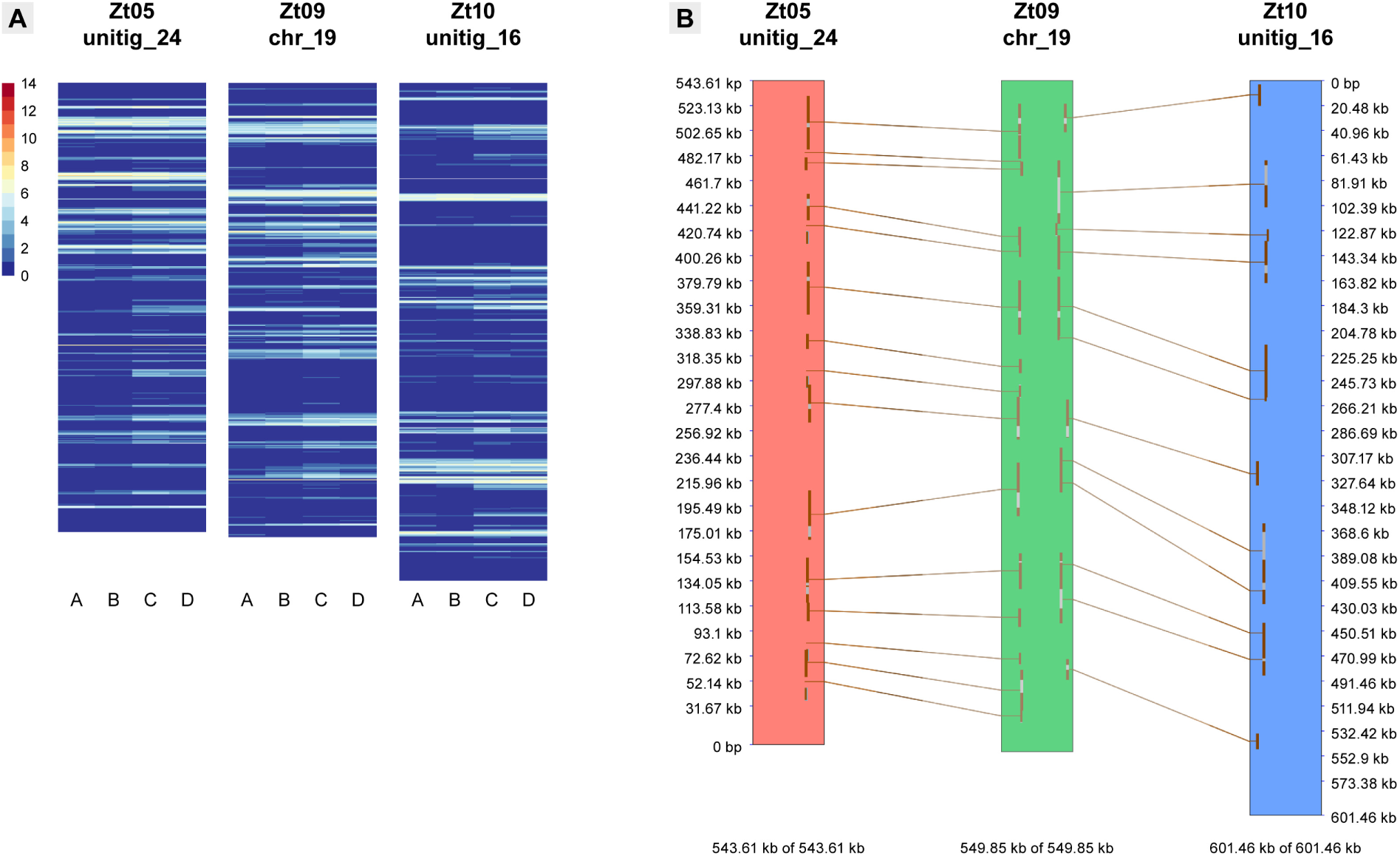
Distribution of transcriptionally active loci on accessory chromosome 19. **(A)** Heatmaps of log_2_-transformed FPKM expression values for 1-kb windows along IPÜ323/Zt09 accessory chromosome 19 and the syntenic unitigs 24 and 16 in Zt05 and Zt10 for the four wheat infection stages. **(B)** Synteny plot comparing IPO323/Zt09 chromosome 19 and unitigs 24 and 16 of Zt05 and Zt10, respectively.

**S1 Animation. *Z. tritici* initial wheat infection stage.**

Tomographic animation of confocal image z-stack showing infection hypha of *Z. tritici* isolate Zt09 entering wheat leaf tissue by open stoma at 4 dpi. The hypha grows closely attached to stomatal guard cell. Nuclei and wheat cells are displayed in *purple* and fungal structures in *green.* Reference transmitted images are in *grey.* Scale bar = 25 μm.

**S2 Animation. *Z. tritici* initial wheat infection stage.**

Tomographic animation of confocal image z-stack showing epiphyllous proliferation, infecting hyphae, and hyphal growth inside wheat sub-stomatal cavity and mesophyll of *Z. tritici* isolate Zt05 at 3 dpi. Nuclei and wheat cells are displayed in *purple* and fungal structures in *green.* Reference transmitted images are in *grey.* Scale bar = 25 μm.

**S3 Animation. Biotrophic colonization of wheat mesophyll by *Z. tritici* Zt05.**

Tomographic animation of confocal image z-stack showing epiphyllous hyphae as well as the dense biotrophic intercellular hyphal network of *Z. tritici* isolate Zt05 inside wheat mesophyll at 7 dpi. Long, straight hyphae grow in the interspace of wheat epidermis and mesophyll cells. Hyphae grow in close contact to plant cells. Nuclei and wheat cells are displayed in *purple* and fungal structures in *green.* Scale bar = 50 μm.

**S4 Animation. Biotrophic colonization of wheat mesophyll by *Z. tritici* Zt09.**

Tomographic animation of confocal image z-stack showing biotrophic intercellular hyphae of *Z. tritici* isolate Zt09 inside wheat leaf tissue at 11 dpi. Nuclei and wheat cells are displayed in *purple* and fungal structures in *green.* Reference transmitted images are in *grey.* Scale bar = 25 μm.

**S5 Animation. *Z. tritici* Zt05 pycnidium development.**

Tomographic animation of confocal image z-stack showing the development of primal structures of *Z. tritici* Zt05 pycnidium in the wheat sub-stomatal cavity during the early lifestyle transition stage at 11 dpi. Nuclei and wheat cells are displayed in *purple* and fungal structures in *green.* Reference transmitted images are in *grey.* Scale bar = 25 μm.

**S6 Animation. *Z. tritici* Zt09 pycnidium development.**

Tomographic animation of confocal image z-stack showing the development of primal structures of *Z. tritici* Zt09 pycnidium in the wheat sub-stomatal cavity during the lifestyle transition stage at 13 dpi. Nuclei and wheat cells are displayed in *purple* and fungal structures in *green.* Reference transmitted images are in *grey.* Scale bar = 25 μm.

**S7 Animation. Development of two pycnidium initials of *Z. tritici* Zt10.**

Tomographic animation of confocal image z-stack showing *Z. tritici* Zt10 pycnidium in the wheat sub-stomatal cavity developing from two initial stromata during the early lifestyle transition stage at 13 dpi. Nuclei and wheat cells are displayed in *purple* and fungal structures in *green.* The fluorescence of *Z. tritici* hyphae inside plant tissue is very weak. *Purple* fungal nuclei are mainly visible. Reference transmitted images are in *grey.* Scale bar = 25 μm.

**S8 Animation. Mature pycnidia of *Z. tritici* Zt05.**

Tomographic animation of confocal image z-stack showing asexual pycnidia of *Z. tritici* isolate Zt05 with pycnidiospores during necrotrophic infection stage at 21 dpi. The intercellular space of wheat mesophyll is densely colonized by *Z. tritici* hyphae. Nuclei and wheat cells are displayed in *purple* and fungal structures in *green.* The fluorescence of *Z. tritici* hyphae inside the plant tissue is weak. *Purple* fungal nuclei are mainly visible. Reference transmitted images are in *grey.* Scale bar = 50 μm.

**S9 Animation. Pycnidium of *Z. tritici* Zt09.**

Tomographic animation of confocal image z-stack showing asexual pycnidium of *Z. tritici* isolate Zt09 at 20 dpi. Hyphae grow in close contact to collapsing wheat mesophyll cells. Nuclei and wheat cells are in *purple* and fungal structures in *green.* Reference transmitted images are in *grey.* Scale bar = 25 μm.

## Supporting information - S1 Text

### Supplementary Results

#### Comparative analysis of Z. *tritici* infection development by confocal microscopy

We set out to characterize the infection development of the three *Z. tritici* isolates Zt05, Zt09 and Zt10 on the surface as well as within wheat leaves. To this end, we conducted a detailed survey where we analyzed leaf material harvested at 3-5, 7, 8, 10-14, 17, 19-21, 25, and 28 days after inoculation (dpi) by confocal laser-scanning microscopy. We used large z-stacks of longitudinal optical sections to reconstruct the spatial and temporal fungal colonization outside and within infected tissue. We first focused on shared characteristics of the three *Z. tritici* isolates during host colonization and reproduction. Thereby, as described in details below, we identified four distinct infection stages that we define as the core *Z. tritici* infection program (Fig 3). Furthermore, we characterized the differences and isolate-specific aspects of infection development of the three isolates including temporal, spatial, and quantitative variation of host colonization.

#### The shared core infection program of *Z. tritici* is characterized by four infection stages

The first infection stage A is the penetration of wheat leaf tissue by *Z. tritici* hyphae. Germination of fungal cells on the leaf surface is initiated and developing infection hyphae enter wheat stomata. We observed that germ tubes emerge from *Z. tritici* cells at different time points post inoculation indicating that the fungi sense and respond to particular host-derived cues that trigger the developmental switch from spores to hyphal growth [1]. Germ tubes develop into filaments of which some grow directed towards stomatal openings and enter the leaf (Fig 3A, stage A, S1 and S2 Animation). Occasionally, we noticed slight, spatially restricted swelling of hyphae on top of stomata that resemble primitive appressoria as also previously reported [2,3]. However, we never observed a direct penetration of epidermal cells. During stomatal passage and in the sub-stomatal cavities, *Z. tritici* infection hyphae grow in tight contact to the wheat guard cells. The close physical contact between hyphae and plant cells might facilitate delivery of *Z. tritici* effector molecules [4] or serve as a structural scaffold to direct fungal hyphae in the host tissue [5]. However, not all inoculated *Z. tritici* cells caused stomatal penetrations. We found that a portion of cells did not form germ tubes within 28 dpi and that development of filaments was stopped before entering stomata; what we also expect to happen in field infections.

The subsequent infection stage B is characterized by biotrophic growth of *Z. tritici* and the symptomless colonization of wheat mesophyll (Fig 3A, stage B, S3 and S4 Animation). For successful infections, the pathogen must avoid recognition by the host immune system and/or suppress activation of defense responses during biotrophic growth. We observed strict intercellular hyphal colonization, starting from sub-stomatal cavities into adjacent mesophyll tissue, whereat the hyphae grow in close contact with host cells. Remarkably, hyphae first grow in the interspace of epidermis and first mesophyll layer. There, hyphae spread in the grooves between adjacent epidermal cells and only subsequently explore subjacent mesophyll cell layers. The transition from symptomless biotrophic to necrotrophic colonization and the development of disease symptoms like chlorosis and necrotic lesions represent the third infection stage C (Fig 3A, stage C). From there on, *Z. tritici* colonizes a biochemically changing host environment (Fig 2B) and feeds on nutrients released by the host cell death to build pycndia. Hyphae are branching and grow in all mesophyll layers, surrounding individual wheat mesophyll cells. Simultaneously, primal structures of the asexual fruiting bodies, the pycnidia, are established and begin to develop. Hyphae form ring-like scaffolds in the sub-stomatal cavities where hyphae align and build stromata (S5-S7 Animation) that later give rise to conidiogenous cells.

The last stage D concludes the infection and is characterized by necrotrophic colonization and asexual reproduction (Fig 3A, stage D). *Z. tritici* hyphae eventually grow in an environment that is very nutrient rich and attractive to other microbial competitors, but also putatively toxic e.g. due to high concentrations of reactive oxygen species (Fig 2B) In necrotic leaf regions, the dead mesophyll tissue is heavily colonized and the asexual fruiting bodies are visible and maturated (S8 and S9 Animation). Hyphae wrap around dead, collapsed mesophyll cells several times. The pathogen may keep this tight contact to the degrading plant cells to increase the acquisition of nutrients and maybe also to protect them from competing saprotrophic species. Sub-stomatal cavities within the colonized leaf areas are occupied by sub-globose pycnidia that can grow into the adjacent mesophyll tissue. Mature pycnidia harbour hyaline, oblong asexual pycnidiospores that are released through the former stomatal opening.

In general, the described infection stages of *Z. tritici* can be well distinguished by considering the majority of all infection events within inoculated leaf regions. However, we also observed that different infections stages are present simultaneously within one leaf. Infections by individual *Z. tritici* cells occur within a temporal range after inoculation and are not fully synchronized. Moreover, environmental influences and host physiological processes act differently on individual leaves and plants which also can lead to the temporal variation in infection development of *Z. tritici*.

#### Highly differentiated infection phenotypes of the three *Z. tritici* isolates on Obelisk wheat

Although we clearly recognize the four core infection stages for the three *Z. tritici* isolates, we found that the infection phenotypes of Zt05, Zt09, and Zt10 are highly differentiated. We observed temporal, spatial and quantitative variation in the infection development of these isolates on the wheat cultivar Obelisk.

The duration of the initial infection stage A—in particular the period between inoculation and stomatal penetrations—is different in the three *Z. tritici* isolates. For Zt05, infection hyphae enter stomata within 5 dpi. Germ-tube formation and stomatal penetration is usually slower for Zt09 (up to 8 dpi) and most delayed for Zt10 (up to 10 dpi) (Fig 3A, stage A). We also noticed strong epiphyllous proliferation and mycelium formation for Zt05 during all infection stages and frequently, several infection hyphae of Zt05 enter one stoma (Fig 3A, stage A: Zt05). In general, hyphae of Zt05 and Z09 penetrate stomata at high frequencies, while we saw fewer stomatal penetrations for the Zt10 leading to patchy infections within the inoculated leaf areas (Fig 3A, stage C and D: Zt10).

The extent of biotrophic colonization during infection stage B comprises the most pronounced difference between the three isolates. Zt05 builds biotrophic hyphal networks in the mesophyll tissue with long “runner” hyphae growing primarily longitudinally between epidermis and mesophyll (Fig 3A, stage B: Zt05, Fig 3B.1, S3 Animation). Biotrophic hyphal networks of Zt09 are smaller and located mainly in the interspace of epidermis and mesophyll as well as between the cells of the upper mesophyll layer (Fig 3A, stage B: Zt09, Fig 3B.2). Biotrophic colonization by Zt10, however, is very poor and hyphal growth is limited to the mesophyll cells adjacent to sub-stomatal cavities (Fig 3A, stage B: Zt10, Fig 3B.3). Since biotrophic colonization depends on successful evasion of host immunity [6], the different extent of colonization could reflect different strategies to bypass recognition in a given host genotype.

During the later infection stages, differences between the isolates are smaller and primarily relate to temporal variation. Transition to necrotrophic growth usually first occurs for Zt05 (9 to 14 dpi), followed by Zt09 (13 to 16 dpi), and Zt10 (13 and 17 dpi) (Fig 3A, stage C). Studying the development of the asexual fruiting bodies, we frequently noticed the formation of two pycnidia in one sub-stomatal cavity for Zt10 (Fig 3A, stages C and D: Zt10, S7 Animation). This was observed less often for the other two isolates. The onset of infection stage D occurs in the same temporal order as for stage C, first for Zt05, followed by Zt09, and last by Zt10. At 28 dpi, inoculated leaf areas are usually fully necrotic for Zt09 and frequently covered by several distinct necrotic lesions for Zt10 (S1 Fig).

Taken together, we observed highly differentiated infection phenotypes for the three *Z. tritici* isolates due to isolate-specific infection development. However, the final production of asexual pycnidia did not differ significantly between the three isolates (Fig 1), suggesting that the isolate-specific aspects in host-pathogen interaction sum up to equally good strategies for host colonization and asexual reproduction.

With several independent plant infection experiments using the three isolates Zt05, Zt09, and Zt10 on the wheat cultivar Obelisk, we found that the temporal disease progress and, consequently, the duration of the different infection can stages vary between experiments. However, although the precise timing for the onset of the four stages can differ between experiments, the relative temporal differences between the isolates, as described above, remain consistent.

#### Karyotypes and synteny analyses of the three *Z. tritici* isolates

Putatively dispensable chromosomes in the size range of 225 to 1,125 kb were separated by pulsed-field gel electrophoresis (PFGE) and visualized for the three *Z. tritici* isolates (S4 Fig). The previously reported loss of chromosome 18 (~574 kb) in Zt09 [7] could not be demonstrated by PFGE, as the chromosome could not be separated from the chromosomes 17 (~584 kb) and 16 (~607 kb) with almost the same size.

We observed intense chromosomal bands around 540 kb and 710 kb in Zt05 and around 615 kb in Zt10 (S4 Fig). This indicates that both isolates possess additional chromosomes to the seven (Zt05) and four (Zt10) chromosomes that we identified based on separated chromosomal bands by PFGE. Indeed, analyses of *de novo* genome assemblies based on long-read SMRT Sequencing data for Zt05 and Zt10, show eight and five mainly full chromosome unitigs (indicated by telomeric repeats at both ends) in the size range of 290 to 905 kb (S2 Table) with synteny to IPO323/Zt09 chromosomes 14 to 21 (S5 and S6 Fig).

On the PFGE gel, we identified a chromosomal band around 640 kb for Zt10 that possibly represents unitig 15 (634 kb). This unitig shares no synteny with an IPO323 chromosome suggesting that this is a hitherto not described accessory chromosome in the species. However, as there is only one telomeric repeat present at one end of unitig 15 the assembly does not represent the full chromosome. Moreover, transcribed regions on unitig 15 are syntenic to a larger block on Zt05 unitig 20 that was identified as homologous to chromosome 18 of IPO323. We further conducted blast searches with the sequences of the transcribed regions on unitig 15 and received hits for genes on chromosomes 18, 20, and 21 of the reference IPO323/Zt09 indicating breakage of macrosynteny [8].

#### Percentage of mapped RNA-seq reads reflects infection stage-specific fungal biomass

For transcriptome datasets representing initial infection (stage A) and biotrophic growth (stage B), where comparably little fungal biomass is present and the wheat tissue is still fully intact, on average 8.02% and 8.2% of the filtered reads were aligned to the fungal genomes (Table 2, S5 Table). Exceptionally high alignment rates were obtained for isolate Zt05, (average stage A: 13.52%, average stage B: 12.41%), likely reflecting the strong proliferation on the leaf surface as well as the expanded biotrophic hyphal networks (Fig 3: Zt05, S3 Animation). For RNA-seq samples covering the lifestyle transition (stage C) and necrotrophic growth (stage D), where *Z. tritici* hyphal networks rapidly expand and the wheat mesophyll cells die, the amount of fungal-derived reads increased to 23.95% and 55% on average, respectively. The constant increase in fungal-derived RNA-seq reads during wheat infection reflects the increase in fungal biomass within the leaf tissue due to mesophyll colonization and lifestyle transition.

## Supporting information - S2 Text

### Supplementary Materials and Methods

#### Detection of H2O2 in *Z. tritici* infected wheat leaves

To visualize and localize the accumulation of the reactive oxygen species H_2_O_2_ within *Z. tritici* infected leaf tissue, we conducted 3,3’-diaminobenzidine (DAB) staining [1] at 4, 10, 14, 18, and 21 days post inoculation (dpi). Inoculated leaf parts were excised with a razorblade and immersed in DAB solution (1 mg/mL 3,3’-diaminobenzidine tetrahydrochloride (Thermo Fisher Scientific, Rockford, USA) in 0.05% [v/v] Tween 20). Samples were protected from light and DAB solution was infiltrated in two steps: 1^st^ at low pressure (600 mbar) for two times 15 min and 2^nd^ at gentle shaking (22 rpm) for 90 min. Subsequently, leaf samples were incubated overnight in de-staining solution (96% ethanol: acetic acid = 3:1 [v/v]) at gentle shaking (25 rpm). Cleared samples were stored in 96% ethanol and examined in 40% glycerol. Presence of H_2_O_2_ is indicated by reddish-brown precipitate in cleared leaf tissue. Infected leaf samples were documented by an iPhone 7 camera prior to the DAB staining and by a Canon EOS 600D post staining.

#### Staining of infected wheat leaves and confocal laser-scanning microscopy

Infected leaf parts were excised and de-stained in 96% ethanol. Samples were transferred to 10% KOH [w/v] at 85°C for 3 min to increase tissue permeability. For neutralization, leaf material was washed three times with 1X phosphate-buffered saline (PBS, pH 7.4) and subsequently incubated in a staining solution of 0.02% Tween 20 in 1X PBS (pH 7.4) with 10 μg/mL wheat germ agglutinin conjugated to fluorescein isothiocyanate (WGA-FITC) and 20 μg/mL propidium iodide (PI). Samples were protected from light and the staining solution was vacuum-infiltrated for 2 h where pressure was continuously reduced to 400 mbar for 5 min followed by ventilation of the desiccator and return to standard pressure. The staining solution was replaced by 1X PBS (pH 7.4) and the stained leaf samples were directly subjected to confocal microscopy analysis or stored lightproof at 4°C for later use. WGA was used to specifically label fungal hyphae [2] but was occasionally found to also bind to plant cell walls and bacteria. PI stains DNA and binds to plant and fungal cell walls [3]. FITC was excited with an argon laser at 488 nm and fluorescence was detected between 500 and 540 nm. A diode-pumped solid-state laser at 561 nm was employed for excitation of PI and emission was detected from 600 to 670 nm.

#### Generation of DNA plugs and karyotyping by pulsed-field gel electrophoresis

A non-protoplast protocol was used to produce DNA plugs for separating small chromosomes (~0.2 - 1.6 Mb) by pulsed-field gel electrophoresis (PFGE) [4]. Single cells of the three *Z. tritici* isolates were harvested from liquid YMS cultures. For preparation of plugs, 5 × 10^8^ cells were used as input and embedded in 1.1% low range agarose (Bio-Rad). Solidified agarose blocks were incubated in lysis buffer (1% SDS, 0.45 M EDTA, 1.5 mg/mL Proteinase K (Roth)) at 55°C for 48 h and subsequently washed three times in 1X TE buffer for 20 min. Plugs were directly submitted to pulsed-field gel electrophoresis or stored in 0.5 M EDTA at 4°C until further use.

PFGE was conducted using a contour-clamped homogeneous electric field (CHEF)-DR III apparatus (Bio-Rad) in 1% agarose in 0.5X TBE buffer applying the following conditions: temperature 14°C, 120° angle, 5 V/cm with a ramped 50 - 150 s switching interval for 48 to 68 h. Chromosomal DNA of *Saccharomyces cerevisiae* (Bio-Rad) was used as standard size marker. Gels were stained for 30 min in 1 μg/mL ethidium bromide solution and chromosome bands were detected with the Thyphoon Trio™ (GE).

## Supporting information - S3 Text

### Supplementary Information

#### Tools and *commands* used for genome analyses and processing and analyses of *Z. tritici* transcriptome data

##### # Quality control of RNA sequencing data

FastQC (http://www.bioinformatics.babraham.ac.uk/projects/fastqc/) version 0.11.2

##### # Removal of residual TruSeq adapter sequences

Trimmomatic [1] version 0.33

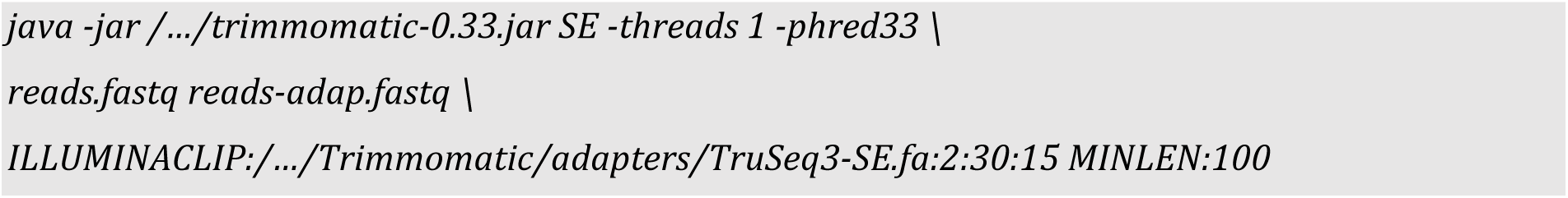

##### # Trimming of 12 nucleotides at 5' end of all reads

Trimmomatic [1] version 0.33

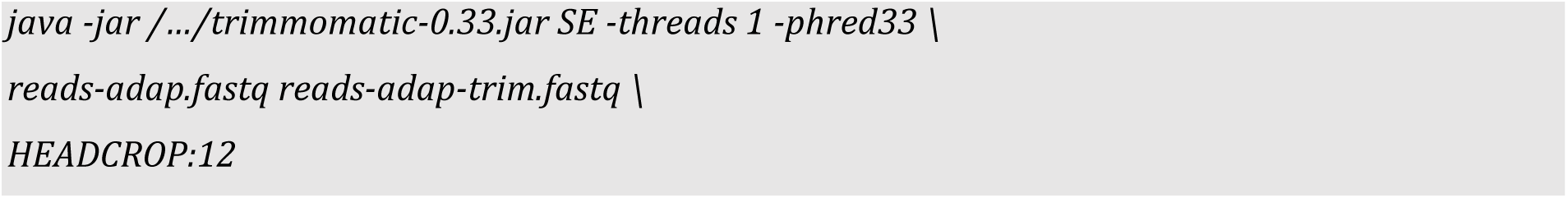

##### # Filtering of reads based on quality scores

**At least 80 % of bases must have a quality score ≥ 20 or read was dropped.**

FASTX-toolkit (http://hannonlab.cshl.edu/fastx_toolkit/) version 0.0.14

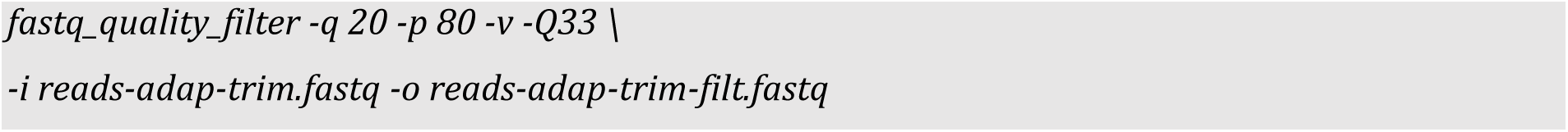

##### # Masking of low quality bases

**Nucleotides with quality score < 20 were masked with 'N'.**

FASTX-toolkit (http://hannonlab.cshl.edu/fastx_toolkit/) version 0.0.14

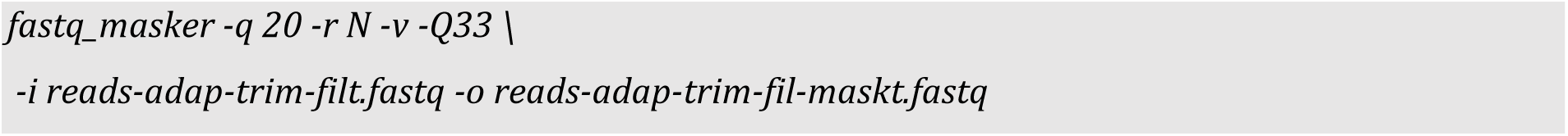

##### # Mapping of reads to genomes of *Z. tritici* isolates

TopHat2 [2,3] version 2.0.9

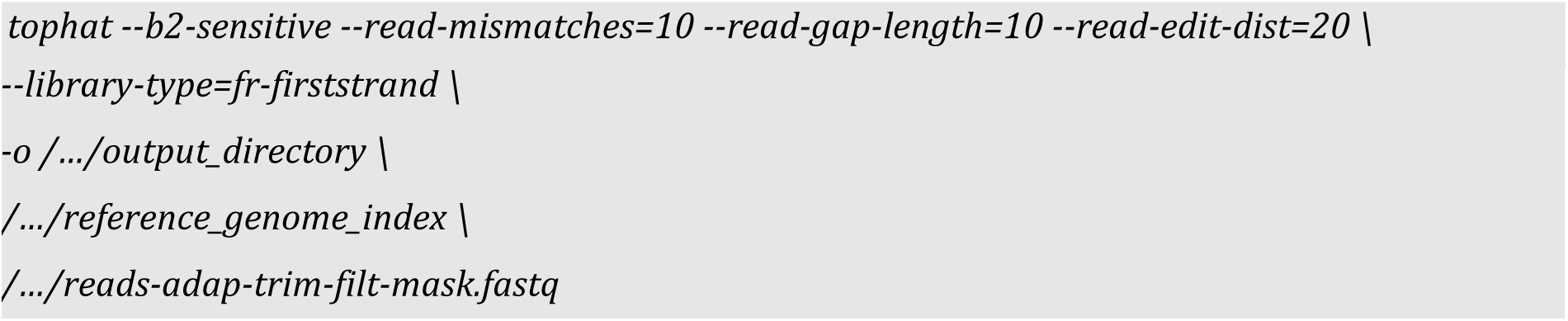

##### # Manipulation of RNA-seq read alignments

SAMtools [4] version 0.1.19

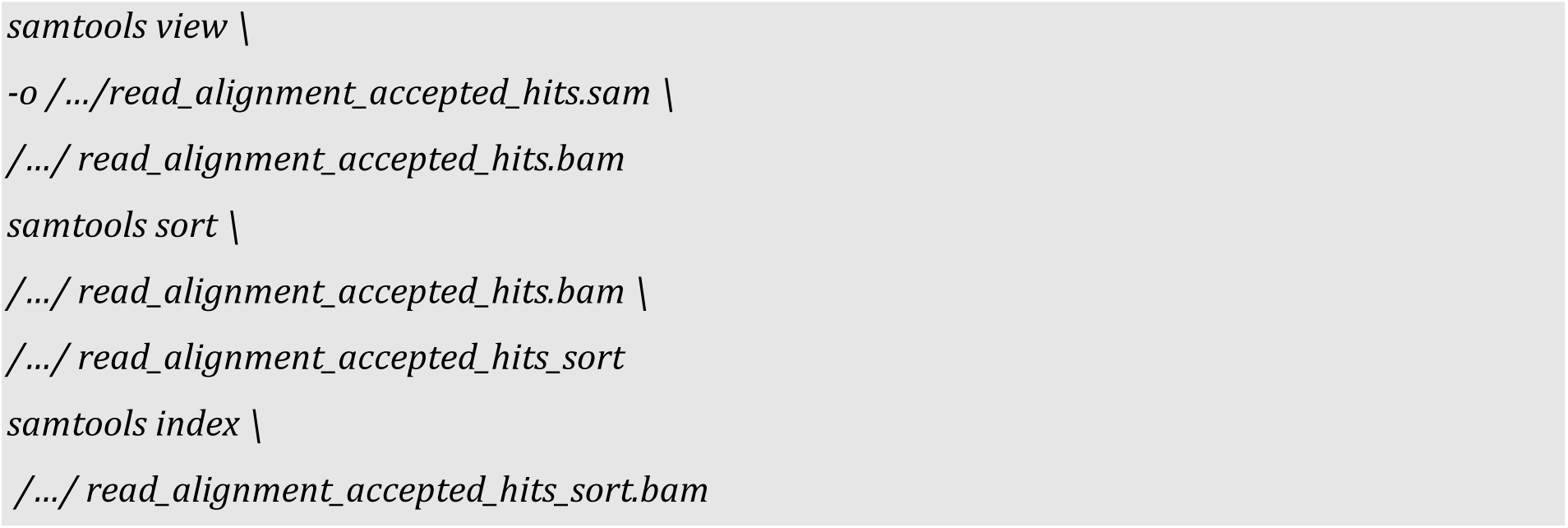

##### # Calculation of relative gene expression levels among the four infection stages within one *Z. tritici* isolate

Cuffdiff2 in Cufflinks [5] version 2.2.1

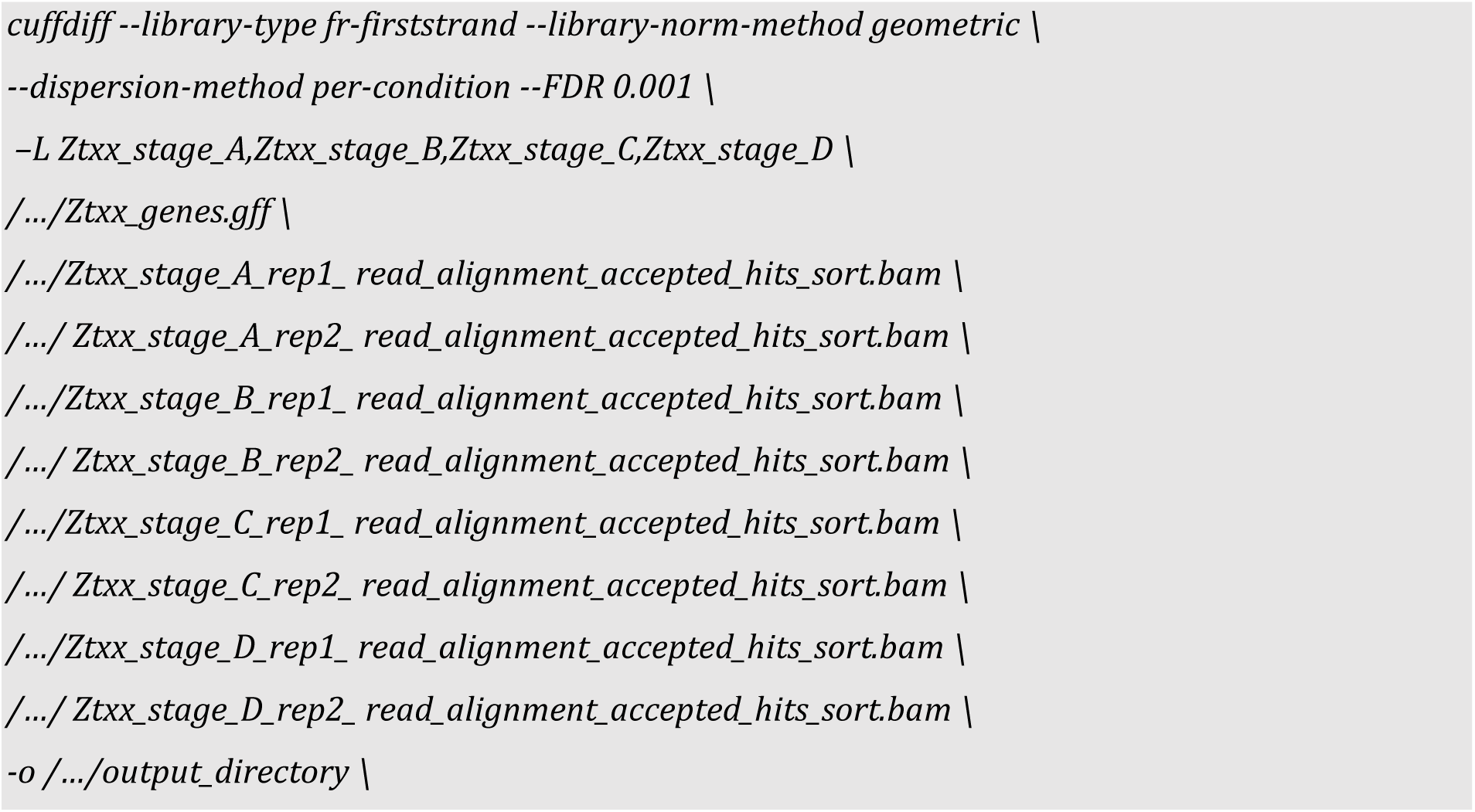

##### # Counting of mapped sequencing reads per gene

HTSeq [6] version 0.6.1p1

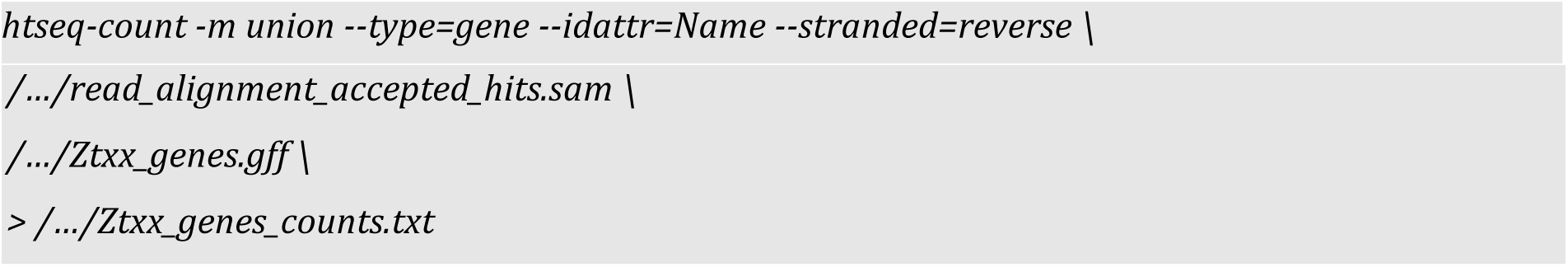

##### # Differential gene expression analyses

R package DESeq2 [7] version 1.10.1

\# Comparison between infection stages across all isolates

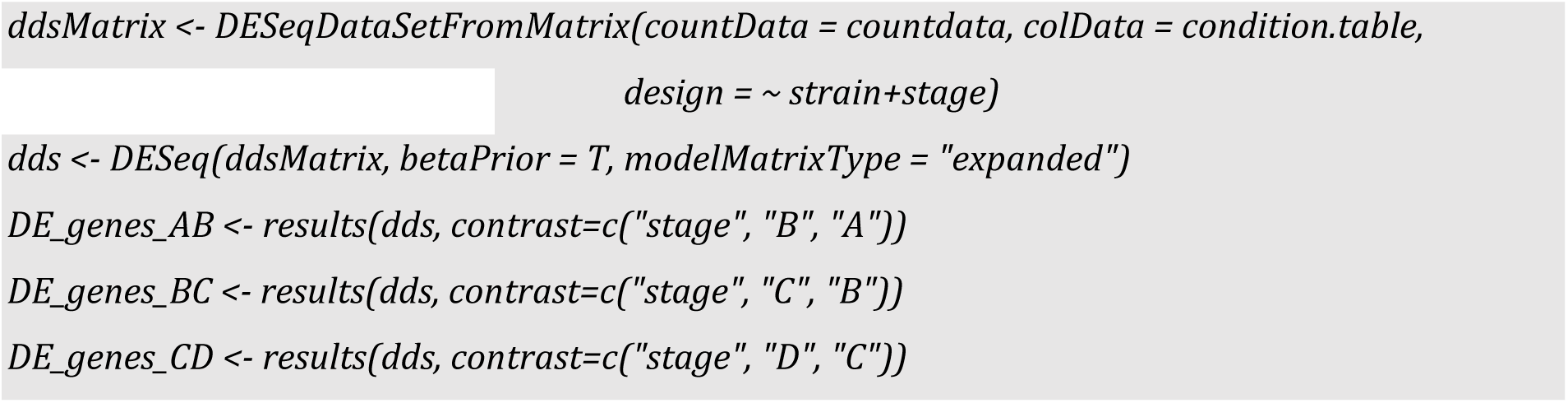

\# Comparison within infection stages between two isolates

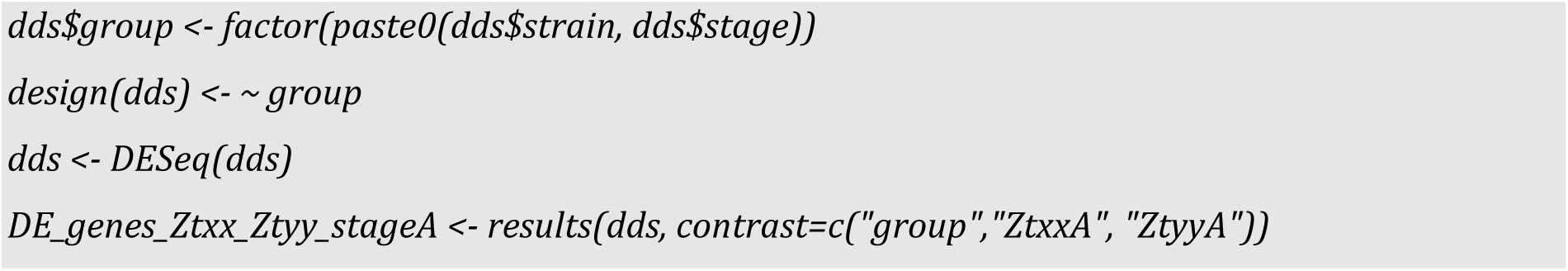

##### # Gene ontology (GO) term enrichment analyses

R package topGO [8] version 2.28.0

##### # Protein families (PFAM) enrichment analyses

Custom python script

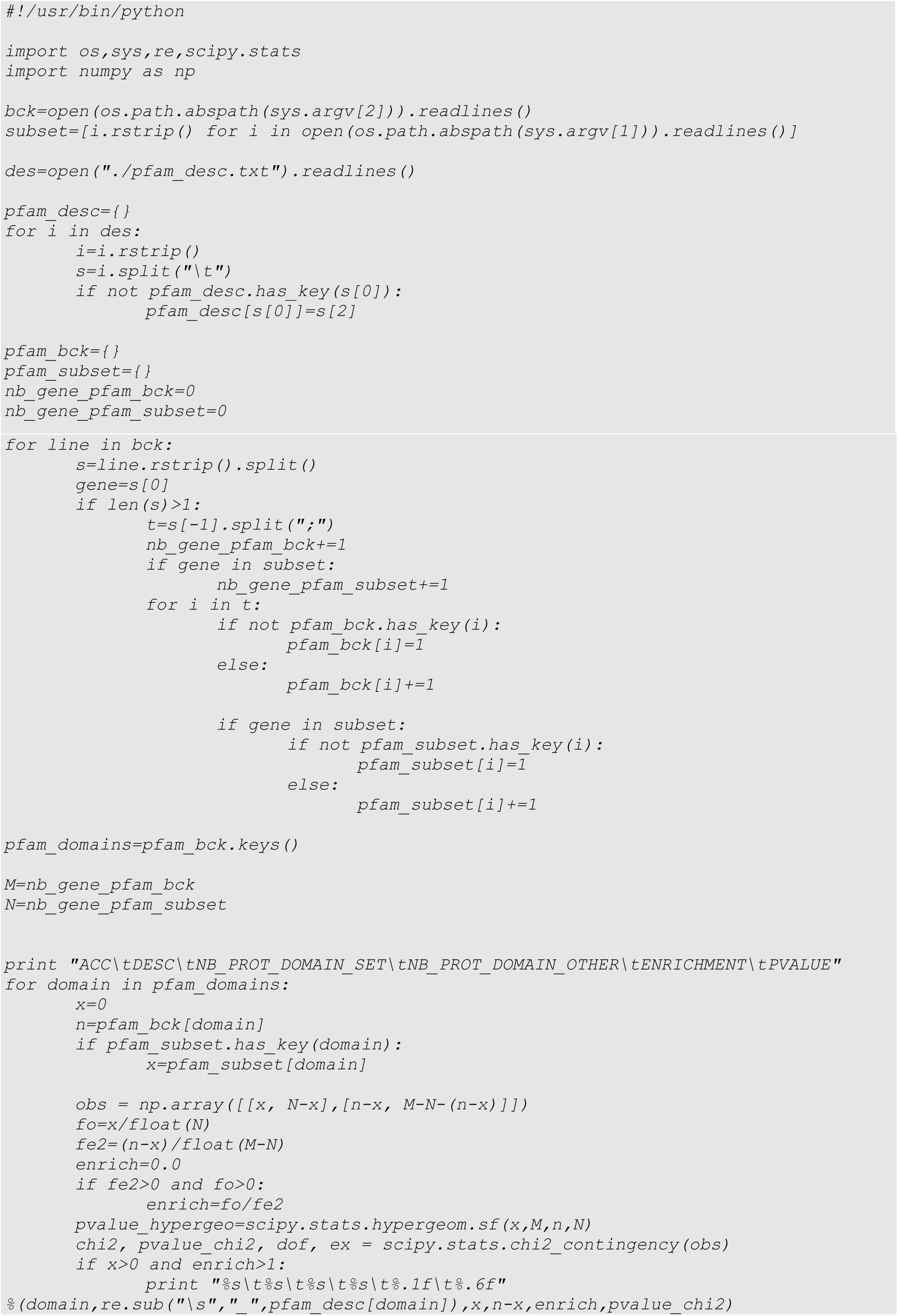

##### # Calculation of genomic distances between genes and TEs / H3K9me3 and H3K27me3

BEDtools [9] version 2.26.0

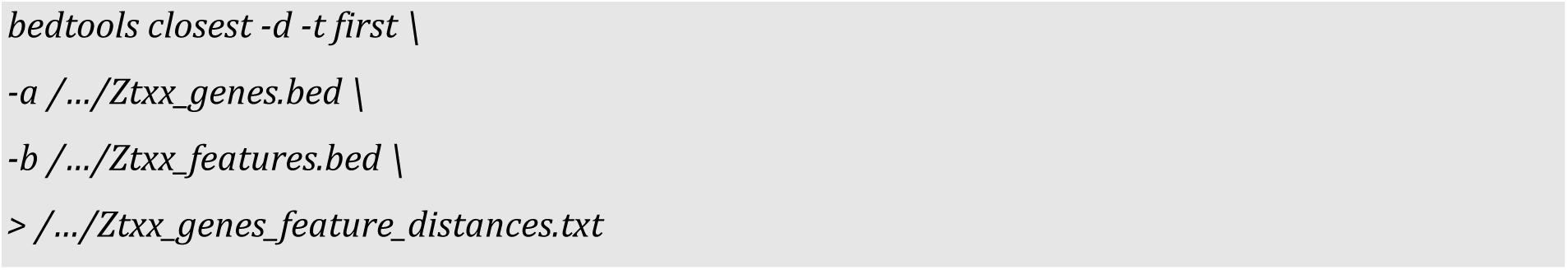

##### # *De novo* genome assemblies of Zt05 and Zt10 based on PacBio long reads

SMARTanalysis suite [10] version 2.3.0

HGAP version 3.0

Quiver

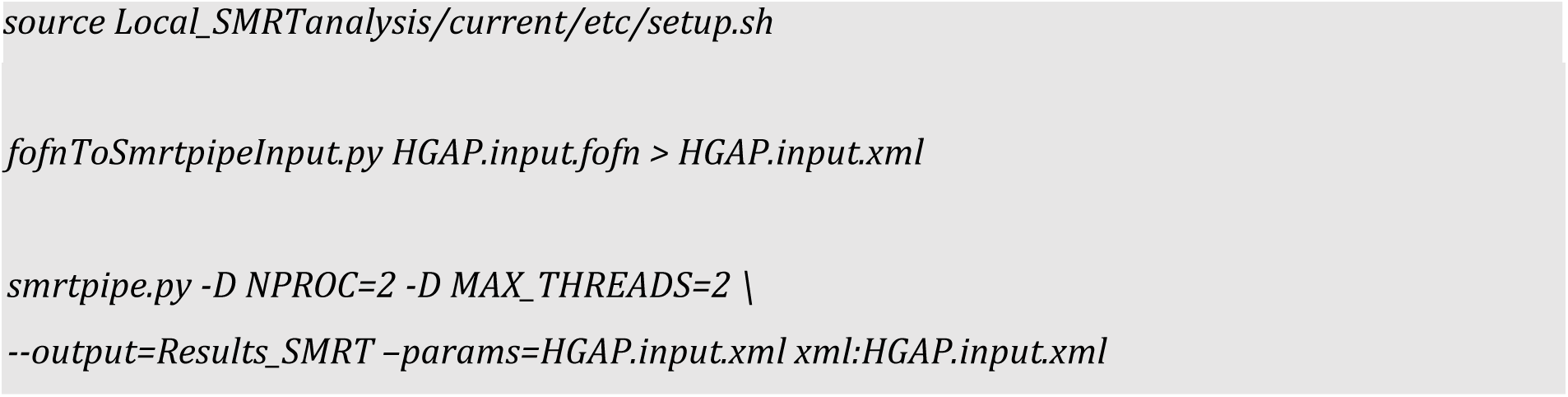

##### # Synteny mapping and analyses of IP0323/Zt09 chromosomes and Zt05 and Zt10 unitigs

SyMAP [11] version 4.2

Applying default settings and running NUCmer and PROmer to compute raw hits for anchor clustering.

Minimal contig size: Zt05: 1,000 kb

Zt09: 100,000 kb Zt10: 10,000 kb

Mugsy [12] version 1.r2.2

Generation of pairwise genome alignments of IPO323 - Zt05 and IPO323 - Zt10 applying default settings of Mugsy.

Custom python script to extract unique DNA blocks with a minimum length of 1 bp and calculate the total amount of unique DNA.

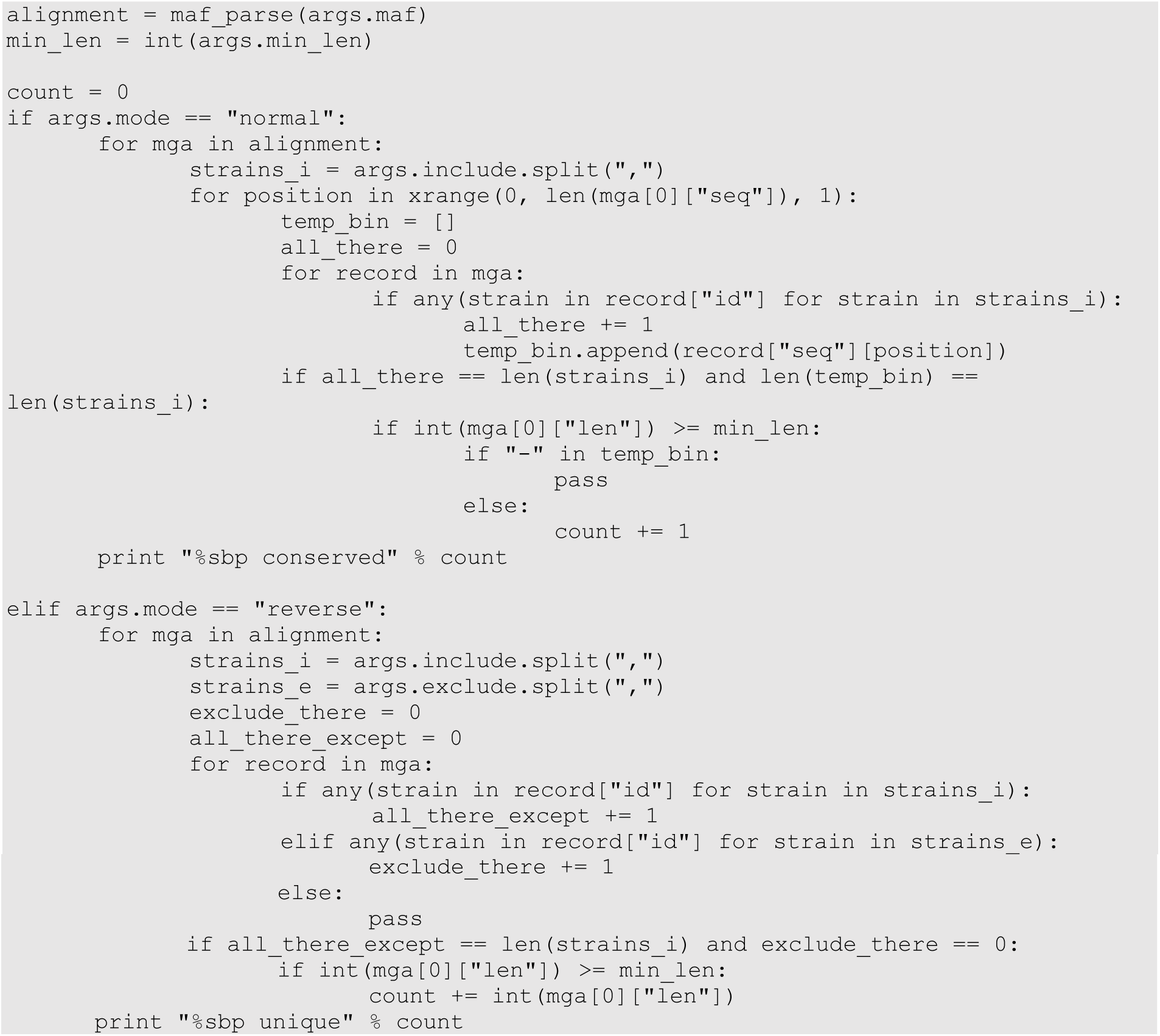

